# Transcriptomic profiling and machine learning uncover gene signatures of psoriasis endotypes and disease severity

**DOI:** 10.1101/2025.07.29.666780

**Authors:** Ashley Rider, Henry J. Grantham, Graham R. Smith, David S. Watson, John Casement, Simon J. Cockell, Jack Gisby, Amy C. Foulkes, Rafael Henkin, Wasim A. Iqbal, Tom Ewen, Shoba Amarnath, Sandra Ng, Paolo Zuliani, Nick Dand, Deborah Stocken, Christopher Traini, Elizabeth Thomas, Shanker Kalyana-Sundaram, Deepak K Rajpal, Kathleen M Smith, Jonathan N. Barker, Christopher E.M. Griffiths, Paola Di Meglio, Catherine H Smith, Richard B. Warren, Michael R. Barnes, Nick J. Reynolds, the PSORT consortium

## Abstract

**Background:** Despite increased understanding of psoriasis pathogenesis, molecular classification of clinical phenotypes and disease severity is poorly defined. Knowledge gaps include whether molecular endotypes of psoriasis underlie distinct clinical phenotypes and the positive and negative molecular regulators of disease severity across tissue compartments.

**Methods:** We performed comprehensive RNA-sequencing of skin and blood (n=718) from prospectively-recruited, deeply-phenotyped discovery and replication cohorts of 146 subjects with moderate-to-severe psoriasis initiating TNF-inhibitor (adalimumab) or IL-12/23-inhibitor (ustekinumab) therapy.

**Results:** Using two complementary methods for dimensionality reduction, we defined distinct but interconnected co-expression modules and factors within skin and blood that were significantly associated with disease phenotypes and disease severity, as measured by Psoriasis Area Severity Index (PASI). We identified a 14-gene signature negatively associated with BMI in nonlesional skin and disease severity in lesional skin, respectively. Genotype integration revealed that HLA-DQA1*01 and HLA-DRB1*15 genotypes were positively associated with baseline disease severity. Using Gaussian process regression followed by SHAP (SHapley Additive exPlanations), we defined two core drug independent and disease severity-associated gene modules in lesional skin - one positive, one negative - and a lesional 9-gene signature predictive of disease severity. Disease severity signatures in blood were only seen following adalimumab exposure, suggesting greater systemic impact of adalimumab compared to ustekinumab, in line with its side effect profile. In contrast, a gene signature in blood linked to HLA-C*06:02 status was independent of disease severity or drug.

**Conclusions:** These findings delineate gene-environmental and genetic effects on the psoriasis transcriptome linked to disease severity.

**Plain Language Summary:** Psoriasis is a common and debilitating skin disease, linked to multiple other inflammatory conditions. A lot is known about the mechanism of psoriasis and its inherited and external influences. Despite this, doctors cannot yet offer personalised treatments as it has been difficult to discover whether biological pathways are associated with disease severity, response to treatment or a person’s likelihood of having other linked diseases.

To help address this, we collected skin and blood samples and the personal characteristics of a group of people with severe psoriasis across the United Kingdom. Using computer-based methods, we discovered common biological processes underlying different psoriasis types, including genes that connect psoriasis severity with obesity, and another set of genes that help predict disease severity.

## Introduction

Psoriasis is a common, multifaceted immune-mediated inflammatory disease (IMID) characterised by symmetrical erythematous, hyperplastic and scaly plaques affecting the skin, with associated systemic inflammatory disorders including psoriatic arthritis, cardiovascular disease and metabolic syndrome which contribute to premature mortality (1). The aetiology and pathophysiology of psoriasis is complex and multifactorial (1). Over the last two decades significant progress has been made in understanding the pathophysiology of psoriasis and the contributions of genetic predisposition (2), environmental factors including infection, trauma and diet, acquired immune (e.g. T helper 1 (Th1), T17 cells; IL-17, IL-23 cytokines) and innate autoinflammatory factors (e.g. TNF, IL-36) which represent targets for highly effective biologic therapies. However, less progress has been made in translating these advances into individualised patient care. In part, this relates to significant knowledge gaps along the translational pathway that include: a) whether molecular endotypes within clinically homogeneous stable plaque psoriasis underlie distinct clinical phenotypes (e.g. sub-groups of subjects with specific comorbidities), b) the positive drivers and negative molecular regulators of disease severity across tissue compartments, and c) the relationship between molecular endotypes and clinical response to therapy including the side effect profiles of targeted therapies.

To address these questions, Psoriasis Stratification to Optimise Relevant Therapy (PSORT), an academic-industrial UK stratified medicine consortium (3), prospectively recruited formally-powered, deeply-phenotyped discovery and replication psoriasis patient cohorts, during the early phase of treatment with two distinct biologics, adalimumab (TNF inhibitor) and ustekinumab (IL-12/23 inhibitor) (Fig. 1a, b). Utilising this large multi-dimensional dataset, we aimed to identify gene networks that associated with specific disease endotypes, defined by clinical and phenotypic features (e.g. BMI) before commencing treatment (baseline); and disease severity endotypes, defined by Psoriasis Area and Severity Index (PASI)-associated gene expression profiles (Fig. 1c-d).

**Figure 1.**
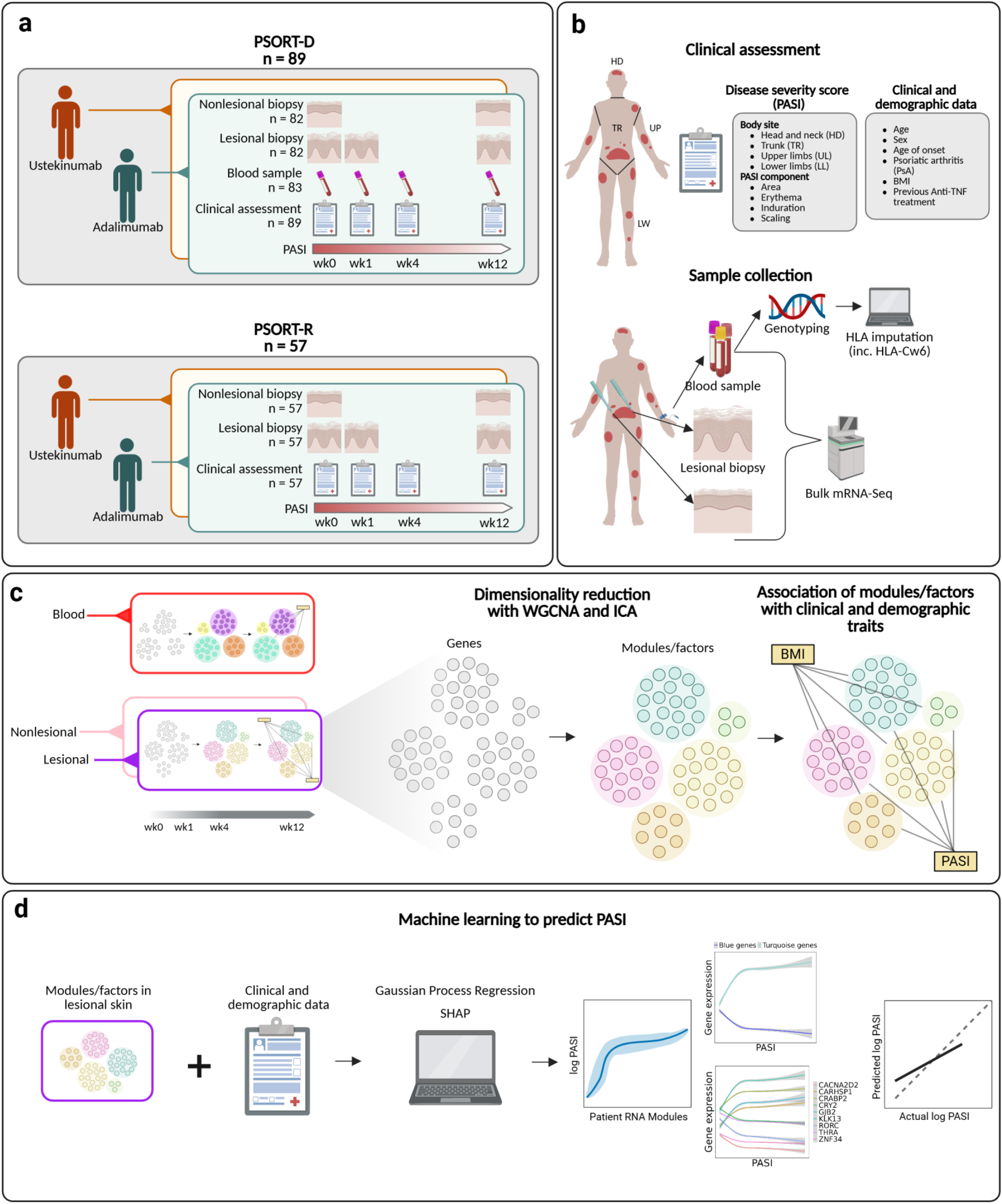
Summary of study design, including patient recruitment, sample collection and analysis methodology. **(a)** and **(b)** Participants with moderate-to-severe psoriasis eligible for biologic therapy were enrolled into discovery (PSORT-D, *n* = 89) and replication (PSORT-R, *n* = 57) cohorts and treated with either adalimumab or ustekinumab. At baseline and follow-up visits (weeks 0, 1, 4 and 12), lesional kin, nonlesional-skin and peripheral blood samples were collected, and Psoriasis Area and Severity Index (PASI) scores obtained. **(c)** To discover disease **endotypes** we applied two complementary dimension-reduction methods separately to skin and blood RNA-seq data. **Weighted Gene Co-expression Network Analysis (WGCNA)** – groups transcripts with similar expression into *modules* and represents each module by its first principal component (the **module eigengene**). **Independent Component Analysis (ICA)** – decomposes the dataset into statistically independent latent factors that capture underlying biological signals. Eigengenes and latent factors were correlated with clinical traits (e.g. BMI, HLA-C*06:02 genotype) and with PASI across time-points to reveal both baseline (disease) and disease-severity signatures. **(d)** Finally, we trained a **Gaussian Process Regression (GPR)** model – a flexible non-linear machine learning approach – using eigengenes and latent factors to predict log-transformed PASI. Model interpretability was provided by SHAP (SHapley Additive exPlanations), which quantifies the contribution of each module or factor for example to the prediction, allowing derivation of a concise gene signature of disease sever *Abbreviations:* PASI, Psoriasis Area and Severity Index; BMI, body-mass index; LS, lesional skin; NLS, non-lesional skin; WGCNA, weighted gene co-expression network analysis; ICA, independent component analysis; GPR, Gaussian Process Regression; SHAP, SHapley Additive exPlanations. Created in BioRender. Rider, A. (2025) https://BioRender.com/7qdytt1

As gene expression is controlled by multiple factors across different cell types, network analysis of bulk RNAseq data across a large number of clinical samples evaluates the coordination of gene expression by detecting underlying latent factors and co-expression modules. Moreover, these gene networks have been shown to more accurately reflect key biological processes and enable generation of signatures for disease classification and prediction of therapeutic outcomes, compared to analysis of individual genes (4). We therefore hypothesised that such network signatures would map to specific clinical phenotypes of psoriasis, including disease severity across time in response to biologic therapy. Consistent with this hypothesis, our integrative multi-omic analysis, across lesional (involved) and nonlesional (uninvolved) skin and whole blood identified gene signatures across distinct tissue compartments and cell types that classify psoriasis endotypes and associate with disease severity. This study yields important insights into pathogenic pathways in psoriatic skin and systemic pathways in blood, distinguishing disease endotypes linked to genetic and environmental factors and identifying reproducible biomarkers of disease severity.

## Materials and Methods

### Study design and patient cohort

We studied 146 subjects with moderate to severe plaque psoriasis commencing treatment with either adalimumab or ustekinumab; 89 subjects were assigned to the discovery cohort (PSORT-D) and 57 were assigned to the replication cohort (PSORT-R). Punch biopsies from involved and uninvolved skin were obtained at baseline and week 12; biopsies from involved skin were also derived at week 1. Blood samples were obtained at baseline, week 1, week 4 and week 12.

### RNA sequencing and genotyping

Following RNA extraction and quality control, the skin and blood samples underwent RNA sequencing using the Illumina HiSeq 3000 (PSORT-D skin); Illumina HiSeq 4000 (PSORT-D blood) and Illumina NovaSeq 6000 (PSORT-R skin). DNA was isolated from the blood samples and genotyping was performed with Illumina HumanOmniExpressExome-8 v1.2 and v1.3 BeadChips. *HLA-C*06:02* genotype was imputed using SNP2HLA (version 1.0.3) based on the Type 1 Diabetes Genetics Consortium reference panel (5).

### Pseudo-alignment

Following RNA sequencing, reads were pseudo-aligned using Kallisto (6) and transcript-level counts were aggregated to gene-level.

### WGCNA

Identification of co-expressed gene modules with WGCNA was carried out separately in the skin and blood samples from the discovery cohort. Within skin and blood, all samples were used for module identification, i.e. both lesional and nonlesional samples (within skin), both drug cohorts and all time points. Following low count filtering and normalisation, all samples for a given tissue compartment (i.e. skin or blood) underwent WGCNA using the *blockwiseModules* function from the WGCNA R package (62, 63) with a minimum module size of 30, a merge cut height of 0.1, and a soft thresholding power of 12 for skin and 5 for blood. The *moduleEigengenes* function was then used to calculate module eigengenes for the resulting gene modules in the discovery and replication cohorts (for skin modules). The *modulePreservation* function was used to derive module preservation statistics for the skin modules in blood and vice versa. The right-tailed Fisher’s exact test was used to test the significance of gene overlap between the skin and blood modules.

### ICA

Independent component analysis (ICA) was carried out separately in the skin and blood samples from the discovery cohort using the *imax* method from the ica R package (9). Within skin and blood, all samples were used for ICA, i.e. both lesional and nonlesional samples (within skin), both drug cohorts and all time points. Factor metagenes, analogous to module eigengenes from WGCNA, were then derived for these factors in the discovery and replication cohorts (for skin factors). The ReducedExperiment package was used to perform enrichment tests and associating testing (10).

### Trait associations

The module eigengenes and factor metagenes were correlated with clinical and demographic factors using Pearson correlation. Significant associations were defined by an adjusted p-value threshold (FDR) of 0.05 in the discovery cohort and nominal p-value threshold of 0.05 in the replication cohort. Significant correlations were also required to be of the same sign in the discovery and replication cohorts.

### Deconvolution

Cell type deconvolution of the bulk expression data was done using Cibersort X (11). This was done for the skin samples using a reference matrix generated from single-cell RNA-Sequencing data from 38,274 skin cells across 5 inflammatory skin conditions, including psoriasis (12). In blood, deconvolution was done using the LM22 signature matrix provided by Cibersort X.

### Differential expression analysis

We implemented two differential expression models using the limma-voom pipeline (66, 67) in R: (i) a disease severity model to identify genes associated with PASI in each tissue (lesional skin, non-lesional skin, and blood), irrespective of time point, and (ii) a disease phenotype model to identify genes in each time point associated with clinical variables of interest (e.g. BMI) at baseline.

### Pathway analysis

Pathway enrichment analysis of the WGCNA module genes, latent factor genes, and DEGs was done using Metascape (15). Additional enrichment analysis of DEGs was done with Ingenuity Pathway Analysis (IPA) (16).

### Machine learning

Gaussian process regression, a family of Bayesian models, was chosen to model the relationship between several features (i.e. Demographics + clinical features, Skin factors and Skin RNA modules) and Psoriasis Area Severity Index (PASI). GPRs can model numerous relationships in a non-linear manner with a limited number of examples. The GPR model parameters were optimised by maximising the marginal likelihood during cross-validation. Specifically, performance and parameter selection were evaluated using repeated 10-fold cross-validation at the subject level. Predictions from the trained models were then assessed using shuffled 10-fold held-out testing subjects. Performance metrics such as the coefficient of determination (R^2^) and mean absolute error (MAE) metrics were calculated for each shuffled dataset. We compared the performance of the GPR model to a linear ridge regression model. Through comparison to this baseline method, we were able to determine whether the non-linearity captured by the GPR models improved the predictive accuracy (Supplementary Tables S7-9). Feature importance was assessed using a model agnostic approach from the SHAP (SHapley Additive exPlanations) python package. Additional additive GPR models were trained using the top 10 aligned genes from each module identified from a correlation analysis with PASI. SHAP method was also applied to understand the contribution of the 10 gene modules driving PASI prediction.

### Statistical information

Module identification with WGCNA was done using pearson correlation with pairwise complete observations. Similarly, trait correlations with WGCNA module eigengenes and ICA latent factors were calculated using pearson correlation with pairwise complete observations. The trait correlations were calculated in lesional skin, non-lesional skin and blood separately and with different sample subsets for each variable type: correlations with the disease endotype variables (i.e. age of onset, onset type, anti-TNF naive status, PsA, sex, age, BMI, Cw6 status) were calculated using the baseline samples from both drug cohorts and correlations with PASI were calculated for each drug cohort separately using samples from all time points. To account for multiple testing within each tissue, the p.adjust function with method “fdr” in R was applied to the trait correlation p-values for modules and factors separately. Additionally, correlations of module eigengenes and latent factors with per-sample imputed absolute cell fractions were calculated using Kendall’s tau coefficient and FDR-adjusted p-values were derived as above. P-values from gene-level differential expression modelling of disease endotypes and disease severity were adjusted using Storey’s method (17). Further details about statistical methodology are available under the relevant subsections in the main and supplementary materials and methods sections.

## Results

### Study Design

Our study design was previously reported (3,18). The timing of sample collection and details of sample numbers are illustrated in Figure 1 (see main and supplementary Materials and Methods). We studied 89 subjects with stable plaque psoriasis initiating biologic therapy, 82 of whom provided skin biopsies (41 with adalimumab and 41 with ustekinumab; 400 total samples) and 83 of which provided blood samples (40 with adalimumab and 43 with ustekinumab; 318 total samples) (discovery cohort) (supplementary Materials and Methods). We replicated findings in a further cohort of 57 subjects who provided skin samples (29 with adalimumab, 28 with ustekinumab; 276 total samples). Power calculations based on Guo et al (19) and a pilot study (20) indicated a discovery cohort sample size requirement of 40 (Supplementary Methods and Supplementary Fig. 1). Subject characteristics for included participants are shown in Supplementary Tables 1a and 1b.

### Identification of gene expression signatures in skin and blood

To define the relationship between transcriptional signatures and clinical phenotypes, we used two complementary methods for dimensionality reduction, Weighted Gene Correlation Network Analysis (WGCNA) and Independent Component Analysis (ICA) (Fig. 2a, b, main and supplementary materials and methods) (4). In brief, WGCNA groups genes into modules based on co-expression and ICA identifies latent variables that describe patterns of expression variation in the data (4). WGCNA-derived eigengenes summarise the main trend in co-expressed gene modules, while ICA-derived metagenes emphasise independent patterns of gene expression (4). To facilitate comparisons between lesional and non-lesional skin and across time, all skin samples from the adalimumab and ustekinumab drug cohorts at weeks 0, 1 and 12 were analysed together. However, there was extensive transcriptomic variation between skin and blood; therefore, these tissues were analysed separately.

**Figure 2.**
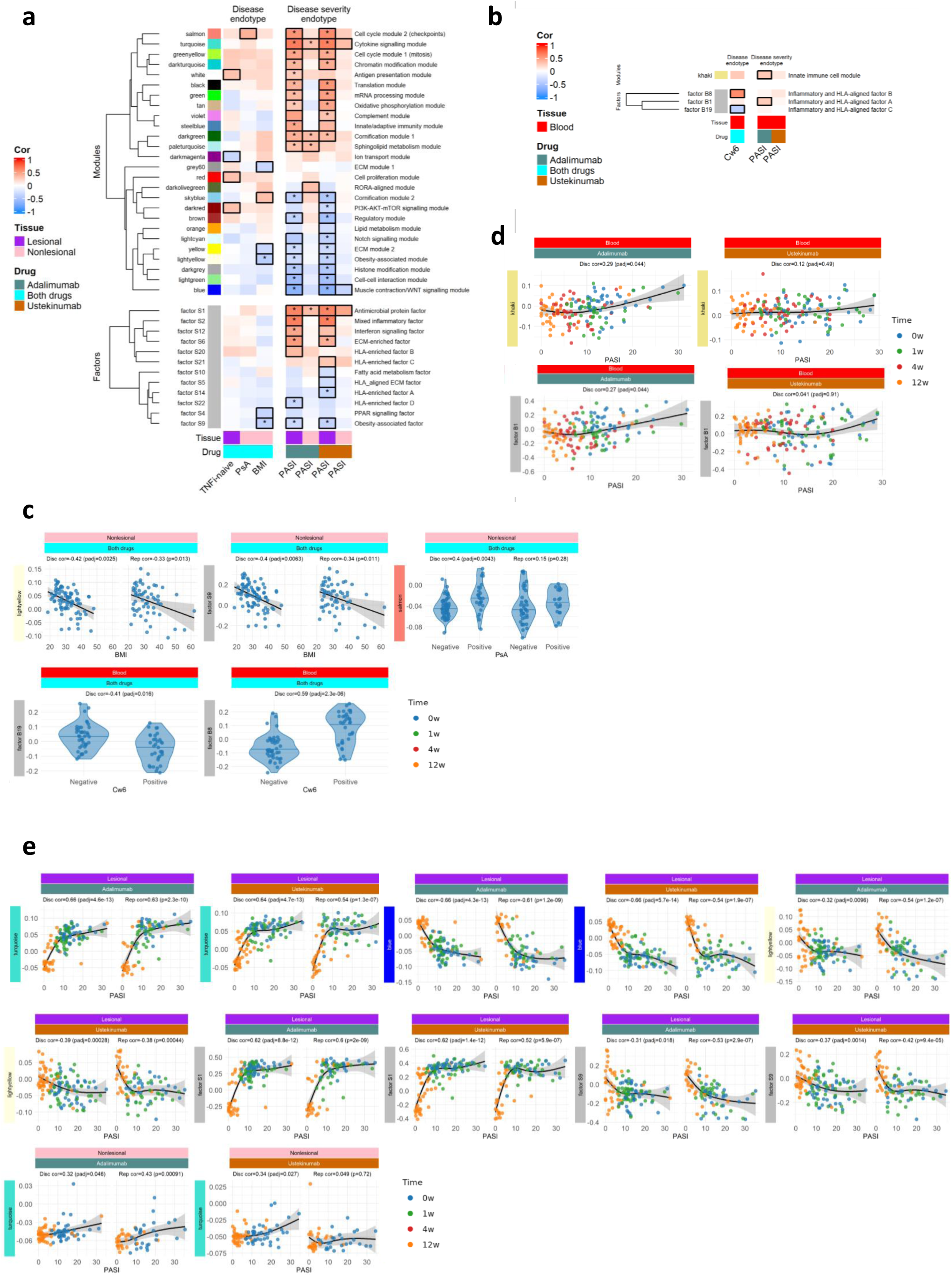
Systems-level gene modules and latent factors correlate with disease and disease severity endotypes in psoriasis. Heatmaps display pearson correlations between (upper panels) **WGCNA module** eigengenes or (lower panels) **ICA latent factors** and clinical/demographic variables for **(a)** skin and **(b)** blood in the discovery cohort. Black outlines denote correlations significant at false-discovery rate (FDR) ≤ 0.05; asterisks indicate the correlation replicates (same sign, *P* < 0.05) in the independent replication cohort. Only modules/factors with at least one replicated correlation are shown; descriptors summarise the dominant biological theme of each module/factor. Panels **(c–e)** illustrate three examples: **Baseline clinical endotypes** – correlations with BMI in skin and *HLA-C*06:02* genotype in blood **(c)**. **Disease-severity endotypes (skin)** – non-linear relationships between eigengenes and PASI across all visits **(e)**. **Disease-severity endotypes (blood)** – analogous plots for blood eigengenes **(d)**. Points are coloured by visit (blue = week 0, green = week 1, red = week 4, orange = week 12). Violin plots include a median line. Curves represent natural-spline fits (3 d.f.) with 95 % confidence bands. *Abbreviations:* as in Fig. 1 plus HLA, human-leukocyte antigen; FDR, false-discovery rate.

WGCNA identified 34 co-expressed gene modules in lesional skin and nonlesional skin across all time points (Supplementary Fig. 2a, Supplementary Table 2). Individual modules contained between 34 and 2677 genes (Supplementary Table 2). ICA identified 24 factors in lesional skin and nonlesional skin across all timepoints (Supplementary Table 3). WGCNA of the blood RNA-Seq data identified 26 co-expressed gene modules (Supplementary Fig. 2b) with individual modules containing between 50 and 1333 genes (Supplementary Table 4). ICA identified 21 factors in blood (Supplementary Table 5). Many of the factor metagenes were highly correlated with module eigengenes in both tissues (Supplementary Fig. 3a-b), indicating that the methods converged on similar key signatures, providing cross-validation.

To define disease and severity endotypes, module eigengenes and factor metagenes were correlated with clinical phenotypes and *HLA Cw6* genotype status at baseline, designated as disease endotype; and with PASI across all time points, termed disease severity endotype. These associations were replicated by independent testing within the replication cohort (Fig. 2a, 2b).

To further define the functional relevance of the co-expressed WGCNA modules and ICA factors, systems analysis was performed; the top pathway enrichments and exemplar aligned genes are described in supplementary tables 2-5 and in the results below.

We carried out module preservation analysis to assess the degree to which skin modules were preserved in blood and vice versa. The skin modules black (translation) and tan (oxidative phosphorylation) exhibited strong evidence for preservation in blood and were found to have the most significant gene overlap with the gold (translation) module in blood (Supplementary Fig. 4a-b). The green module (mRNA processing) was also strongly preserved in blood and exhibited significant gene overlap with the olivedrab (oxidative phosphorylation) module in blood (Supplementary Fig. 4a-b).

### Disease Endotype

Significant associations of clinical phenotypes with WGCNA modules and ICA factors were found in both lesional skin and nonlesional skin (Fig. 2a) at baseline. Notably, we observed that the lightyellow module (insulin and hormone secretion) and factor S9 (obesity-associated) each displayed significant and replicable negative associations between their eigengene expression and i) BMI in nonlesional skin and ii) PASI in lesional skin (Fig. 2a, 2c; Supplementary Fig. 5).

This supports our earlier observed association between high BMI and poor clinical response in a larger cohort of patients (21). Functional enrichments for lightyellow and factor S9 included: secretory pathways, hormone signalling, transport of small molecules and Wnt signalling (Supplementary Tables 2 and 3). Twenty-one genes within the intersection of lightyellow and factor S9 were independently negatively associated with BMI (Fig. 3a-b). Notably 14 of these genes, including *SCGB1D2* (Secretoglobin Family 1D Member 2)*, MMP7*, *DNER* (Delta/Notch Like EGF Repeat, regulates adipogenesis) and *PDE9A* (stimulates lipogenesis) were negatively associated with PASI in lesional skin across all time points (Fig. 3c). Deconvolution of bulk RNAseq data with single cell RNAseq data from 38,274 skin cells (12) revealed that lightyellow, factor S9, and the 14 gene core BMI/PASI signature (including *SCGB1D2, DNER* and *PDE9A*) were highly enriched with genes expressed within the pilosebaceous unit (Supplementary Figs. 6, 7b and 8).

**Figure 3.**
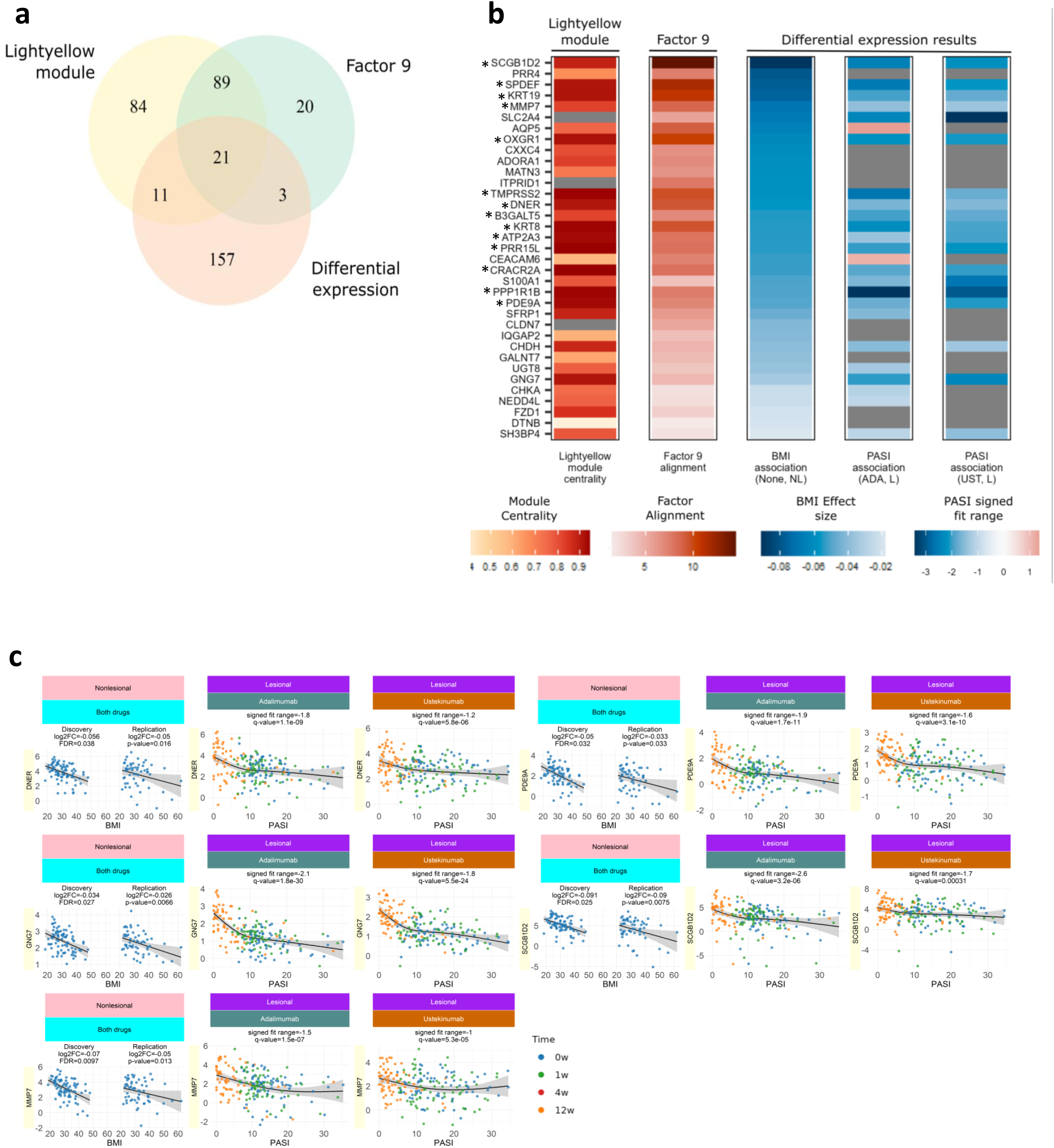
A BMI-related transcriptomic signature in non-lesional skin is also linked to disease severity in lesional skin. **(a)** Venn diagram showing overlap between genes that (i) belong to the **lightyellow WGCNA module**, **(ii)** have high alignment to **ICA factor 9** (absolute scaled loading > 5), and (iii) are differentially expressed with BMI in nonlesional skin (FDR < 0.05, *limma*). **(b)** Heatmap of key genes within this shared signature. Columns correspond to genes; rows encode three attributes: membership of the light-yellow module (yellow), strength of loading on factor 9 (red intensity), and significance of BMI association (blue intensity). Genes marked with * are also significantly associated with PASI in lesional skin (q < 0.05) in both biologic cohorts. **(c)** Representative genes are plotted: expression (log-CPM) versus BMI in non-lesional skin (left) and versus PASI in lesional skin (right); blue regression lines with 95 % CI illustrate consistent direction of effect. *Abbreviations:* BMI, body-mass index; CPM, counts per million; ADA, adalimumab; UST, ustekinumab; LS, lesional skin; NLS, non-lesional skin.

Two blood factors were significantly associated with *HLA-C*06:02* genotype (as a binary trait), with factor B8 (inflammatory and HLA-aligned B) being positively correlated and factor B19 (inflammatory and HLA-aligned C) negatively correlated (Fig. 2b). These factors were enriched for antigen processing, graft versus host disease and allograft rejection, with these enrichments being driven predominantly by *HLA*-genes (Supplementary Table 5). The factors were highly aligned with the expression of genes in the 6p21 gene region, including *PPP1R18*, *HLA-DRB1*, *HSPA1L*, *GPSM3*, *HSPA1A*, *LY6G5B*, *CSNK2B* and *TUBB* (factor B8); and *HLA-DRA*, *HLA-DQB1*, *AGPAT1*, *PSMB9*, *HLA-DQA1*, *HLA-DRA* and *HLA-DMB* (factor B19).

### Disease severity endotype

Few previous studies have studied the relationship between gene expression patterns and disease severity across different tissues in psoriasis (22,23). We therefore investigated whether gene expression within lesional skin, nonlesional skin and blood was associated with whole body disease severity scores at the time of sampling. PASI is the gold-standard disease severity measure and represents the average redness, thickness, and scaliness of psoriasis lesions weighted by the area of involvement (Fig. 1b, top panel) (24).

#### Lesional skin

Remarkably, 21 out of 34 WGCNA modules in lesional skin showed highly significant and reproducible correlations with disease severity in at least one of the drug cohorts (Fig. 2a); 12 showing positive correlations and 9 negative correlations with disease severity. Cross-correlation of the key modules and factors identified two distinct closely clustered blocks (i and ii) (Supplementary Fig. 3a). Strikingly, the first block (i) comprised modules and factors (e.g. turquoise module and factor S1; cytokine and anti-microbial peptide signalling) positively associated with disease severity whereas the second block (e.g. yellow module and factor S9; ECM and Insulin and hormone secretion signalling) (ii) was negatively associated with disease severity (Fig. 2a; Supplementary Fig. 3a). Exemplar scatter plots are shown in Fig. 2c and Supplementary Fig. 9.

To gain further insight, we deconvoluted the bulk RNAseq data using single cell RNA-seq data (12). Cluster analysis showed the presence of two main blocks that were distributed according to positive or negative disease severity associations (Supplementary Fig. 6). For example, we observed resolution of keratinocyte subsets into two distinct clusters, consistent with previous single cell (25) and spatial transcriptomic studies (26). Thus, keratinocyte-1 (KRT1^+^, KRT10^+^ S100A8/9^+^ spinous), keratinocyte-4 (KRT1^+^, KRT10^+^ S100A8/9^+^ spinous), keratinocyte-6 (KRT5^+^, KRT14^+^ KRT1^+^, KRT10^+^ PCNA^+^ supra-basal) and keratinocyte-7 (KRT5^+^, KRT14^+^, COL17A1^+^ basal) subsets (Supplementary Fig. 7a) correlated with positively-associated disease severity modules and factors (e.g. turquoise module, darkgreen module and factor S1) whereas keratinocyte-3 (KRT5^+^, KRT14^+^, KRT1^+^, KRT10^+^ supra-basal), keratinocyte-5 (KRT1^+^, KRT10^+^ IVL^+^ supra-spinous) and keratinocyte-8 (KRT5^+^, KRT14^+,^ COL17A1^+^, PCNA^+^ basal) (25) subsets (Supplementary Fig. 7a) aligned with negatively-associated disease severity modules and factors (e.g. yellow module, skyblue module and factor S9) (Supplementary Fig. 6). Notably, each cluster comprised basal, suprabasal and spinous subsets, reinforcing a mechanistic model of a switch in keratinocyte differentiation program and phenotype with disease severity progression (12,25). Additionally, and in line with the current pathophysiological understanding of the role of innate and acquired immunity in psoriasis (25,27), positively-associated disease severity modules and factors showed strong associations with myeloid-1 (dendritic cells/macrophages), T cell-2 (Th1, Th17), T cell-3 (cytotoxic T lymphocytes) and venule-2 cells (which regulate the trafficking of immune cells into tissues) (Supplementary Figs. 6, 7e, 7f and 7g). Deconvolution of negatively-associated disease severity modules and factors revealed enrichment for fibroblast-5 (SFRP+, MFAP5+; f2/3 stromal/mesenchymal) (28) and Langerhans cells (Supplementary Figs. 6, 7b, 7c, and 7d).

#### Nonlesional skin

Although clinically resembling normal skin, it is recognised from gene and protein expression studies that nonlesional skin represents a pre-psoriatic state that is primed to develop into psoriasis (29). We extend these findings to show that in nonlesional skin, eigengene expression of three WGCNA modules (turquoise, darkgreen (cornification 1) and paleturquoise (sphingolipid metabolism) and factor S1 significantly and reproducibly positively correlated with whole body disease severity in the adalimumab group (Fig. 2a, 2c). Of note, there were no significant negatively-associated disease severity modules or factors in nonlesional skin.

#### Blood

Limited previous studies have systematically investigated mRNA biomarkers of psoriasis disease severity in blood (30). The khaki (innate immune cell; neutrophil degranulation) module and factor B1 (inflammatory and HLA aligned A; phagosome) were positively associated with disease severity in the adalimumab cohort, and deconvolution indicated enhanced representation of neutrophils (Fig. 2b, Supplementary Fig. 10, Supplementary Tables 4 and 5). Strikingly, in the ustekinumab cohort there were no correlations reaching statistical significance (Fig. 2b, c).

### Predicting disease severity using machine learning

To predict disease severity from modules and factors, we employed additive *Gaussian*-process regression (GPR; Supplementary material and methods). GPR was well-suited to our “small-n, large-p” design, which comprised 718 samples from 146 subjects and approximately 20,000 transcripts within 60 modules and 45 factors across skin and blood, as this method combines (i) non-linear flexibility (through radial-basis and Matérn kernels) with (ii) a Bayesian framework that returns per-sample credible intervals, an essential read-out for clinical risk-stratification. Moreover, its additive kernel decomposition facilitated transparent attribution of individual module and factor contributions via SHAP values (Fig. 4b, d), which cannot be obtained as cleanly from tree-based ensembles or neural networks.

**Figure 4.**
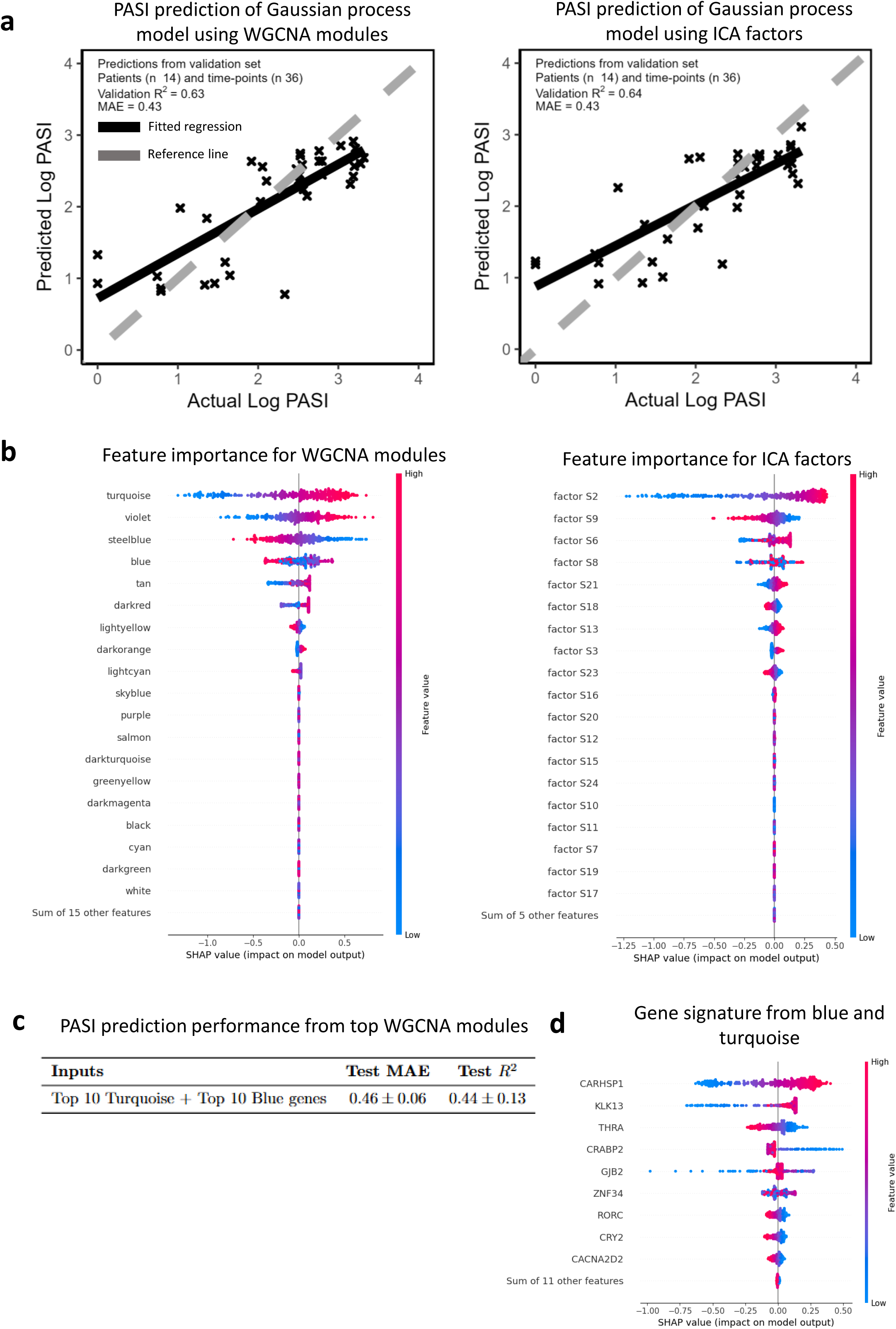
Gaussian Process Regression accurately predicts log PASI from transcriptomic modules and factors. Data from 139 individuals (discovery + replication) were randomly partitioned into training (80 %, *n* = 111), validation (10 %, *n* = 14) and test (10 %, *n* = 14) sets. **(a)** Predicted versus observed log PASI for the optimal models built from **WGCNA eigengenes** (left) or **ICA latent factors** (right) on the validation set. **(b)** SHAP summary plots rank feature importance. Positive SHAP values (right of 0) indicate features pushing predictions **up** (higher PASI); negative values (left) push predictions **down**. **(c)** Removing all but the top eigengenes/factors shows that a parsimonious 10-gene signature (derived from blue + turquoise modules) preserves predictive performance. **(d)** Applying SHAP to the top performing modules (blue and turquoise) identified a 9 gene signature predictive of log PASI. Comparison with a baseline linear ridge-regression (see Supplementary Fig. 11) highlights the gain from modelling non-linear interactions. *Abbreviations:* as in Fig. 1; SHAP, SHapley Additive exPlanations.

A linear ridge-regression (RR) baseline and an extensively tuned XGBoost model were trained for comparison. In five-fold cross-validation GPR achieved MAE = 0.45 ± 0.07 and R² = 0.53 ± 0.09, matching XGBoost while requiring only three hyper-parameters and outperforming RR (ΔMAE = +0.06; ΔR² = –0.08).

The final GPR and RR models, fitted to the combined Discovery + Replication cohorts, tracked the training data well (R² = 0.59–0.78; Table 1, Supplementary Table 7). Crucially, only GPR supplies calibrated uncertainty bands around each prediction, complementing global metrics such as MAE and R².

**Table 1.** PASI prediction performance of additive Gaussian process models. Shown are means ± SD for 20 randomly shuffled 10-fold testing datasets.

| Inputs | Test MAE | Test $R^2$ |
| --- | --- | --- |
| Demographics, clinical features, WGCNA modules, and ICA factors | $0.47 \pm 0.07$ | $0.49 \pm 0.10$ |
| All WGCNA modules | $0.45 \pm 0.07$ | $0.53 \pm 0.09$ |
| Turquoise module | $0.45 \pm 0.07$ | $0.52 \pm 0.10$ |
| Violet module | $0.61 \pm 0.09$ | $0.12 \pm 0.13$ |
| Steelblue module | $0.61 \pm 0.10$ | $0.14 \pm 0.15$ |
| Blue module | $0.45 \pm 0.07$ | $0.54 \pm 0.09$ |
| Tan module | $0.55 \pm 0.06$ | $0.29 \pm 0.09$ |
| All ICA factors | $0.46 \pm 0.06$ | $0.51 \pm 0.09$ |
| Factor S1 | $0.45 \pm 0.06$ | $0.51 \pm 0.09$ |
| Factor S2 | $0.46 \pm 0.08$ | $0.51 \pm 0.11$ |
| Factor S6 | $0.52 \pm 0.07$ | $0.35 \pm 0.14$ |
| Factor S9 | $0.58 \pm 0.09$ | $0.21 \pm 0.10$ |
| Factor S8 | $0.66 \pm 0.09$ | $0.00 \pm 0.04$ |
| Factor S21 | $0.66 \pm 0.09$ | $0.00 \pm 0.03$ |

To assess the GPR and RR models on unseen data points, models maximised on an independent validation set were retrained and assessed on additional held-out subject-level inputs (Table 1). This suggested demographics and clinical features related poorly to disease severity (Supplementary Fig. 11), while both RNA eigengenes and skin factors strongly related to disease severity in both the validation and testing datasets (Fig. 4a and Table 1). To demonstrate how our models can make clinically meaningful predictions, a random selection of subject predictions were plotted over time (Supplementary Fig. 13, 14).

To assess feature importance, we used SHAP (SHapley Additive exPlanations) which shows the contribution of each skin RNA module and skin factor for predicting disease severity (Fig. 4b) (31). Despite differences in model methodology (i.e. linear vs. nonlinear), both regression techniques prioritised similar eigengenes and skin factors, although GPR was more selective than RR (Fig. 4b and Supplementary Fig. 12b). Overall, both methods highlighted turquoise, darkred (PI3K-AKT-mTOR signalling), steelblue (innate/adaptive immunity), violet (complement) and tan (oxidative phosphorylation) modules as important contributors to disease severity prediction (Fig. 4b and Supplementary Fig. 12b). Factors S1 and S2 (mixed inflammatory), which are inter-correlated (Supplementary Fig. 3a), were identified by SHAP analysis of RR and GPR models respectively as the most influential skin factors for predicting PASI (Fig. 4b). Additionally, factors S9 and S6 (ECM signalling) appeared as top features influencing disease severity (Fig. 4b and Supplementary Fig. 12b). By analysing both MAE and R² values, we identified the turquoise module as the strongest positive and the blue module as the strongest negative contributors to disease severity prediction (Table 1). In order to gain a deeper understanding of key genes driving disease severity prediction, the GPR models were retrained with only the top 10 aligned genes of selected modules and factors correlating with disease severity (Fig. 4c and Supplementary Tables 7 and 8). A SHAP analysis of the top 10 aligned genes from turquoise and from blue identified *CARHSP1*, *KLK13*, *CRABP2*, and *GJB2* (turquoise, cytokine) and *THRA*, *ZNF34*, *RORC*, *CRY2* and *CACNA2D2* (blue, WNT signalling) as the 9 key genes driving disease severity prediction (Fig. 4d and Fig. 6d).

Factors S18 (HLA-DQA1*01 / HLA-DRB1*15-associated) and S21 (HLA-enriched factor C) were among the top predictors of disease severity according to both models (Fig. 4b and Supplementary Fig. 12b). Most of the genes highly aligned with these factors were HLA-encoding, with *HLA-DQB1*, *HLA-DQA1*, *HLA-DRA*, *HLA-DRB1* and *HLA-DRB5* aligning with factor S18, and *HLA-E*, *HLA-DQB2*, *HLA-DQA2* and *GSTM1* aligning with factor S21 (Fig. 5a). Previous studies have reported associations between HLA genotypes and psoriasis severity (32). We therefore investigated associations between HLA genotypes and the expression of genes and factors. Among the strongest genotype-gene correlations were those between *HLA-DQA1*01* genotype and the expression of *HLA-DQA1* and *HLA-DQB1*, and between *HLA-DRB1*15* and expression of *HLA-DRB5* and *HLA-DRB1* (Fig. 5b, c). Additionally, the two strongest genotype-factor relationships were between *HLA-DQA1*01* and *HLA-DRB1*15* genotypes and factor S18 (Fig. 5b). We observed three clusters of patients corresponding to factor S18, its constituent genes and the HLA genes. Individuals with both the *DRB1*15* and *DQA1*01* genotypes have high factor S18 expression and high expression of *HLA-DQB1*, *-DQA1*, *-DRB1* and *-DRB5*; subjects with the *DQA1*01* genotype only have moderate factor S18 expression and high expression of *HLA-DQB1* and *-DQA1*, while those with neither allele have low factor S18 expression and low expression of all four of the aforementioned genes (Fig. 5a).

**Figure 5.**
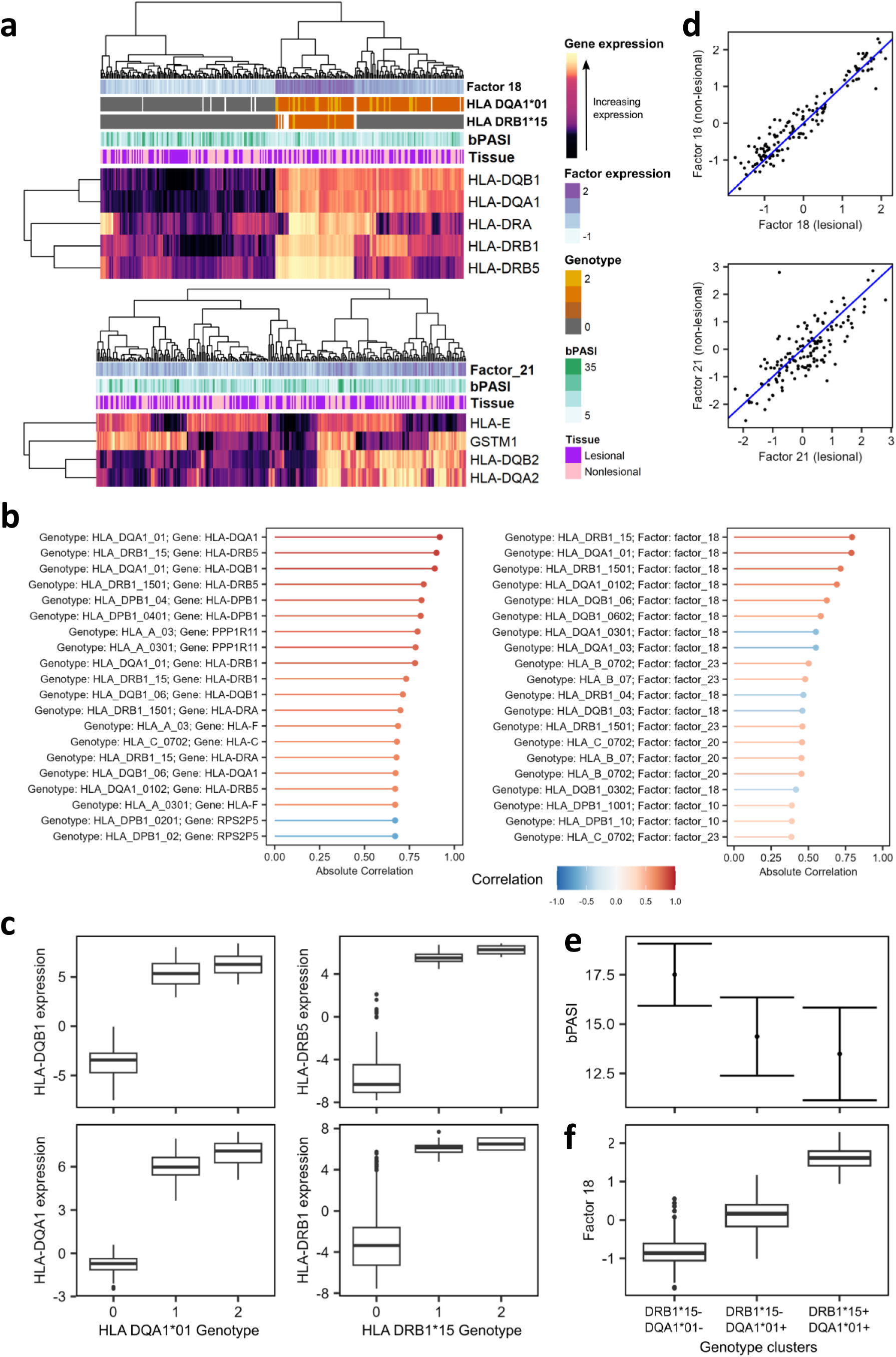
Latent factor S18 defines HLA-driven patient clusters associated with baseline disease severity. ICA factor S18, derived from combined lesional and nonlesional skin data, segregates patients into clusters differing in class-II HLA genotype and baseline PASI. **(a)** Two heatmaps for week-0 samples. **Upper:** expression of factor S18 and its highly aligned genes. **Lower:** expression of factor S21 (another HLA-related factor). Tracks show tissue type (lesional/nonlesional) and baseline PASI (bPASI). For S18, additional tracks display *HLA-DQA1*01* and *HLA-DRB1*15* genotype dosage (0/1/2; grey, orange, yellow respectively, white is missing). **(b)** Bar plots of top genotype-gene (left) and genotype-factor (right) correlations in lesional skin. **(c)** Boxplots of HLA gene expression stratified by genotype which show first quartile, median, third quartile, minimum and maximum values, and points to indicate outliers. **(d)** Scatter plot comparing S18 expression in matched lesional vs nonlesional skin; the identity line (blue) marks equal expression. **(e)** Estimated mean bPASI (95 % CI) by genotype from linear modelling. **(f)** Distribution of S18 expression by genotype in lesional skin. *Abbreviations:* HLA, human-leukocyte antigen; bPASI, baseline PASI; LS, lesional skin.

Given that these HLA-aligned factors were not associated with disease severity across timepoints (Fig. 2a), we investigated why they were considered important for prediction (Fig. 4c). There was no apparent clustering of samples by tissue type (Fig. 5a) and the expression of these factors displayed strong correlations between paired lesional and nonlesional samples (Fig. 5d), suggesting that the expression of these factors does not depend on tissue type. We next considered associations between the factors and baseline disease severity. Factors S1, S2, S17 (matrisome and cytokine receptor) and S18 were significantly correlated (FDR < 0.05) with baseline PASI in lesional skin; factor S18 was the most significantly associated (Pearson’s *r* = - 0.28, adjusted *p*-value = 0.01) (Supplementary Table 9). Furthermore, the patient clusters based on the HLA-DQA1*01 and HLA-DRB1*15 genotypes were significantly associated with baseline PASI (*p*-value = 0.007) (Fig. 5e, f). These data indicate a link between these genotypes, expression of *HLA-DQB1*, *-DQA1*, *-DRB1* and *-DRB5* and psoriasis severity at baseline.

### Disease severity endotype gene level analysis

#### Lesional skin

Extending our investigation to gene level (Fig. 6a), we found a significant nonlinear association between expression of 4108 genes in lesional skin and psoriasis disease severity (Fig. 6b, c). The majority (2926 [71.2%]) of these genes showed an association independent of biologic treatment modality. Approximately 40% of disease severity-associated genes showed an overall positive relationship and approximately 60% a negative relationship with disease severity (Fig. 6c). Previous studies have paid less attention to genes negatively correlated with PASI, but these may represent important regulatory pathways constraining inflammatory responses and represent novel therapeutic targets.

**Figure 6.**
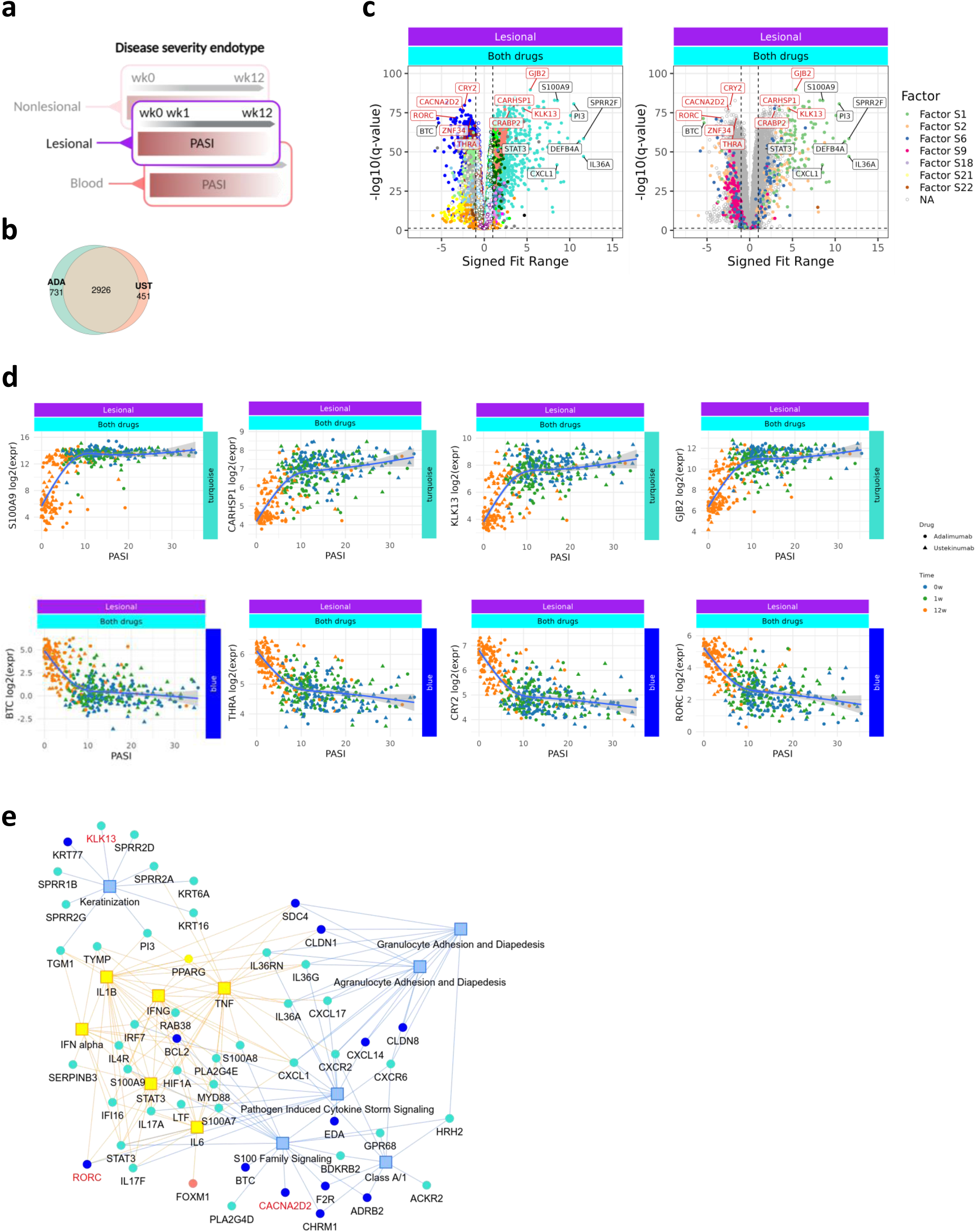
Disease severity endotypes in lesional skin are largely independent of biologic drug. **(a)** Workflow: non-linear (natural-spline 3 d.f.) regression was used to relate gene expression to PASI separately in adalimumab and ustekinumab cohorts; significant genes (*q* < 0.05) were combined for downstream analysis. **(b)** Venn diagram summarising the overlap of significant genes between treatment arms. **(c)** Volcano plot for the pooled dataset showing –log10 *q* versus signed fit range (difference between maximum and minimum fitted expression across the PASI span; sign reflects direction). Left panel colours genes by WGCNA module, right by ICA factor; exemplar genes are annotated. **(d)** Example turquoise- and blue-module genes plotted as log2-CPM versus PASI with spline fits. Curves represent natural-spline fits (3 d.f.) with 95 % confidence bands. **(e)** Network view (Ingenuity Pathway Analysis): circles = genes (module-coloured), yellow squares = predicted upstream regulators, blue squares = enriched canonical pathways. *Abbreviations:* ADA, adalimumab; UST, ustekinumab; others as before.

Of the genes which were associated with disease severity for both drugs, 33% (968 genes) were assigned to the turquoise module (cytokines) and almost 25% (698 genes) were assigned to the blue module (WNT), which were the top-ranked modules in the GPR model positively and negatively associated with disease severity, respectively (Fig 4b, 6c). Genes highly positively correlated with disease severity within the turquoise module are shown in Fig. 6d and Supplementary Fig. 15. Interestingly, the majority of the GPR-derived core gene signature driving disease severity prediction (highlighted in Fig. 6c-e) were not previously recognised to be psoriasis disease severity-associated. Network analyses identify S100 family signalling, keratinisation, interferon, chemokine, granulocyte diapedesis and matrisome signalling as key canonical pathways (Fig. 6e, Supplementary Fig. 16). WNT and collagen genes were prominent within the negatively-associated disease severity network (Supplementary Fig. 17). Network analyses identified TNF, IFN-gamma, STAT3, IL-1B and IL6 as key upstream regulators of disease severity-associated genes (Fig. 6e, Supplementary Fig. 16). 448 disease severity-associated genes were identified as being highly aligned with a factor. Half of these were turquoise module genes (246 [50.4%]), of which the majority of these positively aligned with factor S1 or factor S2, underscoring the similarity between these factors. Factor S9 genes were highly correlated with lightyellow (obesity, 103 genes), as reported above.

Additionally, in keeping with their correlation with single cell keratinocyte signatures (Supplementary Fig. 6), darkgreen (positively associated with PASI) and skyblue (negatively associated with PASI) contained cornification genes, including ABCA12, SBSN, KLK7, and PCSK6 in darkgreen, and FLG, LOR, GAN, and LCE5A in skyblue; the association of these genes with PASI is illustrated in Supplementary Fig. 15d. This highlights that different sets of skin barrier genes exhibit distinct patterns of co-expression and associations with disease severity, with some being positively associated with PASI and others negatively associated.

A small proportion (29%; Fig. 6b) of the disease severity-associated genes in lesional skin were drug-specific and these typically represented relatively subtle differences between the drugs (Supplementary Fig. 15b).

#### Nonlesional skin

Next, we investigated disease severity-associated genes in nonlesional skin. Unsurprisingly, we found that the numbers of disease severity-associated genes in nonlesional skin were much lower than lesional skin (145 genes; Supplementary Fig. 18a); similarly, the -log10 p-values and signed fit ranges were less than lesional skin. Several disease severity-associated genes in lesional psoriasis, such as *S100A7* and *PI3*, were also found in nonlesional psoriasis, whereas the cytokine genes *IFN-gamma*, *IL-17*, *IL-36* and *CXCL1* were notably absent (Supplementary Fig. 18a). Consistent with findings at the modular level and in contrast to lesional skin, 79.3% of genes in nonlesional skin were biologic-agent specific. Thus in nonlesional skin, 105 disease severity-associated genes were adalimumab-specific, 19 were ustekinumab-specific and 21 were common to both drugs (Supplementary Fig. 18a). Adalimumab disease severity-associated genes were enriched for interferon signalling, TNF signalling, keratinisation and granulocyte diapedesis pathways, similar to lesional skin, but not the matrisome canonical pathway, whilst ustekinumab disease-associated genes were enriched for keratinisation and antimicrobial peptides (Supplementary Fig. 18b, Supplementary Fig. 19).

#### Blood

As with nonlesional skin, we found that the numbers of disease severity-associated genes in blood were much lower than lesional skin (181 genes) with lower -log10 q values and signed fit ranges (Fig. 7b-c). Notably, and in line with our module analysis, 100% of the disease severity-associated genes in blood were specific to adalimumab (Fig. 7b-c) and systems analysis showed enrichment of genes related to neutrophil degranulation with gene membership to the navy (innate/adaptive immunity 1), plum (innate/adaptive immunity 2), khaki (innate immune cell) and thistle (inflammatory cell death) modules, in line with our modular analysis (Fig. 7d-e, Supplementary Fig. 20). The adalimumab-specific genes with the greatest disease severity association included the navy module genes *ADGRG3*, *VNN1* and *PADI4* and the thistle module gene *S100P* (Supplementary Fig. 15). Overall, these data highlight an important relationship between expression of neutrophil degranulation genes in blood and psoriasis disease severity in the skin in patients treated with adalimumab.

**Figure 7.**
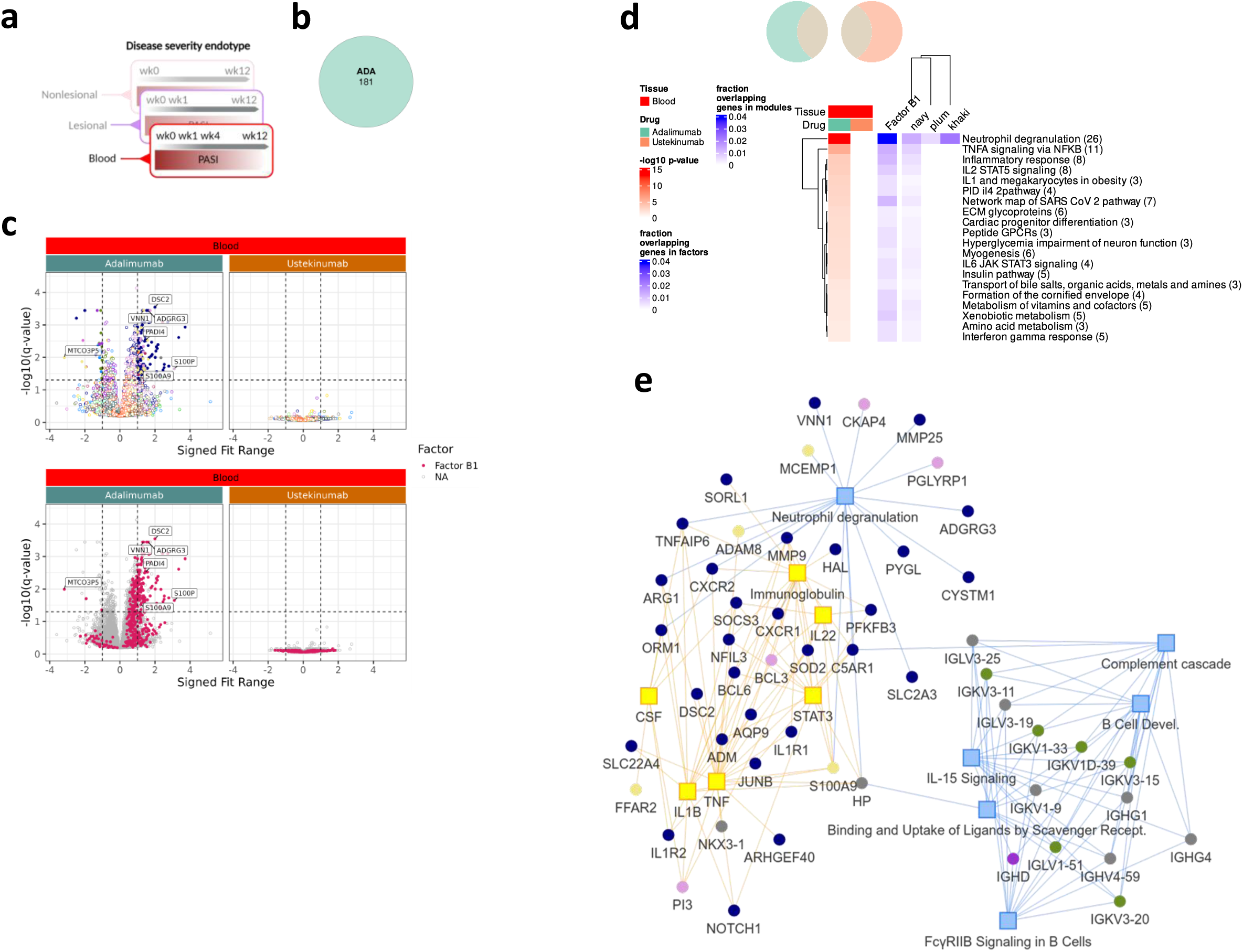
Disease-severity signatures in blood are specific to adalimumab. **(a)** Workflow schematic parallel to Fig. 6. **(b)** Venn diagram showing limited overlap of PASI-associated genes between blood of adalimumab and ustekinumab cohorts (*q* < 0.05). **(c)** Volcano plots for each drug: –log10 *q* versus signed fit range, coloured by WGCNA module (upper row) or ICA factor (lower row); illustrative genes labelled. **(d)** Metascape enrichment heatmap of co-expressed PASI-associated genes in each cohort. Red scale shows –log10 *P* for pathway enrichment; blue scale shows gene-set overlap fraction after module-size normalisation. **(e)** Network of adalimumab-specific PASI genes: nodes coloured by WGCNA module; yellow squares mark predicted upstream regulators and blue squares canonical pathways (Ingenuity Pathway Analysis). *Abbreviations:* as before; q, Benjamini–Hochberg FDR-corrected *P* value.

## Discussion

By integrating clinical, genomic and transcriptomic data derived from skin and blood, collected at baseline and across early time points during therapy of psoriasis, with two distinct classes of biologics, we identify reproducible endotypes (subgroups) of disease associated with distinct clinical phenotypes, disease severity and class of drug treatment. We found that endotypes were most accurately defined by core networks within bulk RNAseq data that represent direct and indirect interactions of genes across several cell types within tissue compartments. The disease severity endotypes in lesional skin were characterised by a core network of co-ordinated genes and pathways that were largely independent of drug treatment. Positive signatures of disease severity linked innate and acquired immunity whilst negative signatures of disease severity were characterised by matrisome and Wnt signalling pathways. The identification of core coordinated gene networks that are negatively associated with psoriasis disease severity is an important finding suggesting key regulatory and restraining functions of the interconnected networks in skin which require further study and potentially represent novel therapeutic targets. ICA but not WGCNA delineated gene signatures in blood that characterised the *HLA-Cw6* associated phenotype of psoriasis. In contrast, disease severity endotypes in nonlesional skin and blood were more specifically connected to the individual biologics. In blood, a key adalimumab disease severity-associated module (khaki, innate immune module) was functionally enriched in neutrophils and genes related to neutrophil degranulation, in line with the concept that the blood transcriptional signature of psoriasis is neutrophil driven (30). Taken together, our findings support a model in which adalimumab and ustekinumab exert differential effects across tissue compartments with adalimumab exerting greater effects in nonlesional skin and blood, in line with the pleiotropic effects of TNF, where ustekinumab effects appear more specifically targeted to the diseased tissue (lesional skin). These data are also consistent with reduced drug survival due to side effects (34) and increased serious infection risk with adalimumab compared to ustekinumab (35).

Our study aimed to investigate endotypes in psoriasis, a complex task that requires sophisticated and multi-dimensional analytical models. Single gene differential expression analysis has been widely used to explore the pathophysiological mechanisms of psoriasis, but potentially oversimplifies the complexity of gene interactions and regulation in biological systems, and may be confounded by measures of disease activity, such as PASI which is unlikely to be a simple linear relationship. This may help to explain the limitations of other transcriptomic studies to date, which to our knowledge have failed to independently replicate response biomarkers in psoriasis.

Applying WGCNA and ICA, two complementary methods, to the large number of samples in PSORT, we have delineated the complex direct and indirect interaction of genes across different cell types that potentially drive diverse phenotypes within a clinically homogeneous archetypal immune-mediated inflammatory disease (IMID) - psoriasis. Finding correlations between these two analyses can provide a more comprehensive understanding of gene interactions, as it suggests that these genes not only share expression patterns but also contribute to distinct biological signals or functions.

The co-ordinated expression between WGCNA modules and ICA factors was reflected in the two main clusters following correlation analysis (Supplementary Fig. 3a). The functional relevance of these two clusters is underscored by their anti-correlation with disease severity (PASI). Thus, strikingly, WGCNA and ICA identified a first block of modules and factors positively associated with disease severity which were also prioritised by machine learning methods and represented cytokine signalling, oxidative phosphorylation, antimicrobial proteins and the extracellular matrix pathways (Fig. 4b). Conversely the second block was negatively correlated with disease severity, representing Wnt signalling, insulin and hormone secretion and obesity-associated pathways (Fig. 4b).

Gene expression in skin has been previously linked to obesity (36) with some of these genes associated with psoriasis. Here, we define a 14-gene signature that negatively correlates with BMI within nonlesional skin and psoriasis disease severity within lesional skin, underscoring common pathways and mechanisms. Psoriasis is strongly associated with the metabolic syndrome, obesity and high BMI (37,38), supported by our previous study of 2889 biologic treated patients (21). Several of these core BMI/PASI signature genes are candidates for further mechanistic studies. For example, *DNER* (Delta/Notch Like EGF Repeat) regulates adipogenesis by modulating mesenchymal stem cell proliferation (39) and polymorphisms of *DNER* associates with risk of type II Diabetes in certain populations (40). In line with our data, *DNER* mRNA was upregulated in lesional psoriasis 12 weeks into therapy with fumaric acid esters (41).

The enrichment of the insulin and hormone secretion signature in sebocytes and the hair follicles following deconvolution is intriguing, given the enhancement of Wnt signalling pathways in our systems analysis. Interestingly, *SCGB1D2* (Secretoglobin Family 1D Member 2) expressed by sebocytes and sweat glands (Supplementary Fig. 8) (12) and previously reported to be down-regulated in psoriasis (42) was one of the top 5 down-regulated genes in white adipose tissue comparing obese insulin resistant to normal glucose tolerance individuals (43). Further, a recent spatial transcriptomic study identified cross-talk between cutaneous inflammation and lipid metabolism in immunocompetent sebaceous glands (44). *PDE9* regulates energy metabolism and inhibition of phosphodiesterase type 9 reduces obesity and stimulates lipogenesis via PPARs (45). Interestingly, a PPARg-expressing Treg cell population identified in a mouse model of psoriasis which suppresses IL17 expression was lost in obese mice (46). Furthermore, previous bulk RNAseq and spatial transcriptomic studies of psoriatic skin identified differentially expressed genes within sebaceous glands that were regulated by PPAR-g, the target of the diabetes drug, pioglitazone, which has shown efficacy in psoriasis (47); we observe *PPARGC1A* within the lightyellow module, in line with these findings. The intersection of psoriasis, metabolic syndrome, and type 1 diabetes highlights the systemic nature of IMIDs. As ustekinumab has shown efficacy in reducing inflammation and preserving insulin function, it could play a role in managing psoriasis patients who are at high risk of developing metabolic syndrome and type 1 diabetes. The strong negative association between hormone secretion/obesity-related genes and psoriasis severity suggests that molecular phenotyping could usefully be built into future therapeutic studies that target both metabolic (e.g. Glucagon-like peptide (GLP)-1 agonists) and inflammatory pathways.

The blood RNAseq data analysis in psoriasis subjects presented here is also the most extensive so far, both in terms of sample number and depth and breadth of variables explored; previous studies have only used data from 6 psoriasis subjects (48) or 23 subjects with psoriatic arthritis (49). Our ICA analysis indicated that factor B8 (inflammatory and HLA-aligned B, antigen processing and presentation) and factor B19 (inflammatory and HLA-aligned C, antigen processing and presentation) modules associated with HLA-Cw6 in a positive and negative manner respectively. While we do not fully understand the relationship between the Cw6 allele and psoriasis susceptibility, a number of mechanisms have been proposed by which the HLA-Cw6 antigen could directly initiate an immune response (50). The link between HLA-Cw6 and expression patterns in the 6p21 region could represent an additional mechanism by which the allele influences the development of psoriasis. Of the genes enriched in factor B8 module, class II *HLA-DRB1* has been associated with psoriasis susceptibility and may be druggable (51). Of the genes associated with factor B10 module, *HLA-DRA* encodes the invariable α-chain partner to DRB1, playing a central role in antigen presentation and *HLA-DMB* is crucial for MHC class II peptide loading.

Although Cw6 status has been linked to expression of individual genes, including *IL-1α* and *IL-6* (52), systems-level expression signatures associated with Cw6 status have not previously been identified, and whilst previous studies have found evidence connecting *HLA-Cw6* to disease presentation and severity (50), the Cw6-associated transcriptomic factors we have identified in blood were not correlated with disease severity. Therefore, these signatures likely indicate markers of psoriasis susceptibility, rather than severity.

Not surprisingly, the association of eigengene expression of modules in blood with cutaneous disease activity was less strong compared to modules in lesional skin. Nevertheless, the khaki (innate immune cell; neutrophil degranulation) module and factor B1 (inflammatory and HLA aligned A; phagosome) were positively and significantly associated with disease severity but only in the adalimumab cohort. Systems, deconvolution and gene level analysis indicated enrichment of neutrophils and genes related to neutrophil degranulation, consistent with previous reports that the blood transcriptional signature of psoriasis is neutrophil driven (30). The data reported here suggests that neutrophil activation in the blood is linked to disease activity and is modified by adalimumab during therapy. Gene level analysis in blood was consistent with our findings at a modular level with significant disease associated genes only being found in subjects treated with adalimumab. For example, increased disease severity-associated expression of the G-protein coupled receptor, ADGRG3, expressed by tissue-infiltrating granulocytes in blood but not in lesional or nonlesional skin is consistent with its known relative tissue specificity towards the cardiovascular system and liver (53). Together, our findings underscore neutrophils as an important therapeutic target of adalimumab with expression of key neutrophil degranulation genes in blood reflecting psoriasis disease severity in the skin and suggest that this signature is linked to TNF.

Genetic studies have identified genetic risk factors in the major histocompatibility region (MHC) for psoriasis that are independent of HLA-Cw6, including variants within the HLA-DQA1 gene (54). In the present study, we discovered three clusters of subjects based on the expression of the genes *HLA-DQA1*, *HLA-DQB1*, *HLA-DRA*, *HLA-DRB1* and *HLA-DRB5*. These clusters appeared to be determined by the HLA-DQA1*01 and HLA-DRB1*15 genotypes and were associated with PASI at baseline (Fig. 5). A transcriptome-wide association study found associations between three of these genes (*HLA-DQA1*, *HLA-DQB1* and *HLA-DRB1*) and psoriasis (55). Furthermore, HLA-DQA1*01:04 has previously been shown to be more common in patients with psoriasis (56). Interestingly, a specific haplotype comprising HLA-DRB1, -DQA1 and -DQB1 has been linked to protection against the development of type 1 diabetes (57). Taken together, these results suggest that a haplotype combining HLA-DQA1*01 and HLA-DRB1*15 genotypes could mediate psoriasis susceptibility and severity through the expression of a small number of HLA genes. These genes encode MHC class II surface proteins that are expressed by professional antigen-presenting cells, including Langerhans cells, with key roles in the maintenance of self-antigen tolerance. Their link with psoriasis may help explain the presence of autoreactive T cells in psoriatic tissue (58).

The complex interplay of a multitude of underlying drivers of biologic responses means that the relationships between gene expression and clinical parameters such as disease severity may be best described by non-linear relationships. Given the relatively limited number of patients available (for machine learning) and incomplete understanding of prior relationships, a Gaussian Process Regression (GPR) probabilistic model was explored, as GPR has been shown to model numerous complex biological relationships such as protein-function relationships with a limited amount of data (59–62), with a linear ridge regression (RR) model acting as a baseline comparison. Although both GPR and RR modelled disease severity with similar accuracy, GPR was more selective than RR, in part due to its ability to flexibly model features more independently as a sum of functions rather than aggregating features and modelling them as part of a single function as is done in the RR. Overall, our model’s performance explains on average half of the variation with disease severity. The probabilistic properties of the Bayesian GPR model provides uncertainty estimates allowing the assessment of unseen patients even in the absence of testing data, which can be further explored in future work.

Expression of disease severity-associated genes essentially reflects the molecular profile of skin or blood in psoriasis subjects that closely mirrors disease activity across the whole body (captured by PASI). Given that the skin transcriptome data was derived from small (5mm) lesional and nonlesional biopsies predominantly taken from the lower back, our findings support a co-ordinated disease response that represents integration of signals across cutaneous and circulating compartments.

The majority of modules and factors associated with disease severity in lesional skin were independent of biologic treatment class, in part reflecting clinical response and clearance of cutaneous psoriasis induced by both adalimumab and ustekinumab over 12 weeks. These findings are consistent with those of Brodmerkel et al., 2019 (63) who showed convergence of gene expression in response to ustekinumab and etanercept (another TNFi) over 12 weeks.

Of the genes associated with disease severity for both drugs, the majority were assigned to the turquoise (cytokines) (which was closely aligned to factors S1 and S2) and blue modules (WNT) (which was closely aligned to factors S9 and S22). Systems analysis showed that the turquoise module comprises genes related to cytokine signalling, antimicrobial peptides and cornification, that are known to play key roles in psoriasis pathogenesis, including S100A8/A9, IFN-gamma, IRF7, STAT3, CXCL1, DEFB4A, IL-17A/F, IL-36A, PI3 and cornification genes (e.g. IVL and SPRR1A) In line with the current understanding of the role of innate and acquired immunity in psoriasis pathogenesis (1), bulk RNAseq cell deconvolution demonstrated that turquoise (cytokine signalling) correlated most closely with myeloid-1 (dendritic cells/macrophages), T cell-2 (Th1, Th17), T cell-3 (cytotoxic T lymphocytes), myloid-1, venule-2 cells and keratinocyte-1, 4, (spinous) 6 (suprabasal) and 7 (basal) subsets. In line with recent literature, highlighting potential role of inflammatory fibroblasts in psoriasis pathophysiology (64), the blue module (WNT signalling) included genes such as BTC, an epidermal growth factor receptor (EGFR) ligand which regulates skin homeostasis and growth. Notably 100% blue (WNT signalling) module genes were negatively associated with disease activity. CYP2W1, an “orphan” member of the cytochrome P450 monooxygenase family may play a role in retinoid and phospholipid metabolism(51, 52) and has previously been characterised as one of top down-regulated genes in psoriasis (67). Metascape analysis placed BTC within the matrisome and deconvolution localised expression to fibroblast-5, fibroblast 2 (universal/reticular) (25,28), keratinocyte-1 (spinous), 5 (supra-spinous) and VSMC cells (Supplementary Fig. 6, keratinocyte-1 (spinous), 5 (supra-spinous) and VSMC cells (Supplementary Fig. A). Network analysis with IPA revealed BTC was associated with the S100 signalling pathway and was upregulated by CD36, along with CCL27 (a skin-specific memory T-cell chemokine that is reduced in lesional psoriasis, facilitating IL-17/-22 T-cell overactivity(68)) (Fig. 6e, Supplementary Fig. 16). CD36 is a cell-surface scavenger receptor with numerous functions including the regulation of angiogenesis, innate immunity and lipid metabolism (69).

Using GPR and SHAP, we identified a 9 gene signature driving PASI prediction (Fig. 4d). This list comprised 4 genes in the turquoise module, positively associated with PASI: *CARHSP1*, *KLK13*, *GJB2* and *CRABP2*; interestingly, these have all been associated with psoriasis previously (70–73), but to our knowledge none have been associated with psoriasis disease severity. The remaining 5 genes in the signature mapped to the blue module and were negatively associated with PASI: *THRA*, *ZNF34*, *RORC*, *CRY2* and *CACNA2D2*. *THRA* encodes for Thyroid Hormone Receptor Alpha and is thought to play a role in healthy skin development and homeostasis (74). This is the first report of an association between *THRA*, *ZNF34* and *CACNA2D2* with psoriasis and underscores the potential for novel drug targets amongst genes negatively associated with PASI. Notably, *RORC* and *CRY2* are core clock genes. It has previously been estimated that the circadian clock - representing a group of autoregulatory transcription factors - governs the rhythmic expression of half of all the genes in the human body (75). Circadian rhythms are disrupted in several inflammatory diseases including psoriasis, in which clock gene expression including *CRY2* but not *RORC* has been shown to be disrupted (76). A core gene signature, comprising a limited number of genes, is more easily assayed than a gene module (which might comprise hundreds, or even thousands, of genes) and therefore represents a potential biomarker that can be tested for clinical utility.

Strikingly, the expression levels of several modules in nonlesional skin were also significantly correlated with PASI. This is interesting in light of work by Gudjonsson et al (2009) (29), which found that nonlesional skin is transcriptionally “pre-psoriatic”, and suggests that reduction in disease activity over the course of treatment is accompanied by systems-level changes in expression in nonlesional skin as well as lesional skin and further underscores the systemic nature of psoriasis beyond the visible plaques.

In nonlesional skin and to an even greater extent in blood, there was a much greater disparity between the drug groups with respect to disease severity-associated modules, factors and genes. These findings suggest that adalimumab, by targeting the pleiotropic cytokine TNF, leads to systemic transcriptome modulation in nonlesional skin and blood (in addition to lesional skin) whereas the transcriptome changes induced by ustekinumab through inhibition of IL-12/23 are largely restricted to lesional skin. Moreover, this disparity with greater modulation of both innate and acquired immune pathways in blood by adalimumab compared to ustekinumab is consistent with a relative increase in secondary infections including TB with adalimumab compared to ustekinumab (35).

This experimental medicine study utilised the first classes of biologics introduced for psoriasis. Although there are now newer classes of biologics available that are more effective for managing psoriasis, adalimumab and ustekinumab continue to represent the most widely prescribed biologics in clinical practice in many countries, including the UK (34). Although our study of the psoriasis blood transcriptome is the largest reported to date, and reproduces a neutrophil signature (30), further independent validation analysis is required with respect to the disease severity endotype identified exclusively in the blood of patients treated with adalimumab. In contrast to our combined analysis of lesional and nonlesional psoriasis, our separate analysis of skin and blood precluded direct comparisons between these tissue compartments.

The systematic nature of the dataset across skin and blood and its integration with key clinical features and outcomes to therapy provides a comprehensive and global perspective of psoriasis that are not apparent from previous studies. Prior pharmacogenomic evaluations of patient cohorts have centred on the use of genetic or genomic techniques, predominantly using skin biopsies, although several studies have used skin and blood (57, 58). Importantly though, no prospective biomarkers of disease severity or response have yet been validated in adequately powered cohorts (79). Our results are fully reproducible via companion R markdown documents and, as in our pilot investigation, we encourage readers to replicate our analysis in independent cohorts, and we continue to advocate the open access of data and code for all studies to improve reproducibility and drive excellent science (20). Our workflow can be applied to evaluate datasets of disease severity and clinical response to novel therapies outside of dermatological indication. Further work is required to derive scalable biomarkers for testing in the clinical setting. One of the advantages of co-expression analysis is dimensionality reduction which facilitates the further development of scalable and clinically applicable biomarkers. Thus, for example, in our on-going work we are investigating the relationship between co-expression modules, clinical traits, a wider range of genetic markers and eQTLs though machine learning. Integrated analysis of samples across drug cohorts, time points, and lesional and non-lesional skin when carrying out WGCNA and ICA greatly reduced the dimensionality of the dataset. However, this approach precluded identification of modules and factors which might be specific to lesional or non-lesional skin, or to certain time points; future work might involve identification of modules and factors separately in these different subsets of samples. Methods such as module preservation analysis for WGCNA, which has been used in this study to compare skin and blood (Supplementary Fig. 4), could then be used to assess the degree to which module co-expression changes across tissues and time points. Crucially, differences in co-expression may be separate to differences in absolute expression levels, which are the focus of our endotype analyses in this paper and more suited to the integrated dimensionality reduction approach taken here. We are planning a separate paper that will examine time point-associated gene expression changes to define drug endotypes.

The authors believe it will be standard practice that molecular phenotyping will drive personalised therapy across therapeutic targets for immune-mediated inflammatory diseases in the future. Potential limitations of the study include lack of randomisation of subjects to treatment arms (although the study cohorts were well-matched) and the absence of treatment adherent assessment (although the response rates seen suggests that participants were adherent to therapy). The insights provided here of the complex interface between these disease and disease severity endotypes provides a framework for delineating molecular and clinical response across tissues and predicting individual subject response in future studies.

## Acknowledgements

*Non-author contributions*

We are grateful to all subjects for their participation. We thank the Independent Advisory Board who provided valuable advice and an independent international stakeholder perspective to the PSORT Consortium (Iain McInnes [Chair], John Armstrong, Anne Bowcock, James Krueger, Christy Langan and Peter van de Kerkhof). We are grateful to the Psoriasis Association for their Patient and Public Involvement and Engagement. We acknowledge the enthusiastic collaboration and support of dermatologists, specialist and research nurses in the UK who recruited to this study including Alberto Barea (Kingston Hospital NHS Foundation Trust), Dr Anna Chapman (Lewisham & Greenwich Trust), Dr Rob Ellis (South Tees Hospitals NHS Foundation Trust), Dr Abigail Fogo (Kingston Hospital NHS Foundation Trust), Dr Bronwyn Hughes (Portsmouth Hospitals University NHS Trust), Dr Evmorfia Ladoyanni (Dudley Group NHS Foundation Trust), Dr Philip Laws (Leeds Teaching Hospitals NHS Trust), Dr Richard Parslew (Liverpool University Hospitals NHS Foundation Trust), Dr Gayathri Perera (Chelsea and Westminister Hospital NHS Foundation Trust), Dr Beth Poyner (Newcastle upon Tyne Hospitals NHS Foundation Trust), Sara Wilkinson (Newcastle upon Tyne Hospitals NHS Foundation Trust). We thank Hira Ali, Rosa Andres-Ejarque, Zaynep Catak, Tejus Dasandi, Nadya Dinev, Michael Duckworth, Katarzyna Grys, Freya Meynell, Alice Russel and Isabella Tosi (London), and Dhanisha Lukka and Panagiotis Maniatis (Newcastle) for sample and data management. We thank Federica Villanova (London) for her contribution to obtain ethical approval.

*Funding*

The Medical Research Council (MRC)(MICA MRC Precision Medicine Consortium; MR/L011808/1).

The Psoriasis Association.

Partners of the PSORT consortium are AbbVie, the British Association of Dermatologists, Becton Dickinson and Company, Celgene Limited, GlaxoSmithKline, Guy’s and St Thomas’ NHS Foundation Trust, Eli Lilly, Janssen Research & Development, King’s College London, LEO Pharma, MedImmune, Novartis Pharmaceuticals UK, Pfizer Italy, the Psoriasis Association, Qiagen Manchester, Queen Mary University of London, the Royal College of Physicians, Sanquin Blood Supply Foundation, the University of Liverpool, the University of Manchester, and Newcastle University. We particularly acknowledge generous in-kind support from the PSORT industrial partners GSK and Abbvie who supported the RNA sequencing. All decisions concerning analysis, interpretation, and publication are made independently of any 13 industrial contributions.

The British Association of Dermatologists (BAD). The Rosetrees Trust.

UCB.

The NIHR Newcastle Biomedical Research Centre (BRC).

MRC/BAD/British Skin Foundation clinical research training fellowship to HJG.

NJR’s research/laboratory is funded in part by the NIHR Newcastle HealthTech Research Centre in Diagnostic and Technology Evaluation and the NIHR Newcastle Patient Safety Research.

Collaboration. NJR is a NIHR Senior Investigator.

CEMG and RBW are funded in part by the Manchester NIHR BRC (NIHR203308).

MRB and DW were funded by the NIHR as part of the portfolio of translational research of the

NIHR BRC at Barts and The London School of Medicine and Dentistry.

CS receives research funding from European consortia with multiple industry partners (see BIOMAP-imi.eu and HIPPOCRATES-imi.eu, MRC-industry PhD studentships from AstraZeneca, Boehringer Ingelheim).

This project was enabled through access to the MRC eMedLab Medical Bioinformatics infrastructure supported by the Medical Research Council [grant number MR/L016311/1].

## Author contributions

Conceptualization: MRB, JNB, CEMG, NJR, CHS, DS, RW

Data Curation: AR, DSW, ND

Formal Analysis: HJG, AR, DSW, GRS, JC, JG, SJC, SN, WAI

Funding acquisition: MRB, JNB, CEMG, NJR, CHS, DS, RW, PZ

Investigation: ACF, SKS, TE, CT, ET, DKR, KMS, JNB, CEM, CHS, RW, NJR

Methodology: DS, SKS, CT, ET, DKR, KMS, WAI, ND

Project administration: MRB, JNB, PD, CEMG, NJR, CHS, RW

Resources: MRB, JNB, CEMG, NJR, CHS, RW

Software: RH

Supervision: MRB, NJR

Validation: HJG, AR, GRS, JC, JG, WAI

Visualization: HJG, AR, GRS, JC, JG, WAI

Writing – original draft: HJG, AR, JG, WAI, MRB, NJR

Writing – review & editing: all authors

## Competing interests

NJR has received research grants from GSK-Stiefel and Novartis; and other income to Newcastle University from Almirall, Amgen, Janssen, Novartis, Sanofi Genzyme Regeneron and UCB Pharma Ltd for lectures/attendance at advisory boards. MRB has received research funding from Janssen and Benevolent AI, and consultancy with United Health group, Eli Lilly and Sanofi.

