## Supplementary text, figures, and tables 1 and 6-9 for "Transcriptomic profiling and machine learning uncover gene signatures of psoriasis endotypes and disease severity"

### Supplementary Materials and Methods

#### Prospective observational study

This study included 146 subjects with moderate to severe chronic plaque type psoriasis (PASI>10) recruited prospectively into the Psoriasis Stratification to Optimise Relevant Therapy (PSORT) study at 6 centres in the UK between May 2015 and May 2018 and due to start biologic therapy (ustekinumab or adalimumab) as part of routine clinical practice (1). Exclusion criteria included use of systemic or biologic treatments in the two weeks prior to study entry (or four x t½ of last treatment, whichever was longer), use of PUVA therapy for 3 months or UVB for 1 month prior to study entry or use of topical treatments at the site of biopsies (except for emollients) for 2 weeks prior to study entry, as well as serious/uncontrolled systemic disease. We studied 89 psoriasis subjects within the discovery cohort and replicated findings in a further cohort of 57 subjects. For the replication cohort, samples for RNA sequencing were selected from patients whose treatment response (PASI 50/75/90 and non-responders) broadly matched those in the earlier discovery cohort. Selection was based solely on clinical response and did not consider molecular or demographic data. Subjects commencing adalimumab by subcutaneous injection received 80mg at baseline then 40mg at week 1 then 40 mg every 2 weeks as per label and those starting ustekinumab received 45mg or 90mg according to body weight, as per label. Participants self-administered doses that fell between study visits and the time and date of these were recorded in the Case Report Form. The Psoriasis Association provided Patient and Public Involvement and Engagement which influenced the study design. The study was conducted in accordance with the declaration of Helsinki, was approved by the London Bridge research ethics committee (REC reference: 14/LO/1685PSORT) and subjects provided written, informed consent.

Patients completed detailed demographic questioning, including reporting information on comorbidities and concomitant and previous medication. Disease severity and response to therapy were assessed using the PASI, Physician Global Assessment (PGA) and DLQI. Clinical samples including blood and lesional skin punch biopsies (edge of psoriasis plaque with site preference for lower back or buttock) were collected under local anaesthetic at baseline, one week (prior to the second injection of adalimumab) and 12 weeks of treatment. Biopsies were derived from the same body sites at each time point and, if possible, lesional biopsies were taken from the same plaque as baseline. Non-lesional skin with a minimum distance of 2 cm from the edge of nearest plaque was also collected at baseline and week 12 (and a minimum distance of 2 cm between initial and subsequent biopsies) and a further blood sample was taken at 4 weeks. Patients had been identified in line with recommendations for initiation of biologic therapies in the UK. Screening investigations had been completed prior to recruitment. Adverse events were recorded but did not form part of primary analysis.

**Power calculation**

Based on our affiliated pilot investigation (2), using the method of Guo et al (3), we calculated the requisite sample size to achieve 90% power to detect differential expression associated with response. Imposing a 5% FDR threshold and a target log fold change of 1.5, we determined that a study would require 40 subjects to achieve 90% power to identify transcriptomic markers of biologic response for patients with chronic plaque psoriasis. Power curves projected across an expected range of fold changes at 1% and 5% DE in Supplementary Figure 1.

**RNA extraction and quality control**

*Skin samples*

RNA was preserved in the skin punch biopsies using RNAlater Stabilization Solution (Invitrogen AM7022). Biopsies were stored at 4°C in RNAlater overnight and the solution removed prior to long term storage at -80°C (according to the manufacturer's instructions).

Biopsies were transferred into pre-cooled 2ml lysing tubes (Precellys, CK mix) containing lysis buffer (10µL 2-ME/mL RLT Plus) supplied in the Qiagen AllPrep DNA/RNA Kit (Cat. No. 80204). Homogenisation was performed in the TissueLyser LT (Qiagen Cat. No. 85600) over 10 x 2-minute cycles at 50Hz. Samples were cooled on wet ice for one minute between cycles. Tissue debris was pelleted at 13,000 RPM for 3 minutes and the supernatant transferred to an Allprep DNA spin column. DNA/RNA was then extracted following the Qiagen AllPrep kit's protocol.

RNA concentration/integrity was checked using the Agilent Bioanalyzer 2100 with the RNA 6000 Nano assay (Agilent: 5067-1511); only samples with an RNA integrity number (RIN) of 8 or more were sequenced.

*Blood samples*

RNA was isolated from human whole blood collected in PAXgene blood RNA tubes (Qiagen #762165) utilizing the QIAsymphony SP (Qiagen #9001297) with the QIAsymphony PAXgene Blood RNA kit (96) (Qiagen #762635). The manual processing of the whole blood samples was performed following the manufacturer's protocol and loaded onto the QIAsymphony SP. The RNA isolation protocol implemented was a custom protocol based on the standard automation protocol PAXgene_RNA_V5.xml. The custom protocol is PAXRNA_CR22332_2915.xml and contains the following modification to the standard protocol “elution buffer taken out of accessory trough. Accessory trough will be displayed as ETOH on the touch screen.” The elution buffer Genomic Technologies Sample Management and Processing team uses is Invitrogen UltraPure DNase/RNase free distilled water #10977015. Whole blood RNA samples were eluted into 80ul of Invitrogen UltraPure DNase/RNase free distilled water #10977015 and plated into 8x12 elution plates. RNA quality control for quantity was performed with Qubit RNA Broad Range (BR) (ThermoFisher Scientific #Q10211) on a Molecular Devices Gemini plate reader following manufacturers protocol for reagent and sample preparation. RNA quality control for integrity was performed with an Agilent TapeStation 4200 (Agilent #G2991BA) using the RNA Screentape assay (Agilent #5067-5576,77,78) following manufacturer’s protocol.

**RNA sequencing**

The sequencing libraries for the PSORT-D skin samples were prepared using the Illumina Truseq stranded mRNA kit and sequenced on an Illumina HiSeq 3000 with 2x101bp read length. The sequencing libraries for the PSORT-D blood samples were prepared from total RNA using the Kapa mRNA HyperPrep kit and depleted of rRNA and globin mRNA using the QIAseq FastSelect RNA Removal Kit by Qiagen; sequencing was done on an Illumina HiSeq 4000 using 2x75bp read length. The sequencing libraries for the PSORT-R skin samples were prepared using the Illumina Truseq stranded mRNA kit and sequenced on an Illumina NovaSeq 6000 with 2x250bp read length. As the approved participant consent forms did not include permissions for sharing of (personally identifiable) raw sequencing data, we provide raw and adjusted gene count data from our RNA-seq analysis.

**Genotype data and HLA imputation**

DNA was isolated from blood using standard methods. Genotyping was performed with Illumina HumanOmniExpressExome-8 v1.2 and v1.3 BeadChips, followed by quality control with standard tools, as previously described (4). *HLA-C*06:02* genotype was imputed using SNP2HLA (version 1.0.3) based on the Type 1 Diabetes Genetics Consortium reference panel (5).

**Genomic and Transcriptomic data analysis workflow**

Analysis was conducted in the R statistical computing environment (R Core Team, 2021).

A graphical summary of the samples used for analysis is available in supplementary figure 21. Several blood samples were RNA sequenced but identified as technical failures based on exploratory analysis and so were excluded. Following RNA-sequencing of all samples, reads were pseudo-aligned using Kallisto (6). Transcript counts aggregated gene-wise and were TMM normalised prior to modelling (7) and transformed to the $log_2$-CPM scale. An expression filter was applied to ensure that a gene has at least one count per million (CPM) in at least 5% of all libraries, leaving 16,172 genes.

The processed data was then subjected to detailed exploratory data analysis as reported in the supplementary material. Gene co-expression modules were identified using both Weighted Gene Correlation Network Analysis (WGCNA) and Independent Component Analysis (ICA) and extensively explored using enrichment techniques (Metascape) and deconvolution analysis (CibersortX).

**Exploratory Data Analysis**

Principal component analysis identified tissue, time, and disease activity as the key drivers of transcriptome variation (Supplementary Fig. 22c), with relatively limited influence of demographic and other factors previously associated with therapeutic response.

**Differential Expression Analysis**

Differential expression was tested using heteroskedastic linear models and empirical Bayes shrinkage as implemented by the voom function *in* the limma software package (8). We compute *q*-values for each differential expression test using Storey's method (9), with a false discovery rate (FDR) threshold of 5%.

We built separate models to test a number of related hypotheses. We have two primary goals for this portion of the experiment: (1) to identify genes that associate with disease phenotype (e.g BMI) and (2) to identify genes that correlate with PASI irrespective of time points. We refer to these as the disease, and disease severity endotypes, respectively.

*Disease Severity Model*

Disease activity is measured at each time point by the psoriasis area severity index (PASI) score. Building on the design of our trial study (2), we split the data by tissue and analysed samples from both treatment arms with coefficients for each drug. We accounted for repeated observations using the *duplicateCorrelation* function, which approximates a mixed model design in which patient ID is treated as a random effect. Because preliminary investigations suggested that gene expression is often a nonmonotonic function of PASI, we expanded the model using a cubic spline basis of degree 3. An intercept term was included for each drug and the spline coefficients were also allowed to vary depending on the Drug, except for the model underlying the volcano plot in Figure 6, which was independent of Drug. In the lesional skin and nonlesional skin models, an intercept for the Cohort (Discovery/Replication) was included. In addition to the q-value criterion, we define a signed fit range equal to the minimum to maximum range of the fitted log2CPM expression, with the sign given by the sign of the gradient at PASI = 0. In the Venn Diagrams and Volcano plots of Figures 6 and 7, we include only those genes that were assigned to a WGCNA module (see section *Identification of gene coexpression modules* below).

**Dimensionality reduction**

WGCNA and ICA decompose transcriptomes of many thousands of gene transcripts into a dataset comprised of a much smaller number of gene modules, overcoming the limitations inherent in gene-level analysis, including lower signal to noise ratios and a higher multiple testing burden (10), enhancing the statistical power to detect true endotype associations. These methods also reduce data complexity, offering a more holistic view of biological pathways and networks by focusing on co-expressed genes organised mutually exclusively into modules (WGCNA) or independent components (ICA), which reduces high-dimensional data into a smaller set of latent variables (or factors), with the objective of describing unobserved processes that explain patterns in gene expression, allowing for genes to belong to multiple pathways. Each module is represented by an eigengene and each factor by a metagene. Eigengenes and metagene represent summary expression values for modules and factors, respectively. These approaches not only provide robustness against noise but also reveal biologically relevant patterns and potential novel mechanistic insights which may be missed in individual gene level analyses.

Whereas individual genes can only be assigned to one up or down regulated WGCNA module, ICA allows single genes to be weighted across multiple factors in both up or down regulated states, reflecting the involvement of genes in multiple shared biological processes (Fig. 1d). These approaches are complementary, with the former being more explainable and the latter more likely to represent biological complexity.

Within skin and blood, all samples were used for WGCNA and ICA, i.e. both lesional and nonlesional samples (within skin), both drug cohorts and all time points.

**Identification of gene coexpression modules**

The following steps were carried out for the PSORT-D skin and blood data separately in order to identify co-expressed gene modules in each tissue compartment. Prior to running WGCNA, the gene-level counts were filtered using a threshold that required at least one CPM in at least *n/k* libraries, where *n* equalled the number of samples and *k* equalled the number of unique combinations of tissue type, drug, and time point. The counts were then normalised using the TMM method and transformed to log2-CPM. Selection of the appropriate soft-thresholding power β was done by plotting the values 1-20 against $R^2$, a measure of scale-free topology, and mean connectivity. The lowest value which reached the $R^2$ threshold of 0.8 was chosen; a β of 12 was chosen for skin and 5 for blood (Supplementary Fig. 23). The *blockwiseModules* function was then used to partition the genes into co-expressed modules. In brief, this function calculated the Pearson correlations between each pair of genes and raised these estimates to the selected β power in order to amplify the differences between high and low correlations. These correlations were then used to generate a topological overlap matrix (TOM) and hierarchical clustering of this matrix was used to group genes with similar expression profiles into modules. Parameters to *blockwiseModules* included a minimum module size of 30, a dendrogram cut height (for merging of similar modules) of 0.1, and the use of a signed network so that the correlations between genes were scaled to lie between 0 and 1.

**Module-trait correlations**

The *moduleEigengenes* function was used to derive eigengene values for the skin and blood modules in every sample. An eigengene represents a summary expression score for a module and is analogous to the first principal component of the expression matrix for that module. Although module identification was not carried out in PSORT-R, the module assignments in PSORT-D were used to derive eigengenes for these modules in PSORT-R as well. Pearson correlation (with pairwise complete observations) was used to identify associations between modules and traits of interest. These included disease traits at baseline: age of onset, onset type (early/late), anti-TNF naïve status (Y/N), PsA (positive/negative), sex, age, BMI, and Cw6 status (positive/negative); PASI across time in each drug cohort; and the treatment effect at week 1 and week 12 in each drug cohort. Binary traits were encoded as one and zero. In skin, the module-trait correlations were carried out separately for the lesional and nonlesional samples. Significant correlations were defined by FDR <= 0.05; in skin, replicable correlations were defined by FDR <=0.05 in the discovery cohort and nominal p-value <=0.05 in the replication cohort and correlation of the same sign in both cohorts. Only traits with at least one significant correlation are displayed in the module-trait correlation heatmaps.

**Independent component analysis**

Independent component analysis (ICA) was applied to identify latent variables separately in both skin (discovery cohort) and blood expression data. In each case, we included samples from both treatments and all timepoints, and we centred the data prior to factor analysis. We used the "imax" method implemented in the ica R package (11), using the maximally stable transcriptome dimension (MSTD) approach to select the optimal number of factors to compute (12) implemented in the ReducedExperiment package (13). The number of factors recommended by MSTD was 24 for the skin expression data and 21 for blood. In order to validate the identified skin signatures, we projected the expression data from the replication cohort into the factor-space defined in the discovery cohort. As a result, the feature loadings (i.e., the source signal estimates) for the skin discovery and replication cohorts are equal, permitting the investigation of the same factors in each cohort.

Factor metagenes were calculated by taking the scaled values of the estimated mixing matrix (Supplementary File Factor tables). These factor metagenes were then associated with phenotype using the same modelling approach as we employed for the module eigengenes (see Module-trait correlations, above). For factors, we additionally calculated correlations with HLA genotypes and baseline PASI (bPASI). This was carried out using the combined discovery and replication cohorts following batch effect correction using limma's removeBatchEffect function. Whereas the WGCNA modules contain non-overlapping sets of genes, the factors identified by

ICA are each aligned with all genes. For analyses that required a defined set of genes, we selected a set of highly aligned genes for each factor based on their loadings (Supplementary File Factor tables). We selected genes with loadings that exceeded a threshold. By default, we defined this threshold at half the maximal loading for that factor (Supplementary File Factor tables). For some analyses (functional enrichment analysis, BMI associations) we used a more relaxed threshold of 5; where this method resulted in the selection of less than 20 genes, we instead extracted the top 20 features.

**Deconvolution**

Abundance of cell types was inferred using the CibersortX online tool (14). To infer cell types in skin, a single-cell reference matrix was generated using single-cell RNA-Sequencing data from 38,274 skin cells across 5 inflammatory skin conditions including psoriasis (15) downloaded from the Single Cell portal developed by the Broad Institute of MIT and Harvard (https://singlecell.broadinstitute.org/single_cell). Due to memory limitations imposed by the CibersortX tool, the size of the reference matrix was reduced by downsampling to a maximum of 200 cells per cell-type. The reference matrix was used to generate a Signature Matrix file using recommended CibersortX settings. Bulk RNA-Seq data for both PSORT discovery and replication cohorts was used to generate the mixture file. Cell fractions were imputed in absolute mode using 'B-mode' batch correction.

To infer cell types in blood, the LM22 signature matrix provided by CibersortX was used. The mixture file was generated from PSORT blood RNA-Seq data. Cell fractions were imputed using recommended CibersortX settings.

**Correlations of cell type fractions with latent factors and module eigengenes**

Per-sample imputed absolute cell fractions for each cell type were tested for association with per-sample WGCNA module eigengene loadings and with per-sample Latent Factor loadings using the cor.test function from the R statistical computing environment. Correlation tests used Kendall's tau coefficient. Adjusted p-values were calculated with the p.adjust function using the method of Benjamini & Hochberg.

**Pathway analysis**

Functional analysis of systems-level upstream regulators responsible for observed differential gene expression related to response was performed using the Upstream Regulator function in Ingenuity Pathways Analysis (16), using all genes with nominal response $p \leq 0.05$ as input. For all gene set enrichment analyses, a right-tailed Fisher's exact test was used to calculate a pathway p-value determining the probability that each biological function assigned to that data set was due to chance alone. All enrichment scores were calculated in IPA using all transcripts that passed QC as the background data set. Upstream regulator analysis is based on prior knowledge of expected effects between regulators and their known target genes according to the IPA database. The prediction of activation state is based on the global direction of changes of differentially expressed genes, a z-score is calculated and determines whether gene expression changes for known targets of each regulator are correlated with what is expected from the literature for an activation of this pathway.

Enrichment analysis to annotate the function of WGCNA defined gene modules was conducted using Metascape, a platform used for inclusive gene list annotation and source analysis (https://metascape.org/) (17).

**Predicting PASI scores using gaussian process and ridge regression models**

The end goal of this study was to determine overall the disease endotypes and phenotypes responsible for the progression and severity of Psoriasis. Machine learning (ML) models are being increasingly adopted across the life sciences as decision-making tools. ML models involve general-purpose algorithms that learn patterns from high-dimensional datasets for performing prediction tasks. On the other hand, statistical models (e.g. linear regression) are often more suitable for inference (i.e. distinguishing whether one or more variables are signals or noise) (18). Most machine learning algorithms involve supervised tasks i.e. mapping one or more feature inputs to corresponding labels (i.e. the ground truth) and making predictions for similar but unseen labels. A neural network is a popular machine learning choice for supervised tasks but requires thousands of labelled examples for learning. Given the size of this study's dataset (n 139 patients and 339 time-points) a Gaussian process regression (GRP) was explored (19). Gaussian processes (GPs), a family of Bayesian models, have shown to perform well on a range of modelling tasks given a limited amount of data (46, 48, 77, 78).

Two key features allow GPs to model a range of problems with limited data. Firstly, in the absence of testing data, GPs provide measures of uncertainty to determine how close predictions are to examples in the training dataset(19). Secondly, GPs attempt to model a distribution over functions f(x). This is specified using a kernel covariance function that makes some basic prior assumptions about the relationship such as whether the functions are smooth, linear or rough. The kernel covariance function also makes the basic assumption that data inputs that are closely related are more likely to have similar labels. When the relationship is unknown, a popular choice of kernel is the non-linear Matern 5/2 kernel, which was adopted in this study (19). Further, the kernel function can be decomposed into low-order functions that can be used to model feature inputs additively (24). Many relationships can be decomposed into additive parts, for instance, the price of a building can be broken down into the individual building materials. If the relationship depends jointly on additive low-order interactions, the sum of kernels can be used to model the relationship. If not, the kernels will still specify a suitable model. In this study an additive non-linear kernel function and gaussian process model was implemented using the GPflow python package (version 2.5.2) (25).

During training, the kernel hyperparameters including the length scale ı and variance $\sigma^2$ were tuned for each feature input by maximising the probability of observing the data points, known as the marginal likelihood, on an independent 10-fold validation dataset. Predictions were then obtained from the trained GPR regression.

As a baseline comparison model, a linear ridge regression model was chosen. This model is a special case of a linear regression model that includes a L2 regularisation function to reduce overfitting. A range of hyperparameters were chosen to tune the L2 regularisation parameter on an independent 10-fold validation dataset.

Predictions from the trained models were assessed using shuffled 10-fold held-out testing patient datasets. Performance metrics such as the coefficient of determination ($R^2$) and mean absolute error (MAE) were calculated for each shuffled dataset.

A total of 4 GPR regression and 4 ridge regression models were trained and tested for predicting

Psoriasis Area Severity Index (PASI) scores using several features (i.e. Demographics + clinical features, Skin factors and Skin RNA modules). Notably, to reduce the influence of extreme outliers and allow more direct comparison between feature inputs, models adopted log transformed PASI scores and feature inputs normalised using the robust scaler technique. The robust scaler technique uses the feature median and interquartile range rather than the mean and standard deviation.

To determine the features driving the model relationships, SHAP (SHapley Additive exPlanations) method (26) was adopted. SHAP method is a popular and model-agnostic approach for explaining machine learning outputs. SHAP assesses the impact of each feature on the predicted output while keeping other features unchanged. Higher SHAP values indicate features causing a higher change in the predicted output value and lower SHAP values indicate the opposite.

**PSORT Consortium Membership**

*Executive committee*

Jonathan Barker (St. Johns Institute of Dermatology, Kings College London, London, UK)

Michael Barnes (Centre for Translational Bioinformatics, William Harvey Research Institute, Queen Mary University of London, Charterhouse Square, London, UK)

Paola Di Meglio (St. Johns Institute of Dermatology, Kings College London, London, UK)

Richard Emsley (Institute of Psychiatry, Psychology & Neuroscience, Kings College London, London, UK)

Chris Griffiths (The Manchester Centre for Dermatology Research, The University of Manchester; Manchester, UK)

Nick Reynolds (Institute of Translational and Clinical Medicine, Faculty of Medical Sciences, Newcastle University, Newcastle upon Tyne, UK)

Catherine Smith (St. Johns Institute of Dermatology, Kings College London, London, UK)

Richard Warren (The Manchester Centre for Dermatology Research, The University of Manchester; Manchester, UK)

*Recruiting Principal Investigators (not part of executive committee)*

Dr Anna Chapman (Lewisham & Greenwich Trust)

Dr Rob Ellis (South Tees Hospitals NHS Foundation Trust)

Dr Abigail Fogo (Kingston Hospital NHS Foundation Trust)

Dr Bronwyn Hughes (Portsmouth Hospitals University NHS Trust)

Dr Evmorfia Ladoyanni (Dudley Group NHS Foundation Trust)

Dr Philip Laws (Leeds Teaching Hospitals NHS Trust)

Dr Richard Parslew (Liverpool University Hospitals NHS Foundation Trust)

Dr Gayathri Perera (Chelsea and Westminister Hospital NHS Foundation Trust)

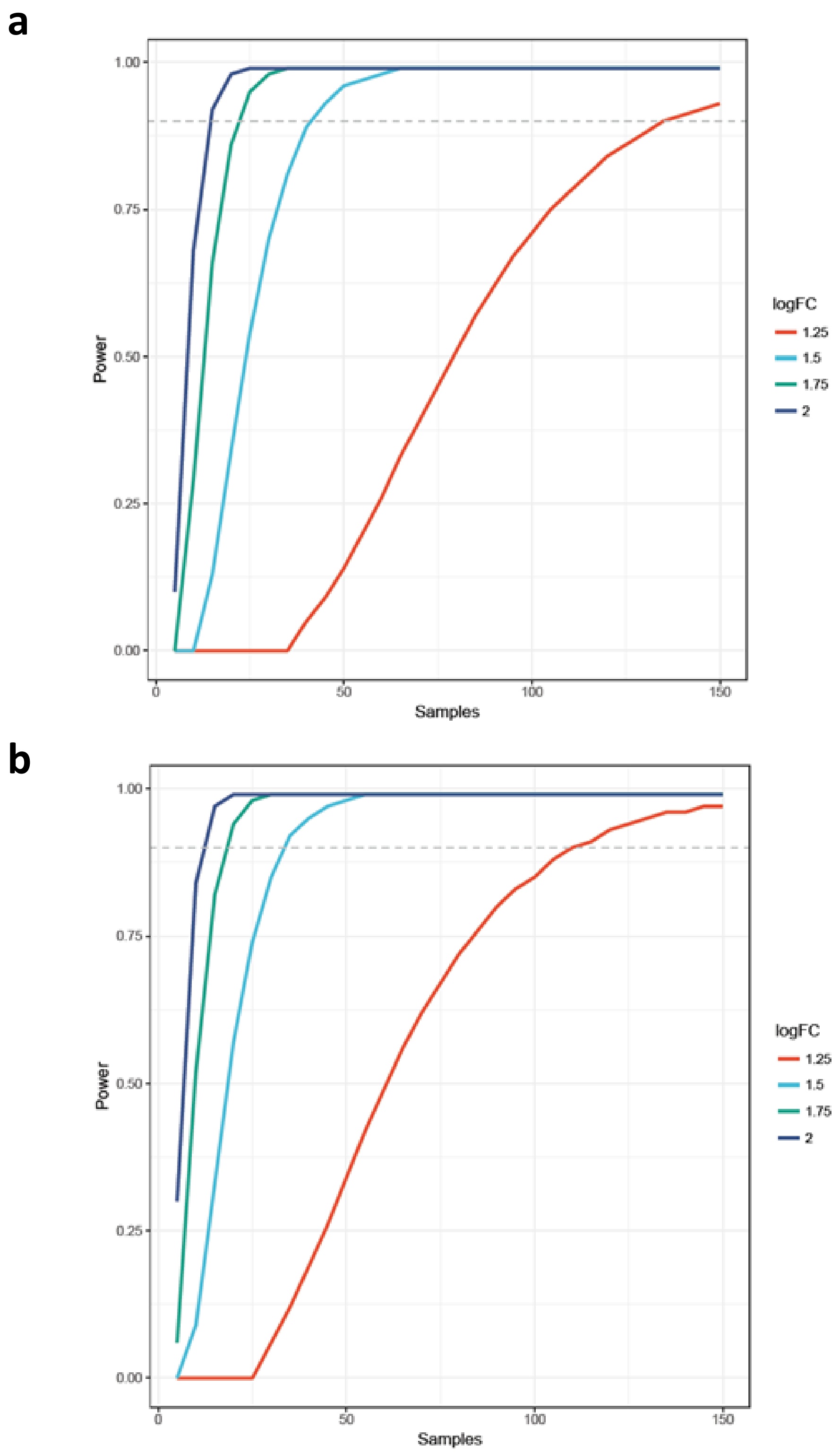


**Supplementary Figure 1. Statistical power calculations for differential expression analysis.**
Power curves projected across an expected range of fold changes based on the assumption that **(a)** 1% or **(b)** 5% of genes are likely to prove prognostic. Power calculations were performed using the method of Guo et al (2014) and parameters derived from our pilot study (Foulkes et al 2018); we calculated the requisite sample size to achieve 90% power (grey dotted line on plot) to detect differential expression associated with response.

**b**

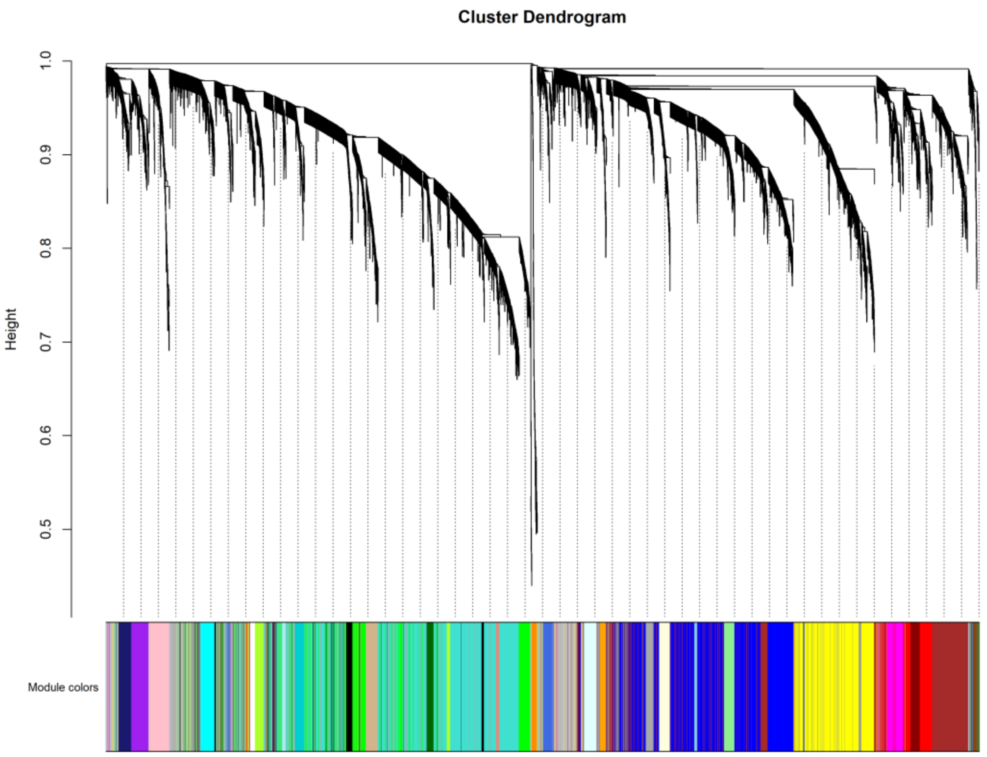
Cluster Dendrogram
Height
1.0
0.9
0.8
0.7
0.6
0.5
Module colors

**b**

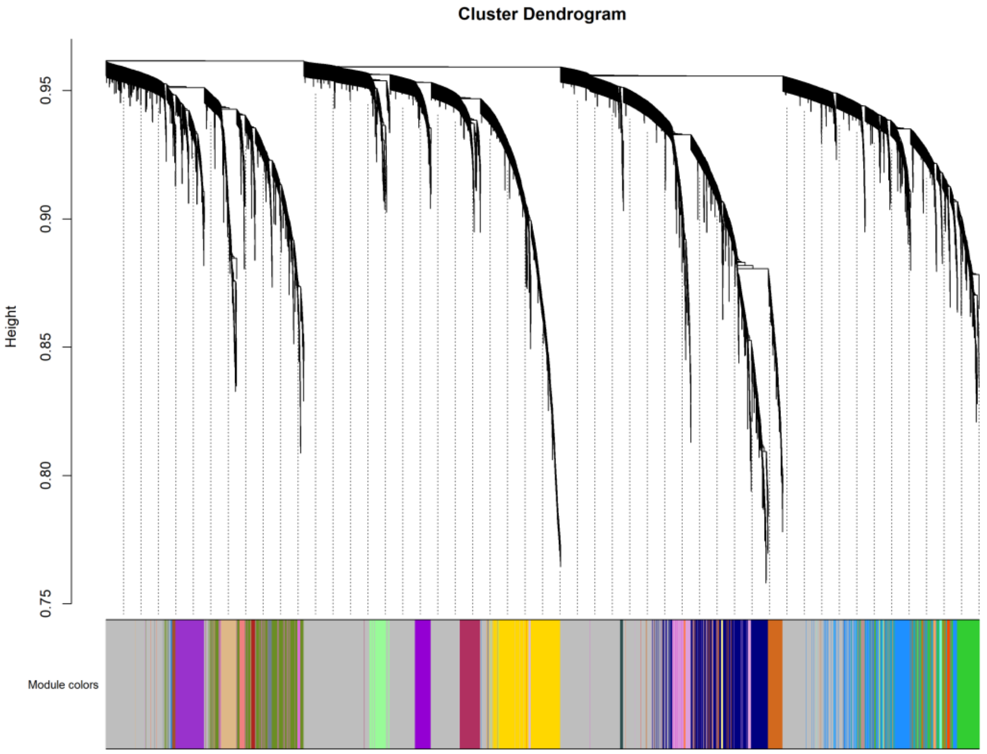
Cluster Dendrogram
Height
0.95
0.90
0.85
0.80
0.75
Module colors

**Supplementary Figure 2. Grouping of genes in skin and blood into co-expressed gene modules using WGCNA.**
WGCNA, a dimensionality reduction method for omics data, was used to group the skin and blood RNA-Seq data into co-expressed gene modules. Briefly, WGCNA works by calculating the correlation between each pair of genes in an expression matrix; the resulting correlation matrix, after some transformations, is then clustered, allowing genes with similar expression profiles to be grouped into modules. Further details are available in the supplementary methods. **(a)** shows hierarchical clustering of genes in the skin data, annotated by colour-coded module assignment and **(b)** shows the same for blood. WGCNA identified 34 modules in skin and 26 modules in blood.
*Abbreviations:* WGCNA, weighted gene correlation network analysis.

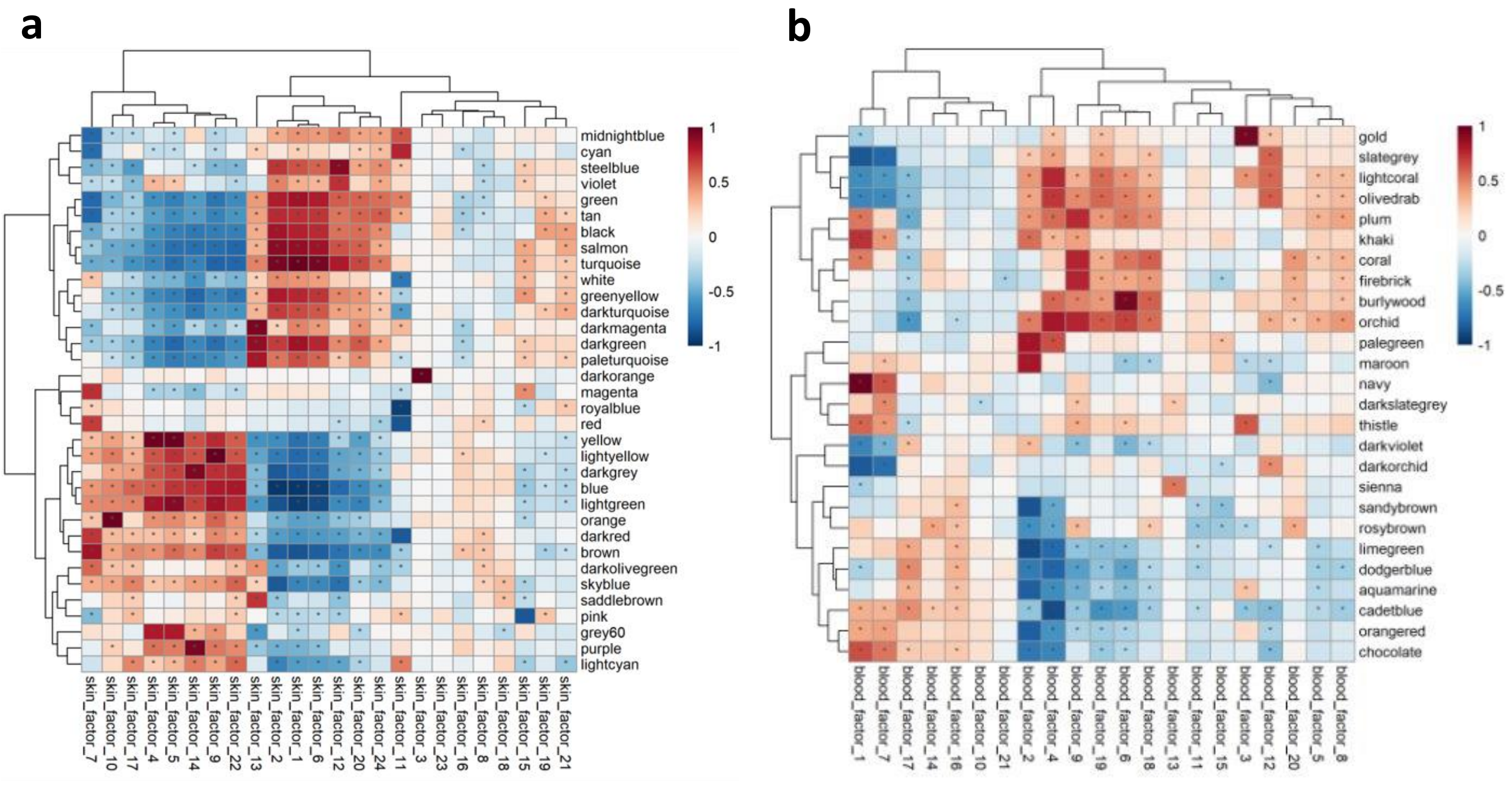


**Supplementary figure 3. WGCNA and ICA identify similar gene expression signatures.**
Correlations between module eigengenes (y-axis) and latent factor values (x-axis) in **(a)** skin and **(b)** blood. The high degree of positive correlation shown in these heatmaps demonstrates that WGCNA and ICA converged on similar gene expression profiles, providing cross-validation of these dimensionality reduction methods.

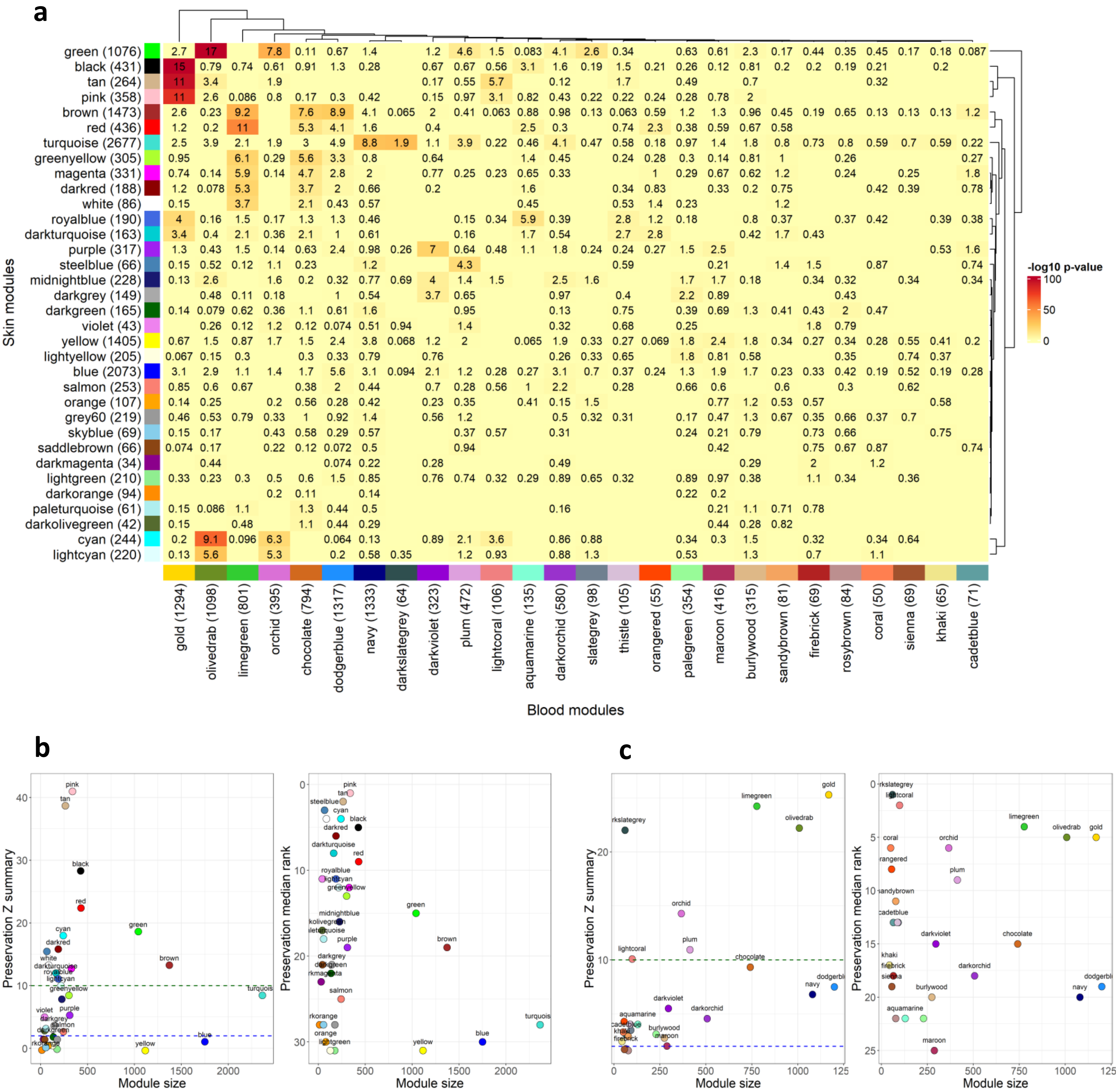
a
Skin modules
Blood modules
-log10 p-value
green (1076)
black (431)
tan (264)
pink (358)
brown (1473)
red (436)
turquoise (2677)
greenyellow (305)
magenta (331)
darkred (188)
white (86)
royalblue (190)
darkturquoise (163)
purple (317)
steelblue (66)
midnightblue (228)
darkgrey (149)
darkgreen (165)
violet (43)
yellow (1405)
lightyellow (205)
blue (2073)
salmon (253)
orange (107)
grey60 (219)
skyblue (69)
saddlebrown (66)
darkmagenta (34)
lightgreen (210)
darkorange (94)
paleturquoise (61)
darkolivegreen (42)
cyan (244)
lightcyan (220)
gold (1294)
olivedrab (1098)
limegreen (801)
orchid (395)
chocolate (794)
dodgerblue (1317)
navy (1333)
darkslategrey (64)
darkviolet (323)
plum (472)
lightcoral (106)
aquamarine (135)
darkorchid (580)
slategrey (98)
thistle (105)
orangered (55)
palegreen (354)
maroon (416)
burlywood (315)
sandybrown (81)
firebrick (69)
rosybrown (84)
coral (50)
sienna (69)
khaki (65)
cadetblue (71)
b
Preservation Z summary
Preservation median rank
Module size
c
Preservation Z summary
Preservation median rank
Module size

**Supplementary Figure 4. Module preservation analysis identifies modules with similar patterns of co-expression in skin and blood, and vice versa.**

WGCNA was used to derive co-expressed gene modules separately in skin and blood; we then sought to identify which modules in skin showed similar patterns of co-expression in blood, and vice versa. We first looked for pairs of modules between skin and blood with significant overlap in gene contents. **(a)** shows gene overlap between the skin and blood modules. The heatmap is coloured by –log10 p-values from right tailed fisher's exact test and the numbers in each cell correspond to the percentage of the union of genes in each module pair that overlap. We also used tested module preservation between the skin and blood modules using functionality available in the WGCNA R package. We derived two composite module preservation scores, preservation Z summary and median rank. **(b)** shows composite scores from testing preservation of the skin modules in blood and **(c)** shows the scores derived from testing preservation of the blood modules in skin. In each plot the left panel shows preservation Z summary and the right panel shows median rank. Each score is plotted against module size. The blue horizontal line indicates moderate evidence for preservation and the green horizontal line indicates strong evidence for preservation. The black , pink, and tan modules in skin exhibited strong evidence for preservation in blood and were found to have the most significant gene overlap with the gold module in blood. The green skin module was also strongly preserved in blood and exhibited significant gene overlap with the olivedrab module in blood.

*Abbreviations:* WGCNA, weighted gene correlation network analysis.

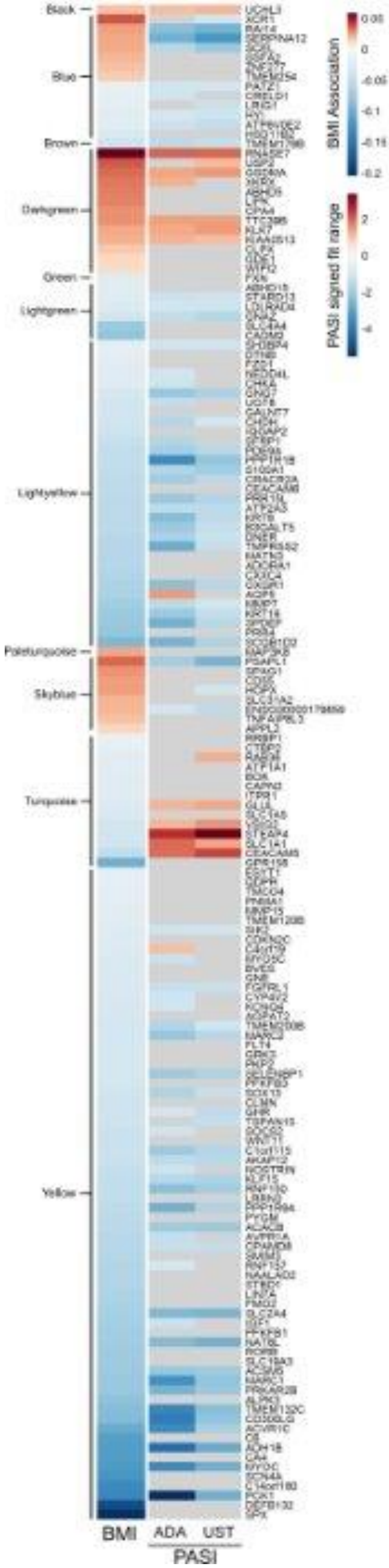
Black
Blue
Brown
Darkgreen
Green
Lightgreen
Lightyellow
Paleturquoise
Skyblue
Turquoise
Yellow
BMI
ADA
UST
PASI
BMI Association
PASI signed fit range

**Supplementary figure 5. A transcriptome signature associated with BMI in non-lesional skin (NLS) is associated with disease severity in lesional skin.**

This heatmap shows genes that were assigned to modules by WGCNA and exhibited a significant association with BMI in NLS based on differential expression analysis; their associations with BMI in NLS and PASI in LS in each drug cohort are shown. Red indicates a positive association, blue indicates a negative association, and grey indicates a non-significant association. Of note, genes in the lightyellow module were negatively associated with BMI in NLS and negatively associated with PASI in LS in both drug cohorts.

*Abbreviations:* BMI, body mass index; WGCNA, weighted gene correlation network analysis; NLS, non-lesional skin; PASI, psoriasis area and severity index; LS, lesional skin.

a

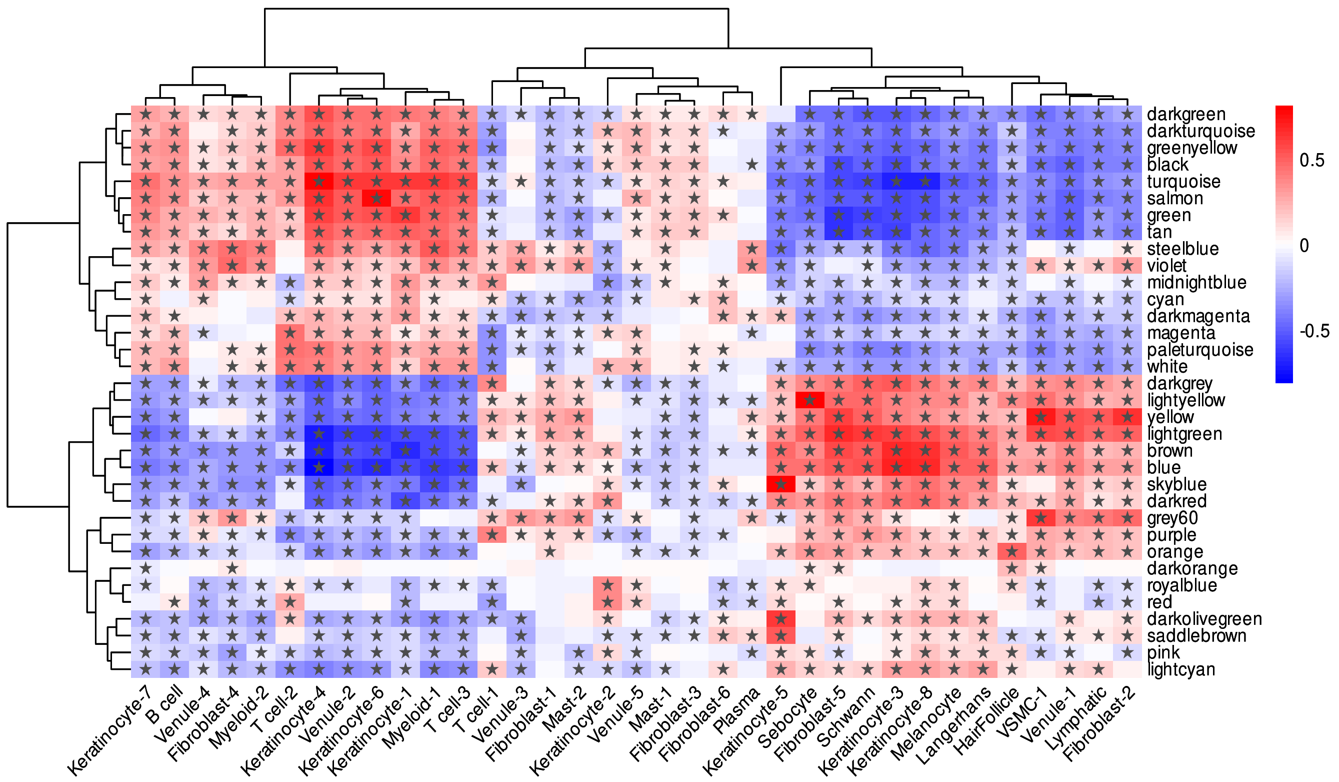
darkgreen
darkturquoise
greenyellow
black
turquoise
salmon
green
tan
steelblue
violet
midnightblue
cyan
darkmagenta
magenta
paleturquoise
white
darkgrey
lightyellow
yellow
lightgreen
brown
blue
skyblue
darkred
grey60
purple
orange
darkorange
royalblue
red
darkolivegreen
saddlebrown
pink
lightcyan
0.5
0
-0.5
Keratinocyte-7
B cell
Venule-4
Fibroblast-4
Myeloid-2
T cell-2
Keratinocyte-4
Venule-2
Keratinocyte-6
Keratinocyte-1
Myeloid-1
T cell-3
T cell-1
Venule-3
Fibroblast-1
Mast-2
Keratinocyte-2
Venule-5
Mast-1
Fibroblast-3
Fibroblast-6
Plasma
Keratinocyte-5
Sebocyte
Fibroblast-5
Schwann
Keratinocyte-3
Keratinocyte-8
Melanocyte
Langerhans
HairFollicle
VSMC-1
Venule-1
Lymphatic
Fibroblast-2

b

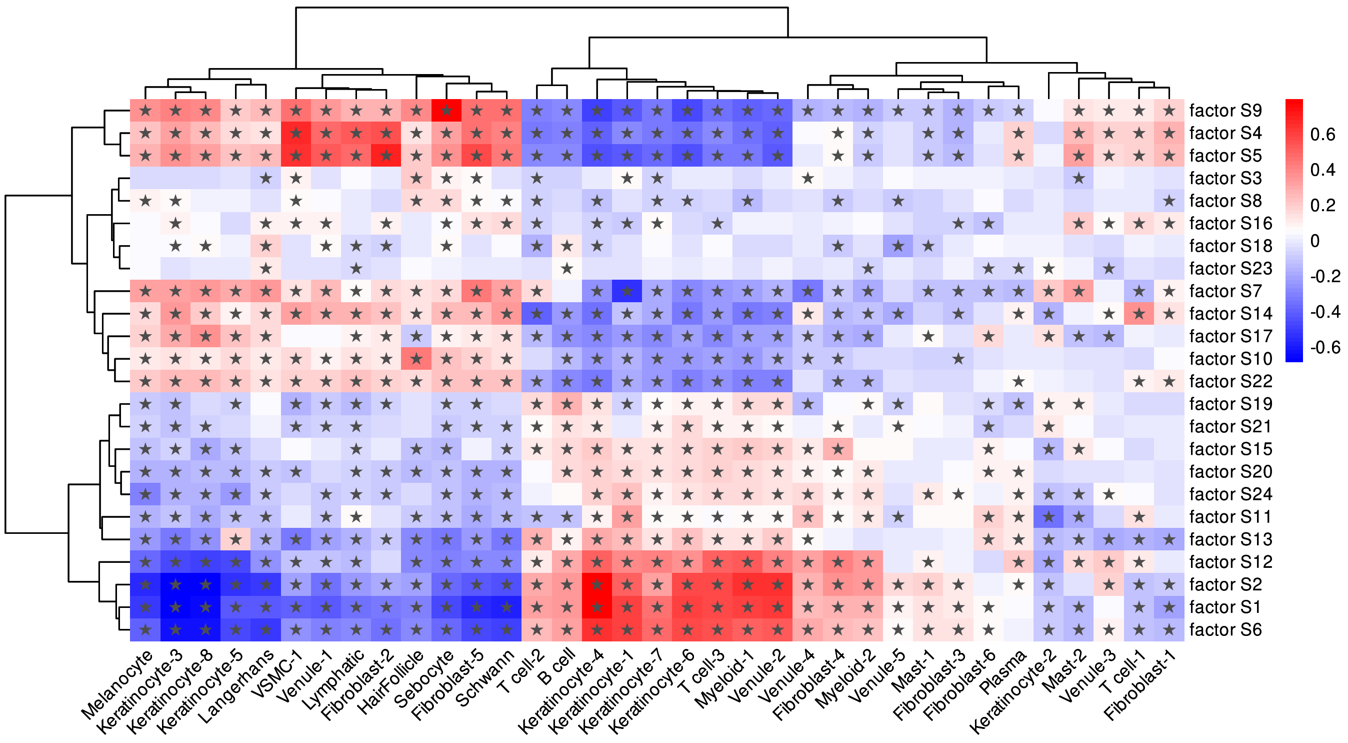
factor S9
factor S4
factor S5
factor S3
factor S8
factor S16
factor S18
factor S23
factor S7
factor S14
factor S17
factor S10
factor S22
factor S19
factor S21
factor S15
factor S20
factor S24
factor S11
factor S13
factor S12
factor S2
factor S1
factor S6
0.6
0.4
0.2
0
-0.2
-0.4
-0.6
Melanocyte
Keratinocyte-3
Keratinocyte-8
Keratinocyte-5
Langerhans
VSMC-1
Venule-1
Lymphatic
Fibroblast-2
HairFollicle
Sebocyte
Fibroblast-5
Schwann
T cell-2
B cell
Keratinocyte-4
Keratinocyte-1
Keratinocyte-7
Keratinocyte-6
T cell-3
Myeloid-1
Venule-2
Venule-4
Fibroblast-4
Myeloid-2
Venule-5
Mast-1
Fibroblast-3
Fibroblast-6
Plasma
Keratinocyte-2
Mast-2
Venule-3
T cell-1
Fibroblast-1

**Supplementary Figure 6. Cell type deconvolution of the gene modules and latent factors in skin.**
We derived predicted cell-type fractions for the skin RNA-Seq data (discovery and replication cohorts combined) using CibersortX and a skin single cell transcriptome atlas published by Hughes et al (2020), publicly available on the Single Cell Portal (https://singlecell.broadinstitute.org/single_cell). We then correlated the predicted cell-type fractions with **(a)** module eigengenes, derived by WGCNA, and **(b)** latent factor values, derived by ICA. Asterisks (*) indicate statistical significance (adjusted p-value<0.05). *Abbreviations:* WGCNA, weighted gene correlation network analysis; ICA, independent component analysis.

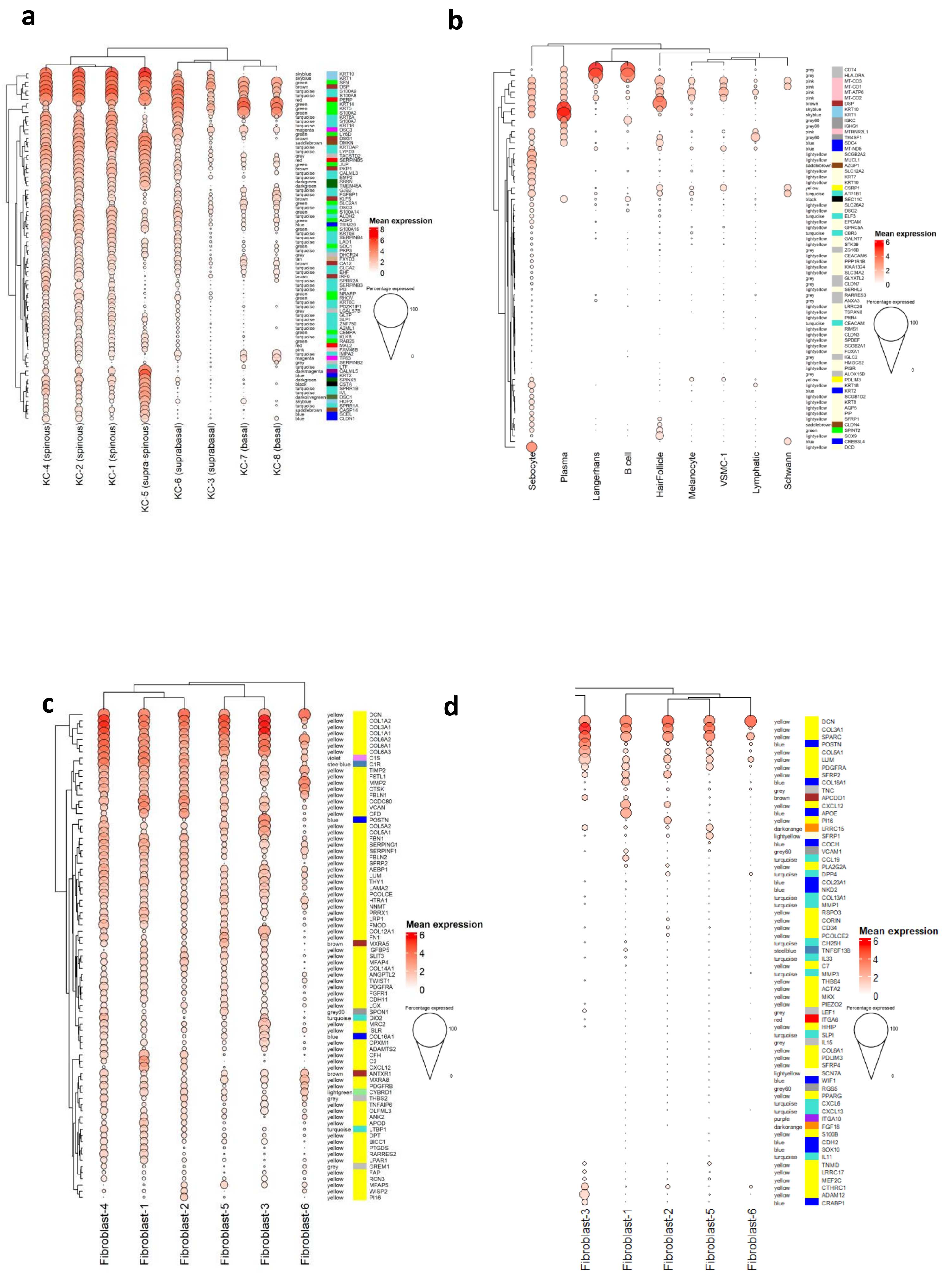
a
KC-4 (spinous)
KC-2 (spinous)
KC-1 (spinous)
KC-5 (supra-spinous)
KC-6 (suprabasal)
KC-3 (suprabasal)
KC-7 (basal)
KC-8 (basal)
skyblue KRT10
skyblue KRT1
green SFN
brown DSP
turquoise S100A9
turquoise S100A8
red PERP
green KRT14
green KRT5
green S100A2
turquoise KRT6A
turquoise S100A7
turquoise KRT16
magenta DSC3
green LY6D
brown DSG1
saddlebrown DMKN
turquoise KRTDAP
turquoise LYPD3
grey TACSTD2
red SERPINB5
green JUP
brown PKP1
turquoise CALML3
turquoise EMP2
darkgreen SBSN
darkgreen TMEM45A
turquoise GJB2
turquoise FGFBP1
brown KLF5
green SLC2A1
turquoise DSG3
green S100A14
turquoise ALDH2
green AQP3
blue TRIM29
green S100A16
turquoise KRT6B
turquoise SERPINB4
turquoise LAD1
green SDC1
turquoise PKP3
grey DHCR24
tan FXYD3
brown CA12
turquoise CLCA2
turquoise EHF
brown IRF6
turquoise SPRR2A
turquoise SERPINB3
turquoise PI3
green NRARP
green RHOV
turquoise KRT6C
turquoise PDZK1IP1
grey LGALS7B
turquoise GLTP
turquoise SLPI
turquoise ZNF750
turquoise A2ML1
green CEBPA
turquoise KLK8
green RAB25
red MAL2
pink FAM46B
turquoise IMPA2
magenta TP63
grey SERPINB2
turquoise LTF
darkmagenta CALML5
blue KRT2
darkgreen SPINK5
black CSTA
turquoise SPRR1B
turquoise IVL
darkolivegreen DSC1
skyblue HOPX
turquoise SPRR1A
saddlebrown CASP14
blue SCEL
blue CLDN1
Mean expression
8
6
4
2
0
Percentage expressed
100
0
b
Sebocyte
Plasma
Langerhans
B cell
HairFollicle
Melanocyte
VSMC-1
Lymphatic
Schwann
grey CD74
grey HLA-DRA
pink MT-CO3
pink MT-CO1
pink MT-ATP6
pink MT-CO2
brown DSP
skyblue KRT10
skyblue KRT1
grey60 IGKC
grey60 IGHG1
pink MTRNR2L1
grey60 TM4SF1
blue SDC4
blue MT-ND5
lightyellow SCGB2A2
lightyellow MUCL1
saddlebrown AZGP1
lightyellow SLC12A2
lightyellow KRT7
lightyellow KRT19
yellow CSRP1
turquoise ATP1B1
black SEC11C
lightyellow SLC26A2
lightyellow DSG2
turquoise ELF3
lightyellow EPCAM
lightyellow GPRC5A
turquoise CBR3
lightyellow GALNT7
lightyellow STK39
grey ZG16B
lightyellow CEACAM6
lightyellow PPP1R1B
lightyellow KIAA1324
lightyellow SLC34A2
grey GLYATL2
grey CLDN7
lightyellow SERHL2
grey RARRES3
grey ANXA3
lightyellow LRRC26
lightyellow TSPAN8
lightyellow PRR4
turquoise CEACAM5
lightyellow RIMS1
lightyellow CLDN3
lightyellow SPDEF
lightyellow SCGB2A1
lightyellow FOXA1
grey IGLC2
lightyellow HMGCS2
lightyellow PIGR
grey ALOX15B
yellow PDLIM3
lightyellow KRT18
blue KRT2
lightyellow SCGB1D2
lightyellow KRT8
lightyellow AQP5
lightyellow PIP
lightyellow SFRP1
saddlebrown CLDN4
green SPINT2
lightyellow SOX9
blue CREB3L4
lightyellow DCD
Mean expression
6
4
2
0
Percentage expressed
100
0
c
Fibroblast-4
Fibroblast-1
Fibroblast-2
Fibroblast-5
Fibroblast-3
Fibroblast-6
yellow DCN
yellow COL1A2
yellow COL3A1
yellow COL1A1
yellow COL6A2
yellow COL6A1
yellow COL6A3
violet C1S
steelblue C1R
yellow TIMP2
yellow FSTL1
yellow MMP2
yellow CTSK
yellow FBLN1
yellow CCDC80
yellow VCAN
yellow CFD
blue POSTN
yellow COL5A2
yellow COL5A1
yellow FBN1
yellow SERPING1
yellow SERPINF1
yellow FBLN2
yellow SFRP2
yellow AEBP1
yellow LUM
yellow THY1
yellow LAMA2
yellow PCOLCE
yellow HTRA1
yellow NNMT
yellow PRRX1
yellow LRP1
yellow FMOD
yellow COL12A1
yellow FN1
brown MXRA5
yellow IGFBP5
yellow SLIT3
yellow MFAP4
yellow COL14A1
yellow ANGPTL2
yellow TWIST1
yellow PDGFRA
yellow FGFR1
yellow CDH11
yellow LOX
grey60 SPON1
turquoise DIO2
yellow MRC2
yellow ISLR
blue COL16A1
yellow CPXM1
yellow ADAMTS2
yellow CFH
yellow C3
yellow CXCL12
brown ANTXR1
yellow MXRA8
yellow PDGFRB
lightgreen CYBRD1
grey THBS2
yellow TNFAIP6
yellow OLFML3
yellow ANK2
yellow APOD
turquoise LTBP1
yellow DPT
yellow BICC1
yellow PTGDS
yellow RARRES2
yellow LPAR1
grey GREM1
yellow FAP
yellow RCN3
yellow MFAP5
yellow WISP2
yellow PI16
Mean expression
6
4
2
0
Percentage expressed
100
0
d
Fibroblast-3
Fibroblast-1
Fibroblast-2
Fibroblast-5
Fibroblast-6
yellow DCN
yellow COL3A1
yellow SPARC
blue POSTN
yellow COL5A1
yellow LUM
yellow PDGFRA
yellow SFRP2
blue COL18A1
grey TNC
brown APCDD1
yellow CXCL12
blue APOE
yellow PI16
darkorange LRRC15
lightyellow SFRP1
blue COCH
grey60 VCAM1
turquoise CCL19
yellow PLA2G2A
turquoise DPP4
blue COL23A1
blue NKD2
turquoise COL13A1
turquoise MMP1
yellow RSPO3
yellow CORIN
yellow CD34
yellow PCOLCE2
turquoise CH25H
steelblue TNFSF13B
turquoise IL33
yellow C7
turquoise MMP3
yellow THBS4
yellow ACTA2
yellow MKX
yellow PIEZO2
grey LEF1
red ITGA6
yellow HHIP
turquoise SLPI
grey IL15
yellow COL8A1
yellow PDLIM3
yellow SFRP4
lightyellow SCN7A
blue WIF1
grey60 RGS5
yellow PPARG
turquoise CXCL6
turquoise CXCL13
purple ITGA10
darkorange FGF16
yellow S100B
blue CDH2
blue SOX10
turquoise IL11
yellow TNMD
yellow LRRC17
yellow MEF2C
yellow CTHRC1
yellow ADAM12
blue CRABP1
Mean expression
6
4
2
0
Percentage expressed
100
0

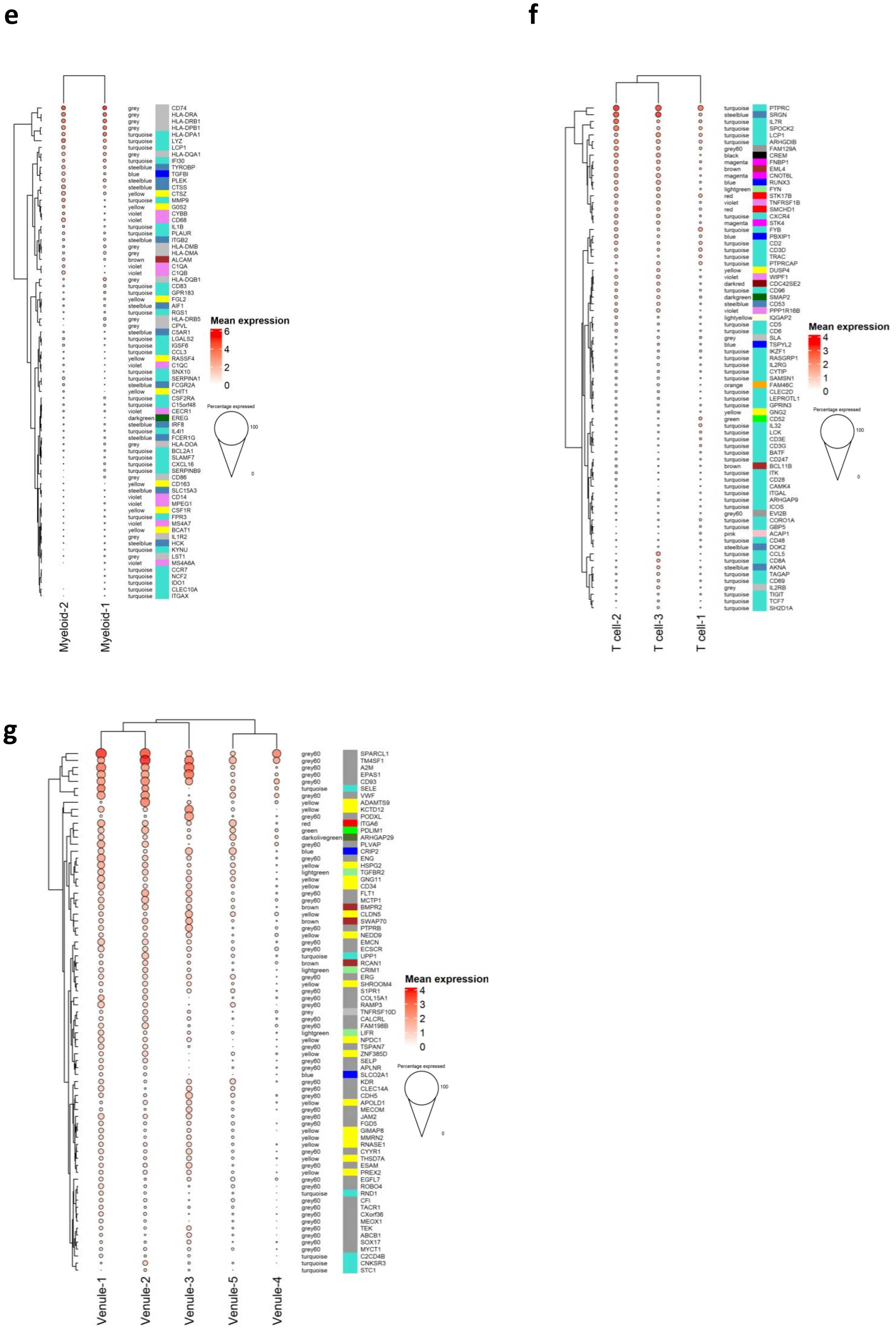
e
grey CD74
grey HLA-DRA
grey HLA-DRB1
grey HLA-DPB1
turquoise HLA-DPA1
turquoise LYZ
turquoise LCP1
grey HLA-DQA1
turquoise IFI30
steelblue TYROBP
blue TGFBI
steelblue PLEK
steelblue CTSS
yellow CTSZ
turquoise MMP9
yellow G0S2
violet CYBB
violet CD68
turquoise IL1B
turquoise PLAUR
steelblue ITGB2
grey HLA-DMB
grey HLA-DMA
brown ALCAM
violet C1QA
violet C1QB
grey HLA-DQB1
turquoise CD83
turquoise GPR183
yellow FGL2
steelblue AIF1
turquoise RGS1
grey HLA-DRB5
grey CPVL
steelblue C5AR1
turquoise LGALS2
turquoise IGSF6
turquoise CCL3
yellow RASSF4
violet C1QC
turquoise SNX10
turquoise SERPINA1
steelblue FCGR2A
yellow CHIT1
turquoise CSF2RA
turquoise C15orf48
violet CECR1
darkgreen EREG
steelblue IRF8
turquoise IL4I1
steelblue FCER1G
grey HLA-DOA
turquoise BCL2A1
turquoise SLAMF7
turquoise CXCL16
turquoise SERPINB9
grey CD86
yellow CD163
steelblue SLC15A3
violet CD14
violet MPEG1
yellow CSF1R
turquoise FPR3
violet MS4A7
yellow BCAT1
grey IL1R2
steelblue HCK
turquoise KYNU
grey LST1
violet MS4A6A
turquoise CCR7
turquoise NCF2
turquoise IDO1
turquoise CLEC10A
turquoise ITGAX
Mean expression
6
4
2
0
Percentage expressed
100
0
Myeloid-2
Myeloid-1
f
turquoise PTPRC
steelblue SRGN
turquoise IL7R
turquoise SPOCK2
turquoise LCP1
turquoise ARHGDIB
grey60 FAM129A
black CREM
magenta FNBP1
brown EML4
magenta CNOT6L
blue RUNX3
lightgreen FYN
red STK17B
violet TNFRSF1B
red SMCHD1
turquoise CXCR4
magenta STK4
turquoise FYB
blue PBXIP1
turquoise CD2
turquoise CD3D
turquoise TRAC
turquoise PTPRCAP
yellow DUSP4
violet WIPF1
darkred CDC42SE2
turquoise CD96
darkgreen SMAP2
steelblue CD53
violet PPP1R16B
lightyellow IQGAP2
turquoise CD5
turquoise CD6
grey SLA
blue TSPYL2
turquoise IKZF1
turquoise RASGRP1
turquoise IL2RG
turquoise CYTIP
turquoise SAMSN1
orange FAM46C
turquoise CLEC2D
turquoise LEPROTL1
turquoise GPRIN3
yellow GNG2
green CD52
turquoise IL32
turquoise LCK
turquoise CD3E
turquoise CD3G
turquoise BATF
turquoise CD247
brown BCL11B
turquoise ITK
turquoise CD28
turquoise CAMK4
turquoise ITGAL
turquoise ARHGAP9
turquoise ICOS
grey60 EVI2B
turquoise CORO1A
turquoise GBP5
pink ACAP1
turquoise CD48
steelblue DOK2
turquoise CCL5
turquoise CD8A
steelblue AKNA
turquoise TAGAP
turquoise CD69
grey IL2RB
turquoise TIGIT
turquoise TCF7
turquoise SH2D1A
Mean expression
4
3
2
1
0
Percentage expressed
100
0
T cell-2
T cell-3
T cell-1
g
grey60 SPARCL1
grey60 TM4SF1
grey60 A2M
grey60 EPAS1
grey60 CD93
turquoise SELE
grey60 VWF
yellow ADAMTS9
yellow KCTD12
grey60 PODXL
red ITGA6
green PDLIM1
darkolivegreen ARHGAP29
grey60 PLVAP
blue CRIP2
grey60 ENG
yellow HSPG2
lightgreen TGFBR2
yellow GNG11
yellow CD34
grey60 FLT1
grey60 MCTP1
brown BMPR2
yellow CLDN5
brown SWAP70
grey60 PTPRB
yellow NEDD9
grey60 EMCN
grey60 ECSCR
turquoise UPP1
brown RCAN1
lightgreen CRIM1
grey60 ERG
yellow SHROOM4
grey60 S1PR1
grey60 COL15A1
grey60 RAMP3
grey TNFRSF10D
grey60 CALCRL
grey60 FAM198B
lightgreen LIFR
yellow NPDC1
grey60 TSPAN7
yellow ZNF385D
grey60 SELP
grey60 APLNR
blue SLCO2A1
grey60 KDR
grey60 CLEC14A
grey60 CDH5
yellow APOLD1
grey60 MECOM
grey60 JAM2
grey60 FGD5
yellow GIMAP8
yellow MMRN2
yellow RNASE1
grey60 CYYR1
yellow THSD7A
grey60 ESAM
yellow PREX2
grey60 EGFL7
grey60 ROBO4
turquoise RND1
grey60 CFI
grey60 TACR1
grey60 CXorf36
grey60 MEOX1
grey60 TEK
grey60 ABCB1
grey60 SOX17
grey60 MYCT1
turquoise C2CD4B
turquoise CNKSR3
turquoise STC1
Mean expression
4
3
2
1
0
Percentage expressed
100
0
Venule-1
Venule-2
Venule-3
Venule-5
Venule-4

**Supplementary Figure 7. Expression of cell cluster marker genes in a skin single cell transcriptome atlas.** Heatmaps showing the mean expression within specified cell clusters (columns) of the top 80 marker genes (ranked by average log fold change; rows) in **(a)** keratinocytes, **(b)** sebocytes, **(c)** fibroblasts, **(e)** myeloid cells, **(f)** T-cells, and **(g)** venules. This data was derived from a skin single cell transcriptome atlas published by Hughes et al (2020), publicly available on the Single Cell Portal (https://singlecell.broadinstitute.org/single_cell). The row annotation indicates the modules to which these genes were assigned in our dataset. **(d)** Shows the expression in this single cell atlas data of marker genes, identified by Steele et al (2024), which differentiate between different sub-types of fibroblasts.

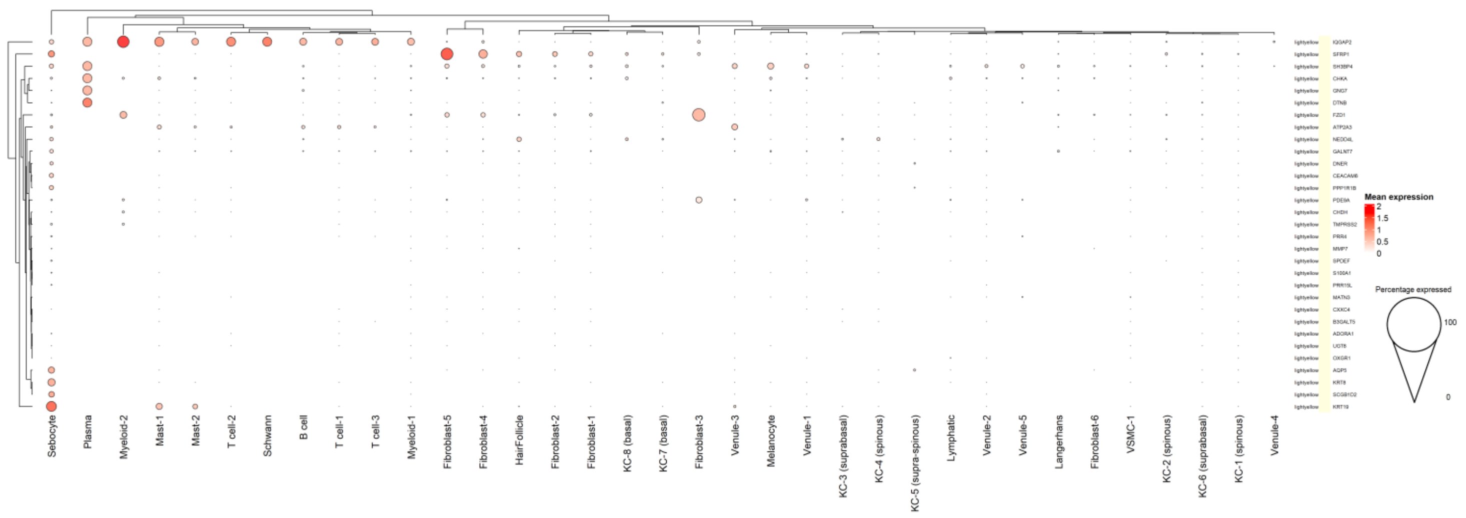


**Supplementary Figure 8. Expression of BMI-associated genes in a skin single cell transcriptome atlas.** Heatmaps showing the mean expression within skin cell clusters (columns) of BMI-associated DEGs that overlap with the lightyellow module. This data was derived from the skin single cell transcriptome atlas published by Hughes et al (2020), publicly available on the Single Cell Portal (https://singlecell.broadinstitute.org/single_cell).

*Abbreviations:* BMI, body mass index.

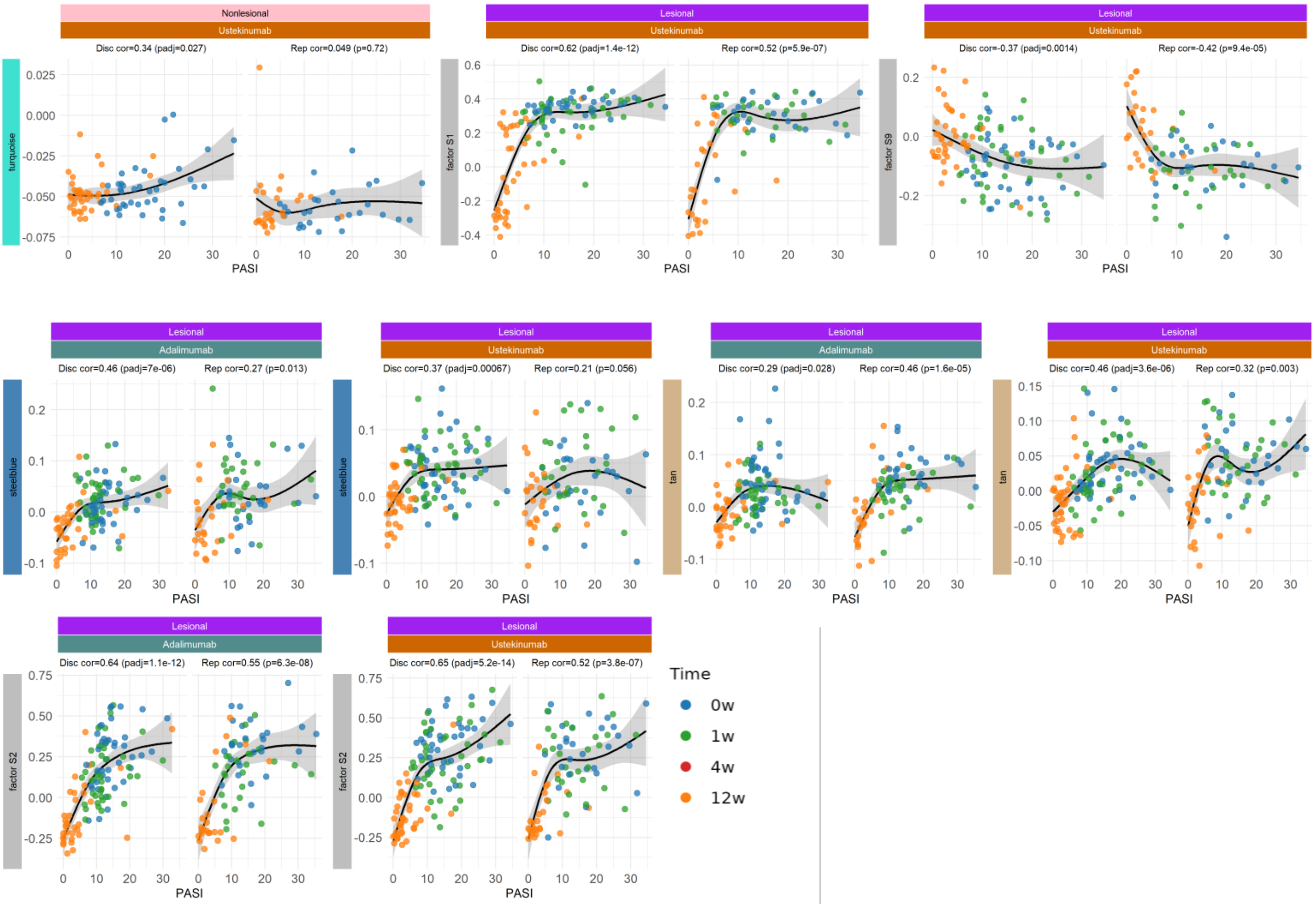


**Supplementary Figure 9. Further exemplar module and latent factor-trait correlation plots.**
Scatter plots of further exemplar WGCNA module and ICA factor trait associations with disease severity (PASI). Each point represents a single sample, is marked with a symbol according to drug cohort and colour-coded according to timepoint (blue = 0w, green = 1w, red = 4w, orange = 12w). The grey shading indicates 95% confidence intervals. The title banner provides information on the tissue and drug cohort. Module membership is denoted opposite the y axis. Curves represent natural-spline fits (3 d.f.) with 95 % confidence bands.
*Abbreviations:* PASI, Psoriasis Area and Severity Index; WGCNA, Weighted gene correlation network analysis; ICA independent component analysis; w, weeks

**a**

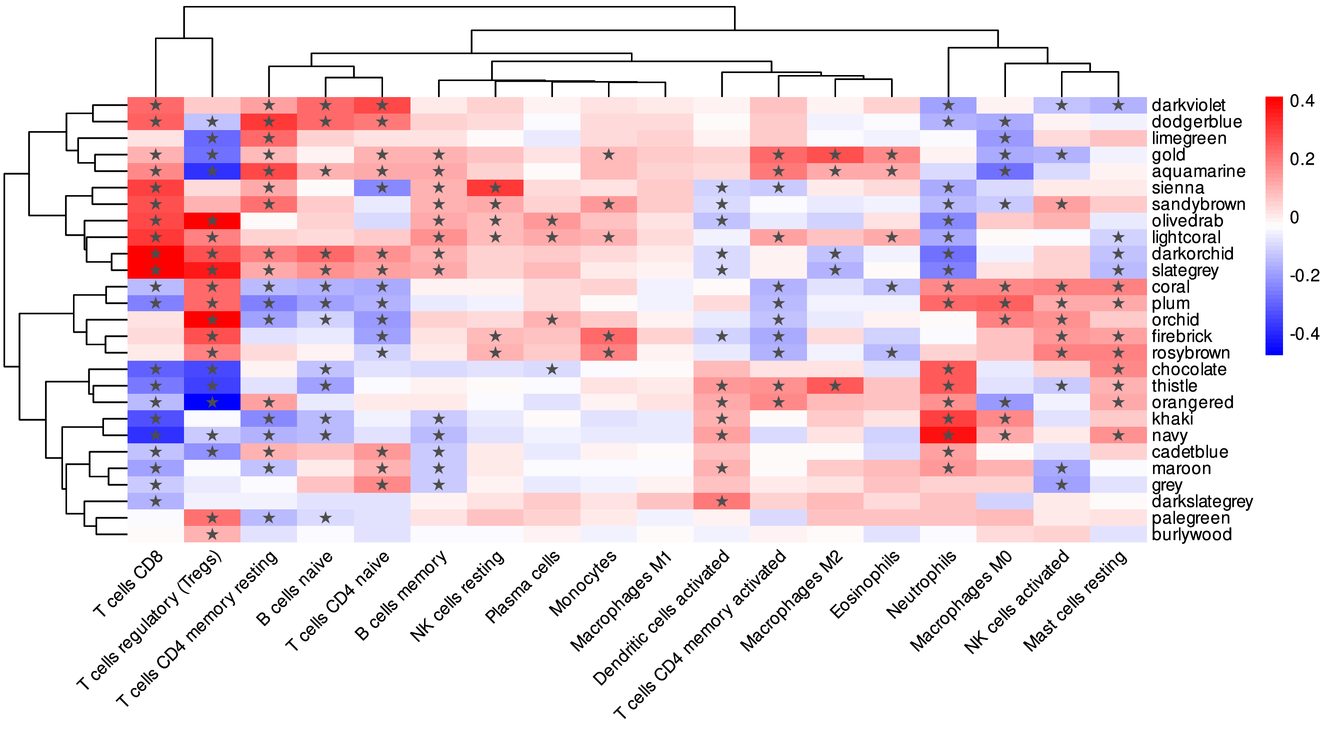

darkviolet
dodgerblue
limegreen
gold
aquamarine
sienna
sandybrown
olivedrab
lightcoral
darkorchid
slategrey
coral
plum
orchid
firebrick
rosybrown
chocolate
thistle
orangered
khaki
navy
cadetblue
maroon
grey
darkslategrey
palegreen
burlywood
0.4
0.2
0
-0.2
-0.4
T cells CD8
T cells regulatory (Tregs)
T cells CD4 memory resting
B cells naive
T cells CD4 naive
B cells memory
NK cells resting
Plasma cells
Monocytes
Macrophages M1
Dendritic cells activated
T cells CD4 memory activated
Macrophages M2
Eosinophils
Neutrophils
Macrophages M0
NK cells activated
Mast cells resting


**b**

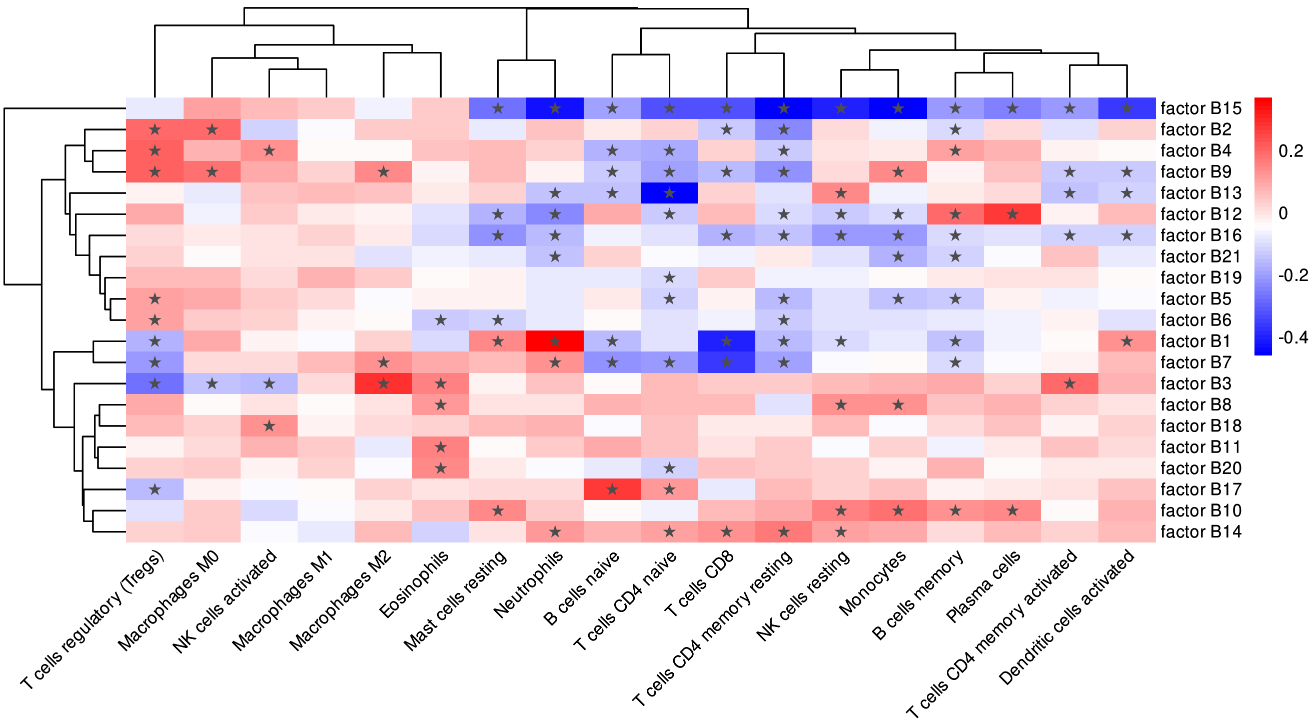

factor B15
factor B2
factor B4
factor B9
factor B13
factor B12
factor B16
factor B21
factor B19
factor B5
factor B6
factor B1
factor B7
factor B3
factor B8
factor B18
factor B11
factor B20
factor B17
factor B10
factor B14
0.2
0
-0.2
-0.4
T cells regulatory (Tregs)
Macrophages M0
NK cells activated
Macrophages M1
Macrophages M2
Eosinophils
Mast cells resting
Neutrophils
B cells naive
T cells CD4 naive
T cells CD8
T cells CD4 memory resting
NK cells resting
Monocytes
B cells memory
Plasma cells
T cells CD4 memory activated
Dendritic cells activated

**Supplementary Figure 10. Cell type deconvolution of the gene modules and latent factors in blood.** We derived predicted cell-type fractions for the blood RNA-Seq data using CibersortX and the LM22 reference dataset. We then correlated the predicted cell-type fractions with **(a)** module eigengenes, derived by WGCNA, and **(b)** latent factor values, derived by ICA. Asterisks (*) indicate statistical significance (adjusted p-value<0.05).
*Abbreviations:* WGCNA, weighted gene correlation network analysis; ICA, independent component analysis.

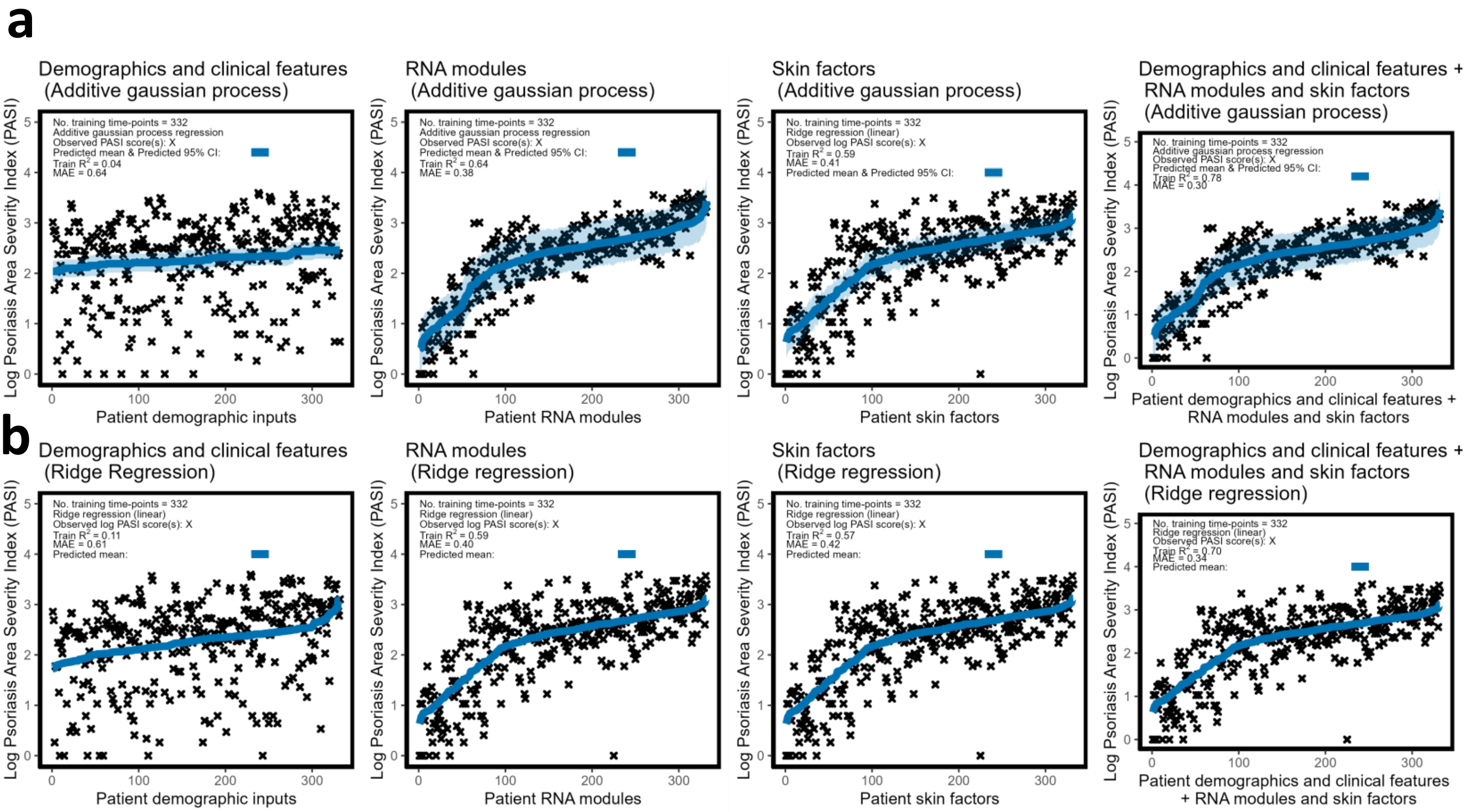


**Supplementary Figure 11**. **Prediction of PASI based on module eigengenes and latent factors in lesional skin and demographic and clinical features using additive gaussian process regression and linear ridge regression.** **(a)** Additive gaussian process regression models depicting the relationship between module eigengenes and latent factor values in lesional skin and demographic and clinical features (discovery and replication cohorts combined) and log scaled PASI. Top row: the first regression model was trained on patient demographic and clinical information. The next two regression models were trained on module eigengenes or latent factors. The third regression model was trained on all features. **(b)** Ridge regression (baseline) models depicting the relationship between patient inputs and PASI. The observed log PASI scores from the training dataset (n=332 time-points) are depicted by the cross symbols. The shaded regions represent the additive gaussian process models predicted 95% confidence intervals.

*Abbreviations:* PASI, psoriasis area and severity index.

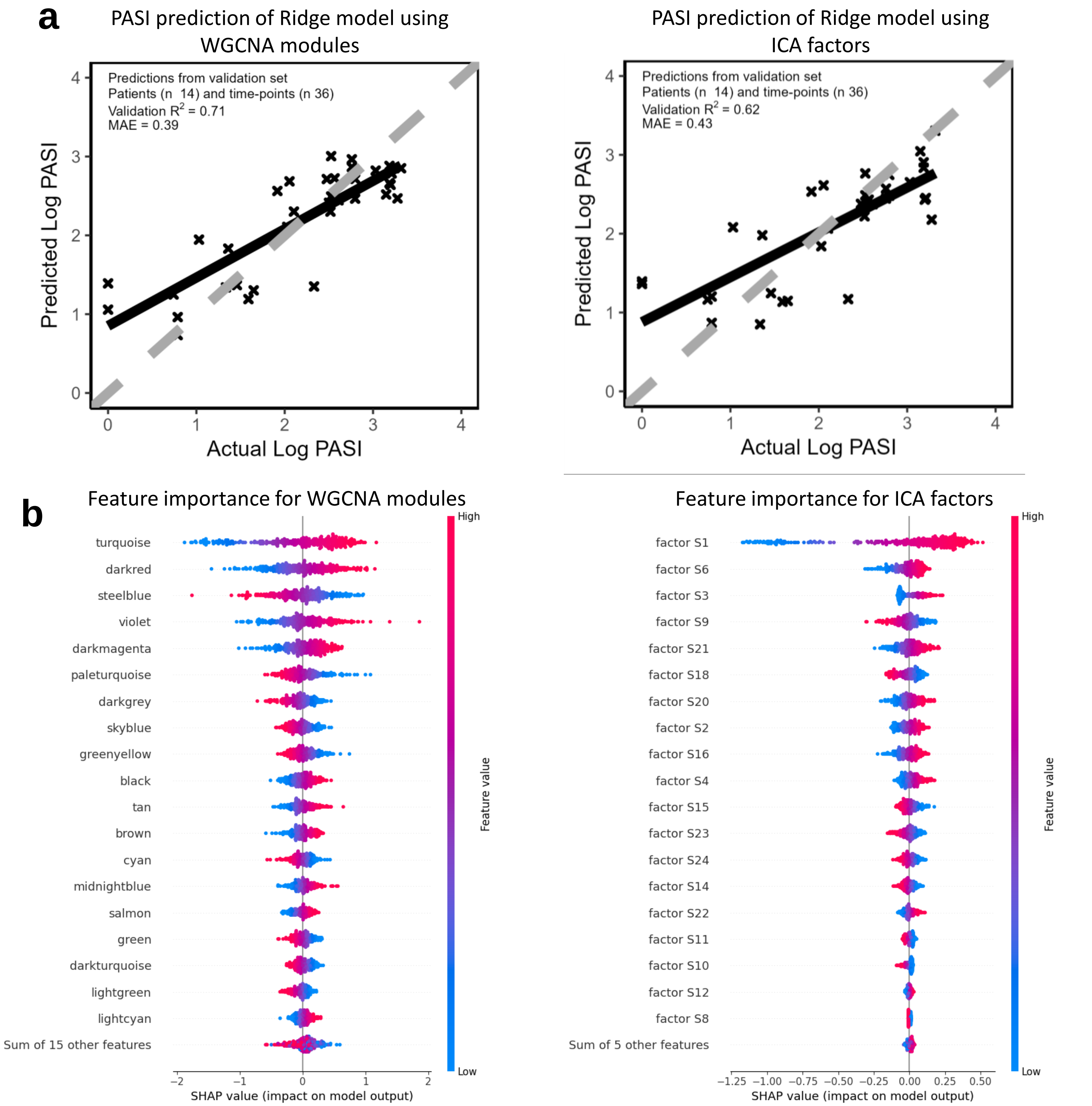


**Supplementary Figure 12. Prediction of PASI based on module eigengenes and latent factors in lesional skin and demographic and clinical features using linear ridge regression.**
**(a)** Linear (ridge) regression models depicting the relationship between module eigengenes and latent factor values in lesional skin and demographic and clinical features (discovery and replication cohorts combined) and log scaled PASI. The performance comparison between the predicted and actual log PASI scores were optimized on the validation dataset. Among all features, modules eigengenes and latent factors gave the best performance. Demographic and clinical features made no difference to performance (Supplementary Fig. 12). The observed log PASI scores from the training dataset (n=332 time-points) are depicted by the cross symbols. **(b)** Scatter plots show the relative feature importance determined using the SHAP (SHapley Additive exPlanations) method. SHAP values were calculated using a model-agnostic approach, where model features are independently altered and the resulting change in the predicted output is recorded while keeping other features unchanged. SHAP values on the right side of the x-axis indicate features decreasing PASI, whilst those shifting right indicate the opposite. *Abbreviations:* PASI, psoriasis area and severity index; SHAP, SHapley Additive exPlanations.

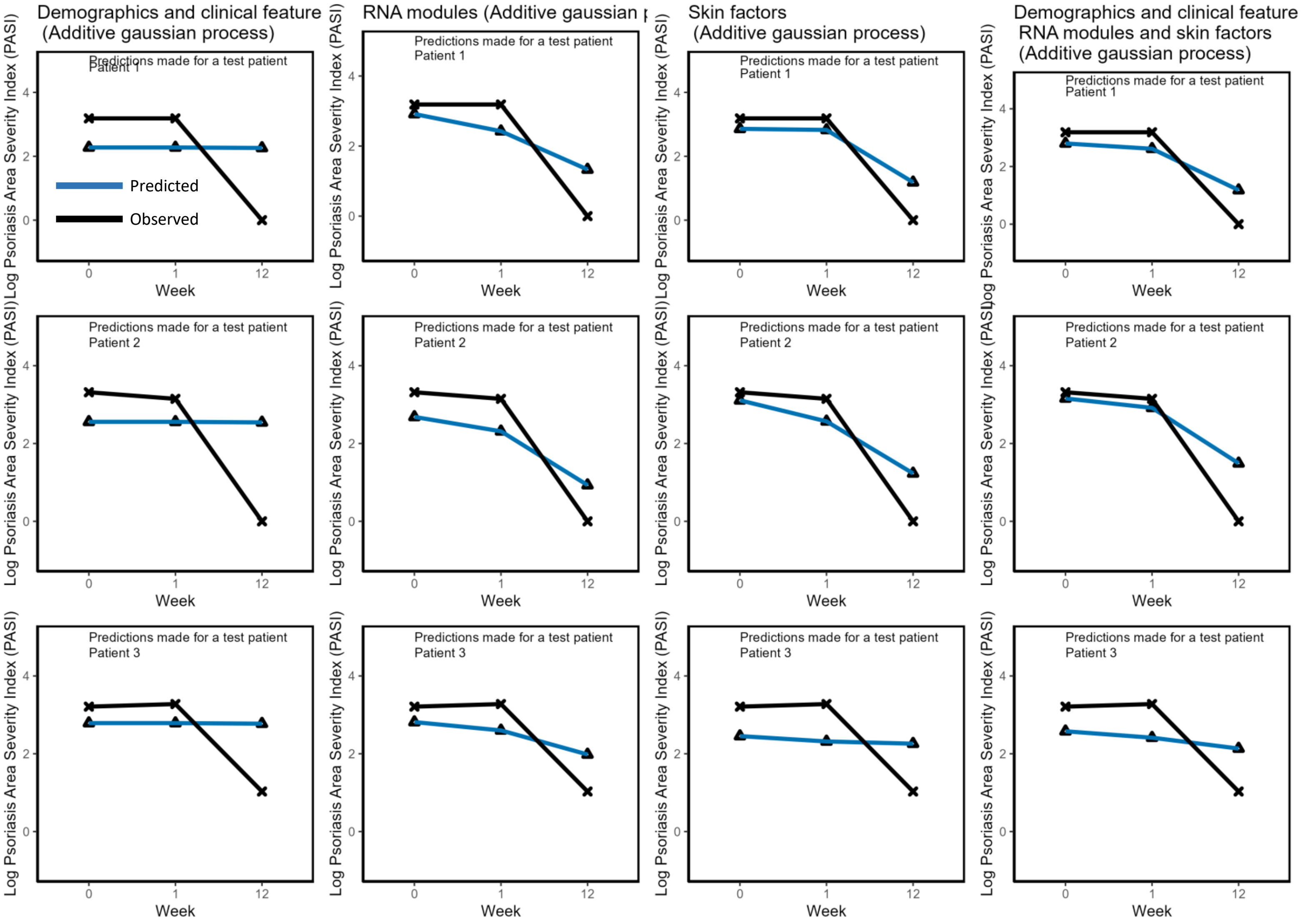


**Supplementary Figure 13. Predicted PASI trajectories across time based on gaussian process regression.** Patient trajectories predicted by the additive gaussian process regression models from a randomly shuffled 10-fold held-out testing dataset. The first regression model was trained on patient demographic and clinical information. The next two regression models were trained on module eigengenes or latent factors. The final regression model was trained on all features.

*Abbreviations:* PASI, psoriasis area and severity index.

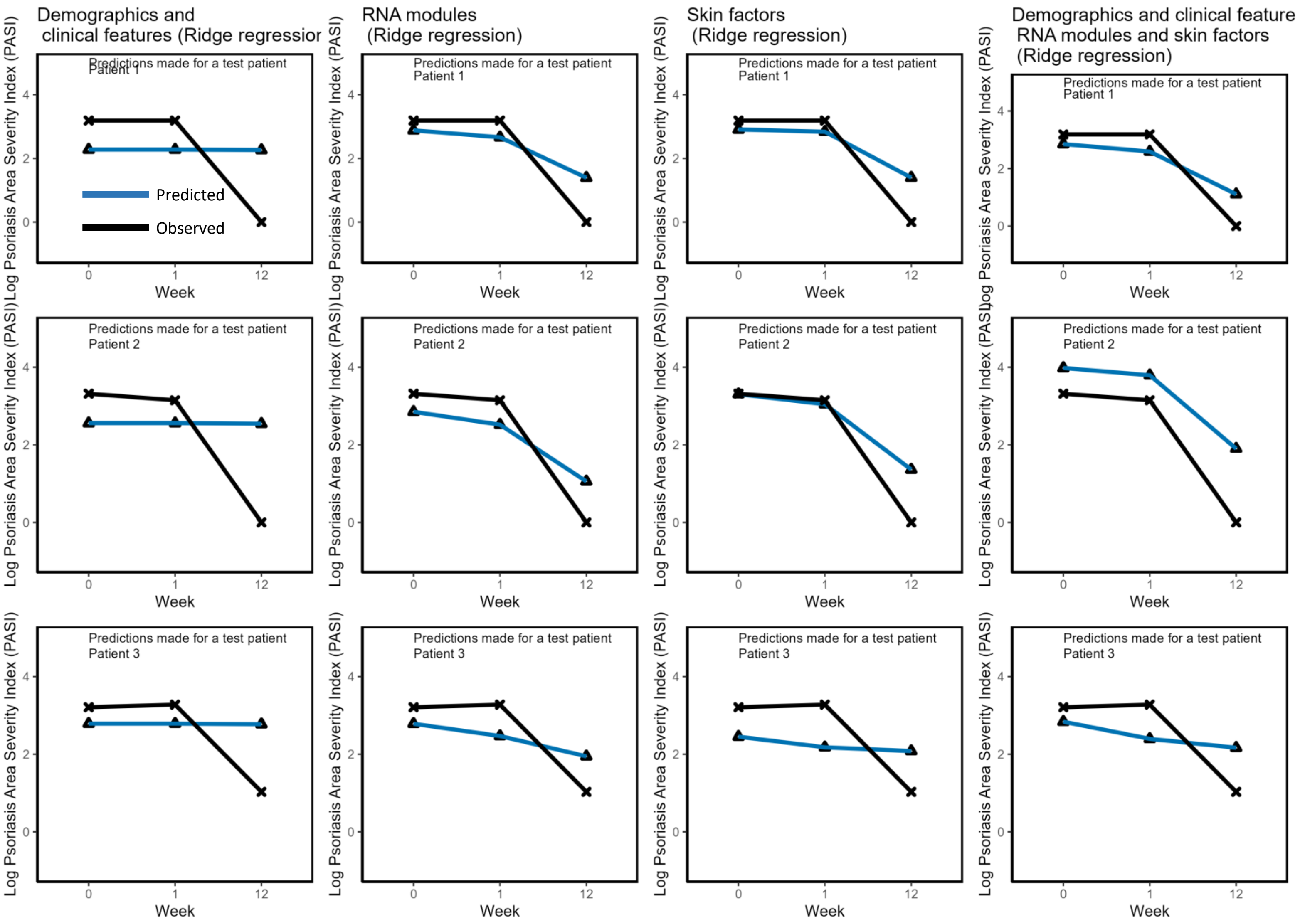


**Supplementary Figure 14. Predicted PASI trajectories across time based on linear ridge regression.** Patient trajectories predicted by the linear ridge regression models from a randomly shuffled 10-fold held-out testing dataset. The first regression model was trained on patient demographic and clinical information. The next two regression models were trained on module eigengenes or latent factors. The final regression model was trained on all features.

*Abbreviations:* PASI, psoriasis area and severity index.

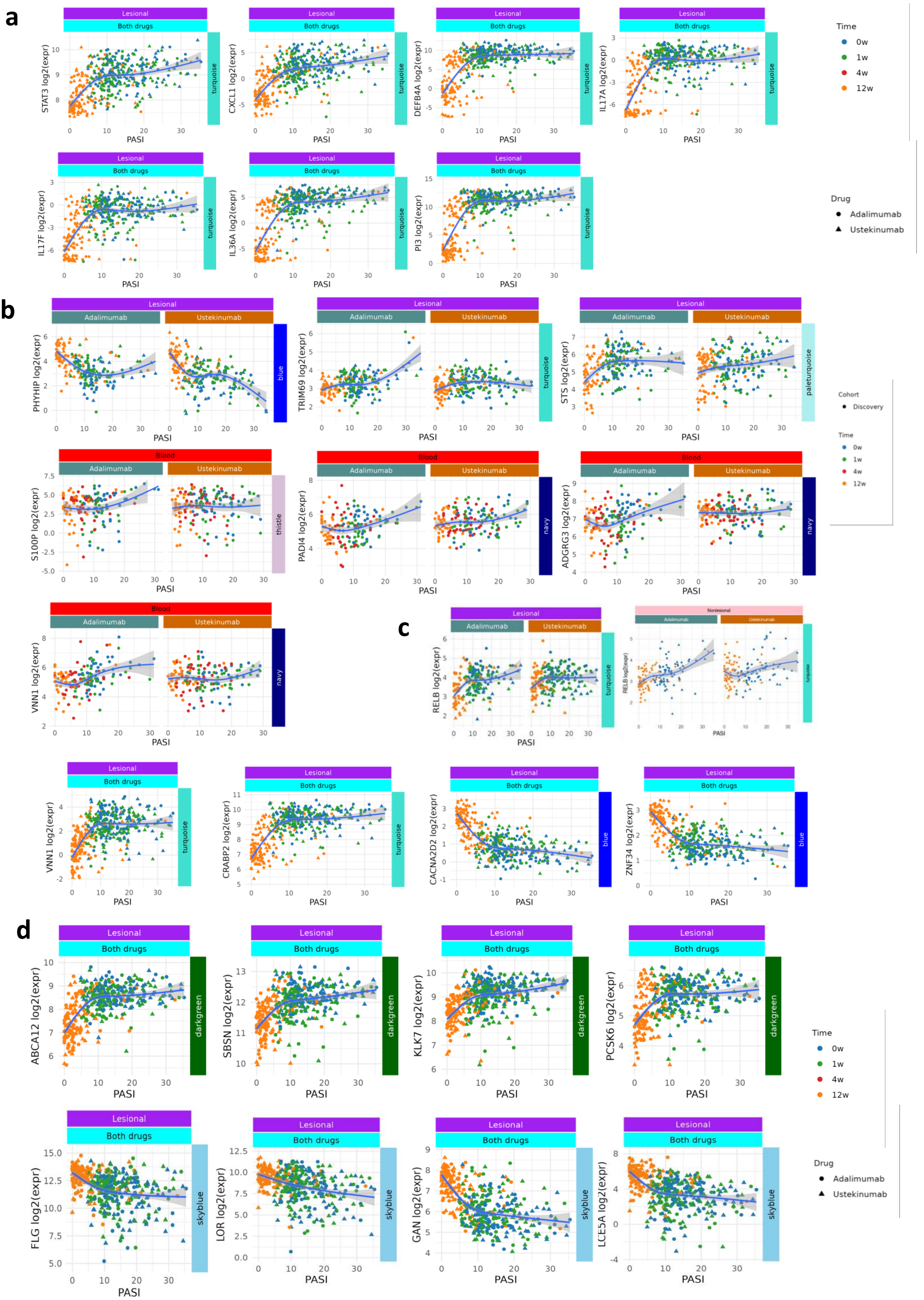
a
Lesional
Both drugs
STAT3 log2(expr)
CXCL1 log2(expr)
DEFB4A log2(expr)
IL17A log2(expr)
IL17F log2(expr)
IL36A log2(expr)
PI3 log2(expr)
PASI
turquoise
Time
0w
1w
4w
12w
Drug
Adalimumab
Ustekinumab
b
Lesional
Adalimumab
Ustekinumab
PHYHIP log2(expr)
TRIM69 log2(expr)
STS log2(expr)
blue
turquoise
paleturquoise
Cohort
Discovery
Blood
S100P log2(expr)
PADI4 log2(expr)
ADGRG3 log2(expr)
VNN1 log2(expr)
thistle
navy
c
RELB log2(expr)
Nonlesional
VNN1 log2(expr)
CRABP2 log2(expr)
CACNA2D2 log2(expr)
ZNF34 log2(expr)
d
ABCA12 log2(expr)
SBSN log2(expr)
KLK7 log2(expr)
PCSK6 log2(expr)
darkgreen
FLG log2(expr)
LOR log2(expr)
GAN log2(expr)
LCE5A log2(expr)
skyblue

**Supplementary Figure 15. Further examples of gene-level associations with disease severity in skin and blood.**

Scatter plots of the log2-CPM expression against PASI of further example disease severity-associated genes. Each point represents a single sample, is marked with a symbol according to drug cohort and colour-coded according to timepoint (blue = 0w, green = 1w, red = 4w, orange = 12w). The grey shading indicates 95% confidence intervals. The title banner provides information on the tissue and drug cohort. Module membership is denoted opposite the y axis. **(a)** Exemplars; **(b)** drug-specific examples; **(c)** PASI-associated signature genes, identified through SHAP analysis of blue and turquoise modules (Fig. 4D); **(d)** skin barrier genes that were also members of the darkgreen or skyblue modules in skin.

Curves represent natural-spline fits (3 d.f.) with 95 % confidence bands.

*Abbreviations:* CPM, counts per million; PASI, psoriasis area and severity index; w, week; SHAP, SHapley Additive exPlanations.

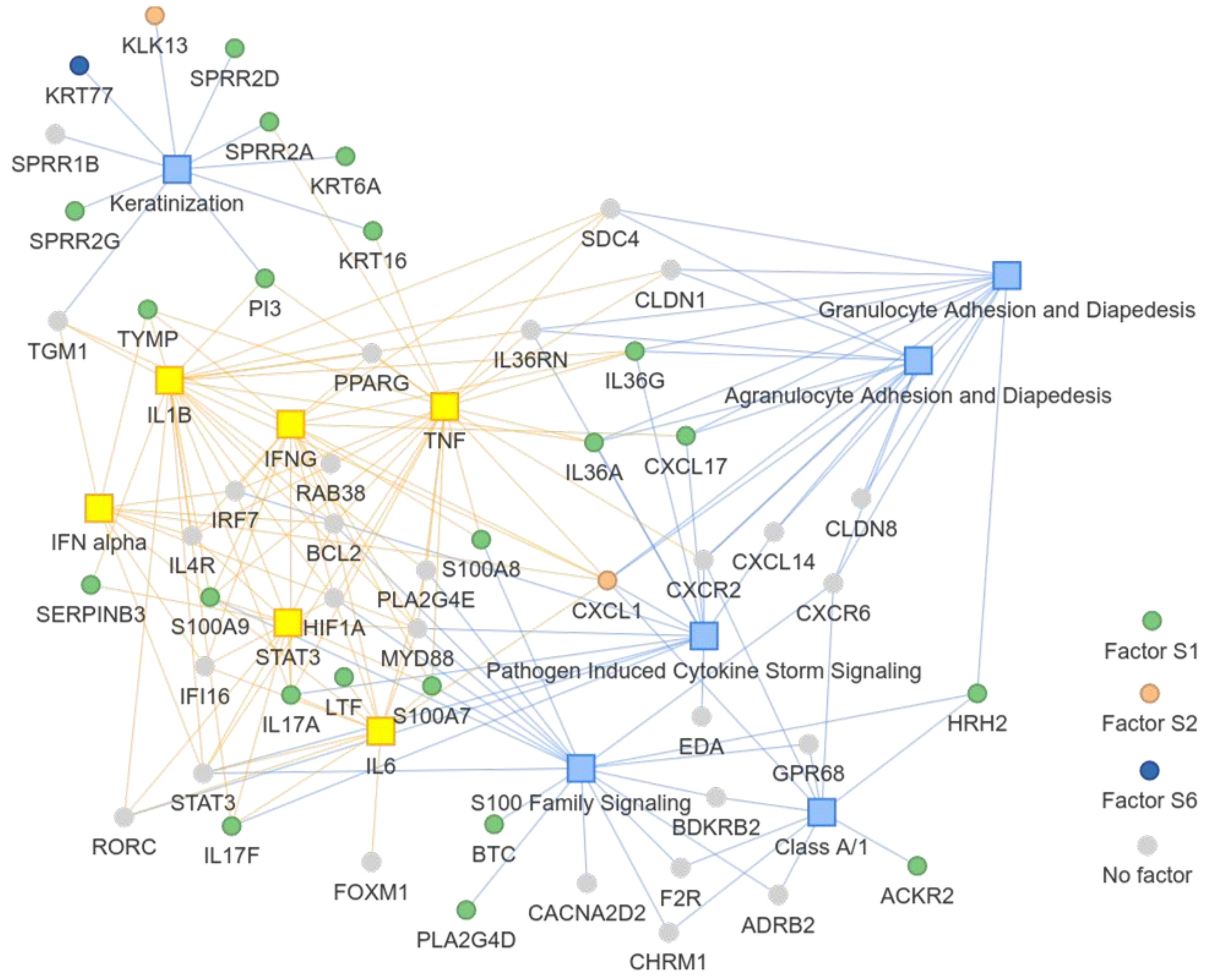


**Supplementary Figure 16. Network analysis of the top disease severity-associated genes in lesional skin for the combined drug cohort.**

Genes (circles) are coloured according to associated skin factors, upstream regulators (yellow squares) and canonical pathways (blue squares) were identified by IPA from all strongly disease severity-associated genes.

**a**

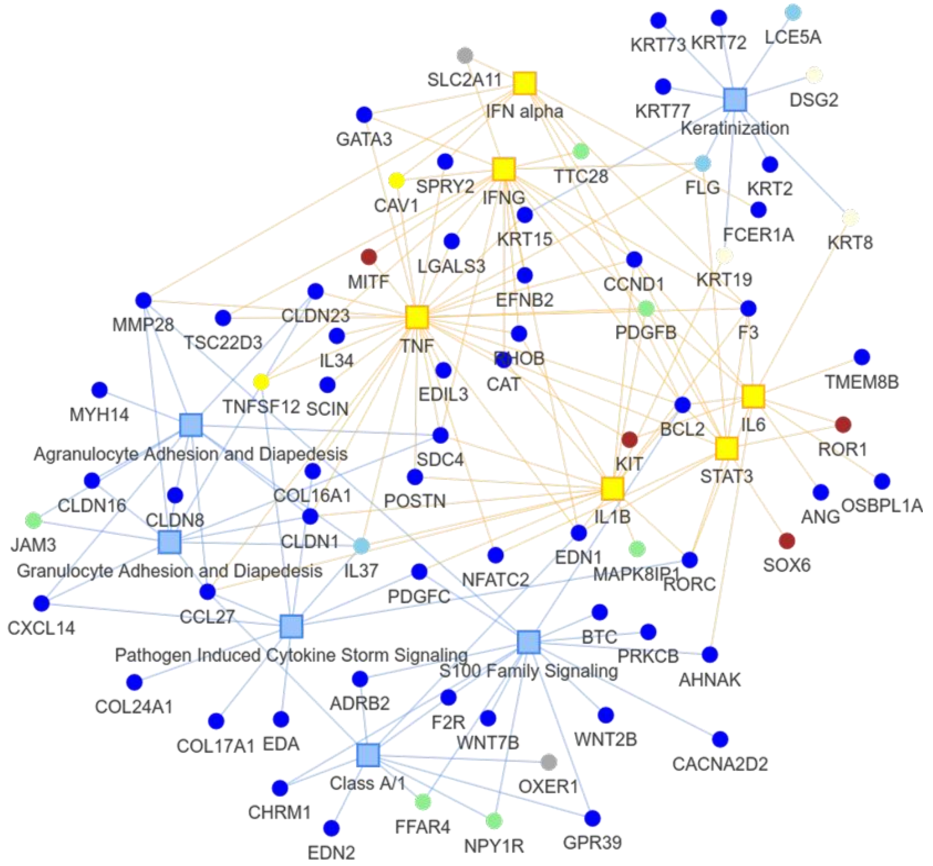


**b**

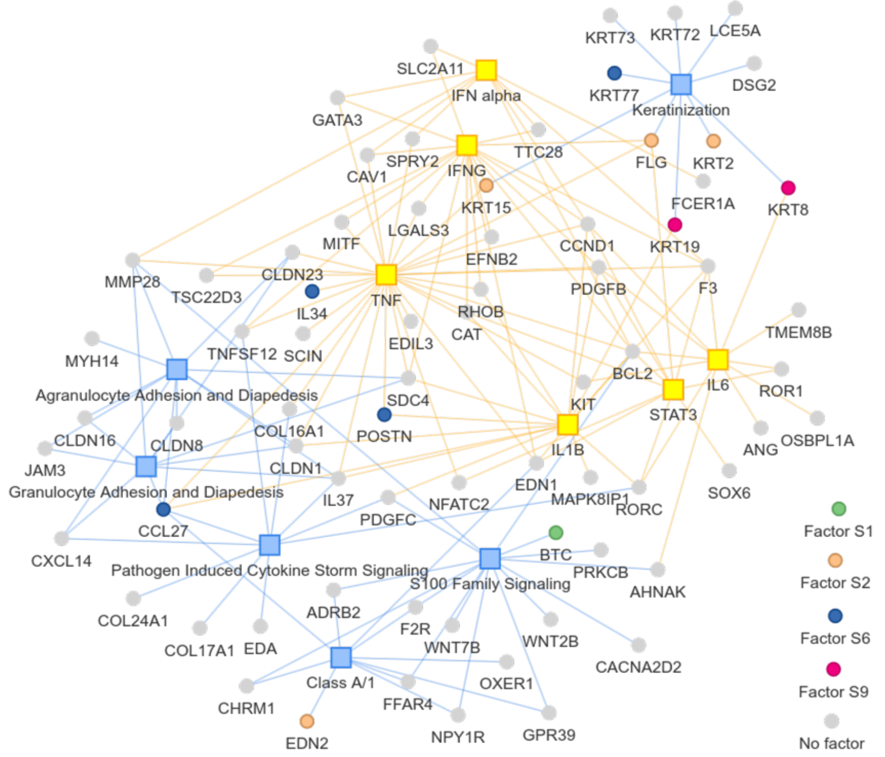


**Supplementary Figure 17. Network analysis of top negative disease severity-associated genes in lesional skin for the combined drug cohort.**

Networks are colour coded **(a)** according their WGCNA module, and **(b)** according to associated skin factors. Genes (circles) are coloured according to associated skin factors, upstream regulators (yellow squares) and canonical pathways (blue squares) were identified by IPA from all strongly disease severity-associated genes.

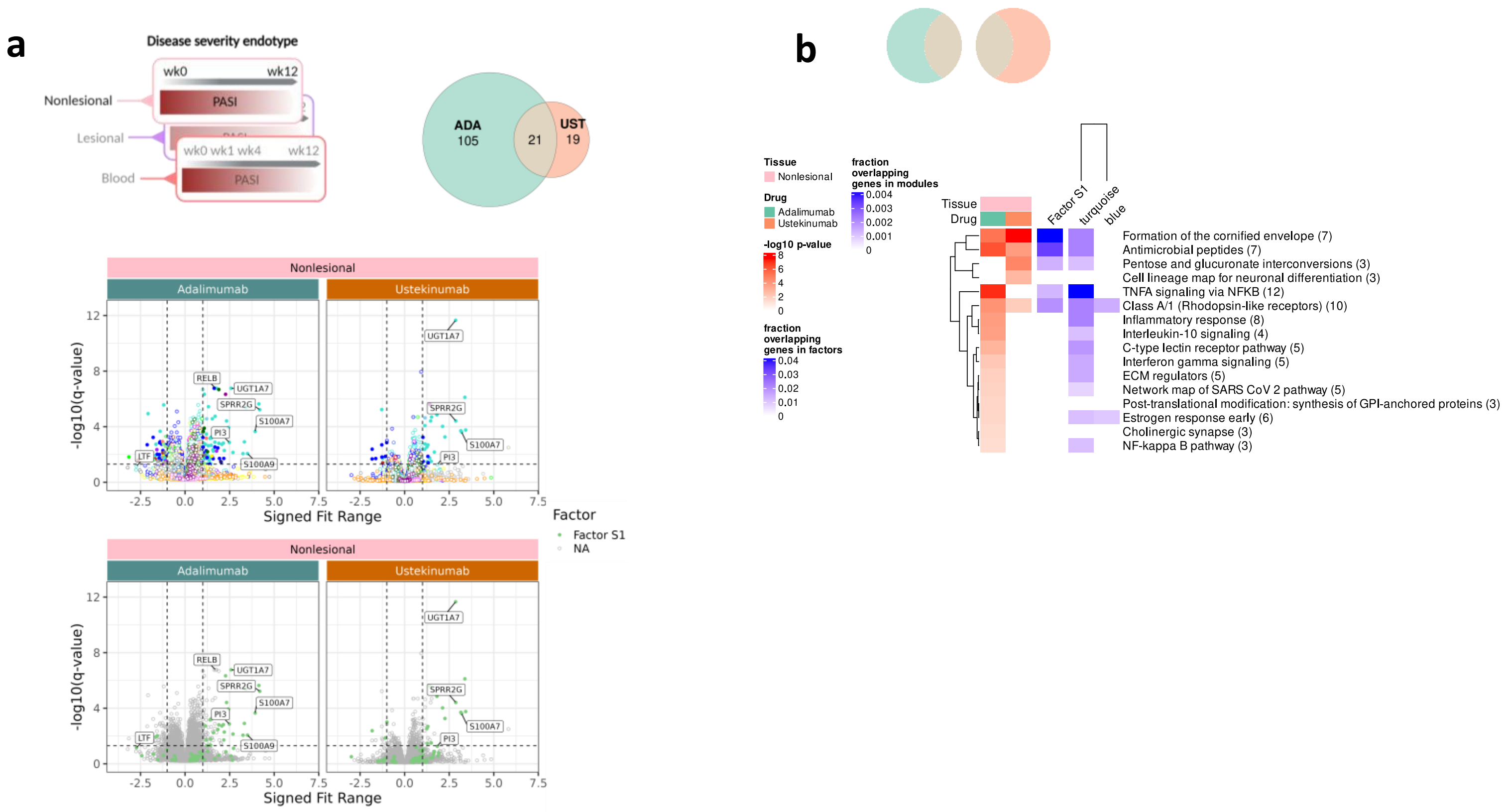


**Supplementary figure 18. Disease severity endotypes in non-lesional skin (NLS).**
**(a)** (Upper left) Schematic. (Upper right) Venn Diagram showing overlap of genes in NLS with significant (q-value<0.05) non-linear association (natural spline with 3 degrees of freedom) of expression with disease severity (DS), measured by PASI, in the adalimumab and ustekinumab cohorts. (Lower) Volcano plots of gene expression within the discovery and replication cohorts, facetted by drug cohort and plotted against signed fit range (overall direction and magnitude of DS association); points are coloured according to the WGCNA modules and ICA latent factors; example genes are labelled. **(b)** Metascape pathways from all co-expressed strongly DS-associated genes in each drug cohort, broken down by WGCNA module. The red heatmap shows the –log10 p-value, the blue shows the fraction of overlapping genes. The blue heatmap is normalised by the number of genes in the union of WGCNA module and Metascape pathway. *Abbreviations:* ADA, adalimumab; UST, ustekinumab; NLS, non-lesional skin; DS, disease severity; WGCNA, weighted gene correlation network analysis; ICA, independent component analysis.

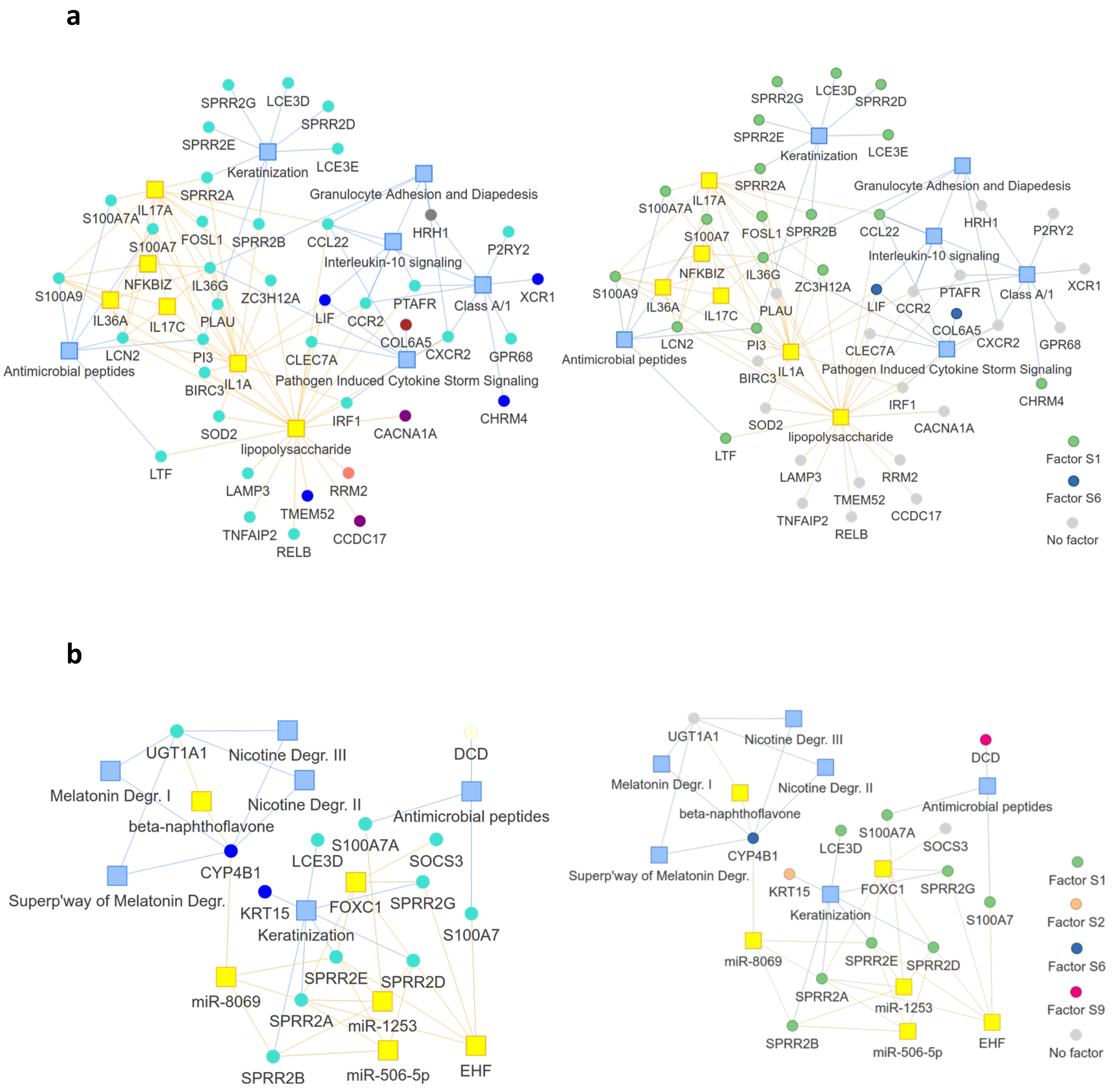


**Supplementary Figure 19. Network analysis of top negative disease severity-associated genes in non-lesional skin (NLS).**

Networks are shown for **(a)** the adalimumab cohort, colour coded according to their WGCNA module (left), and according to associated latent factors (right), and **(b)** the ustekinumab cohort, colour coded according to their WGCNA module (left), and according to associated skin factors (right). Significant genes (circles), upstream regulators (yellow squares) and canonical pathways (blue squares) were identified by IPA.

*Abbreviations:* WGCNA, weighted gene correlation network analysis; IPA, Ingenuity Pathway Analysis.

**a**

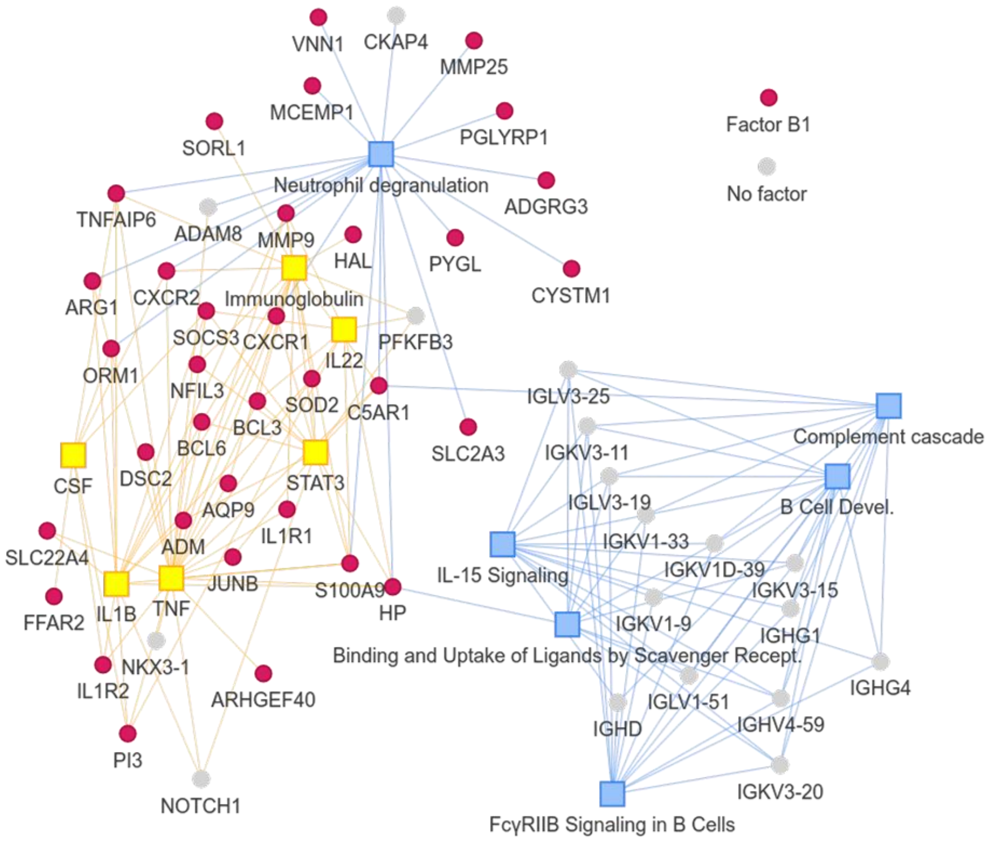


**b**

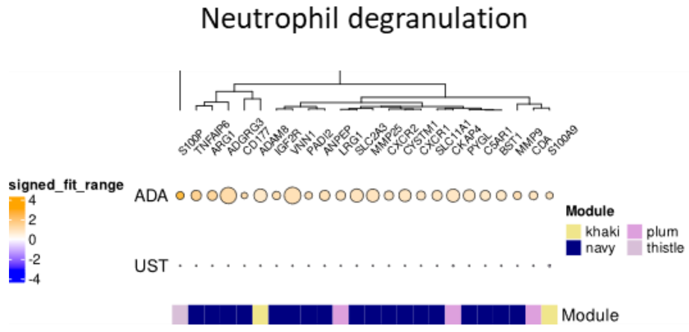


**Supplementary Figure 20. Functional analysis of the top disease severity-associated genes in blood for the adalimumab cohort.**

**(a)** Network analysis of the top disease severity-associated genes in blood for the adalimumab cohort. Genes (circles) are coloured according to associated latent factors, upstream regulators (yellow squares) and canonical pathways (blue squares) were identified by IPA from all strongly disease severity-associated genes. **(b)** Dot plots of the disease severity-associated genes within the Metascape neutrophil degranulation pathway. Gene expression is separated by drug cohort and clustered by signed fit range. The radius of the dot represents significance –log10 q-value and colour represents signed fit range. Module membership is recorded below each gene.

*Abbreviations:* IPA, Ingenuity Pathway Analysis.

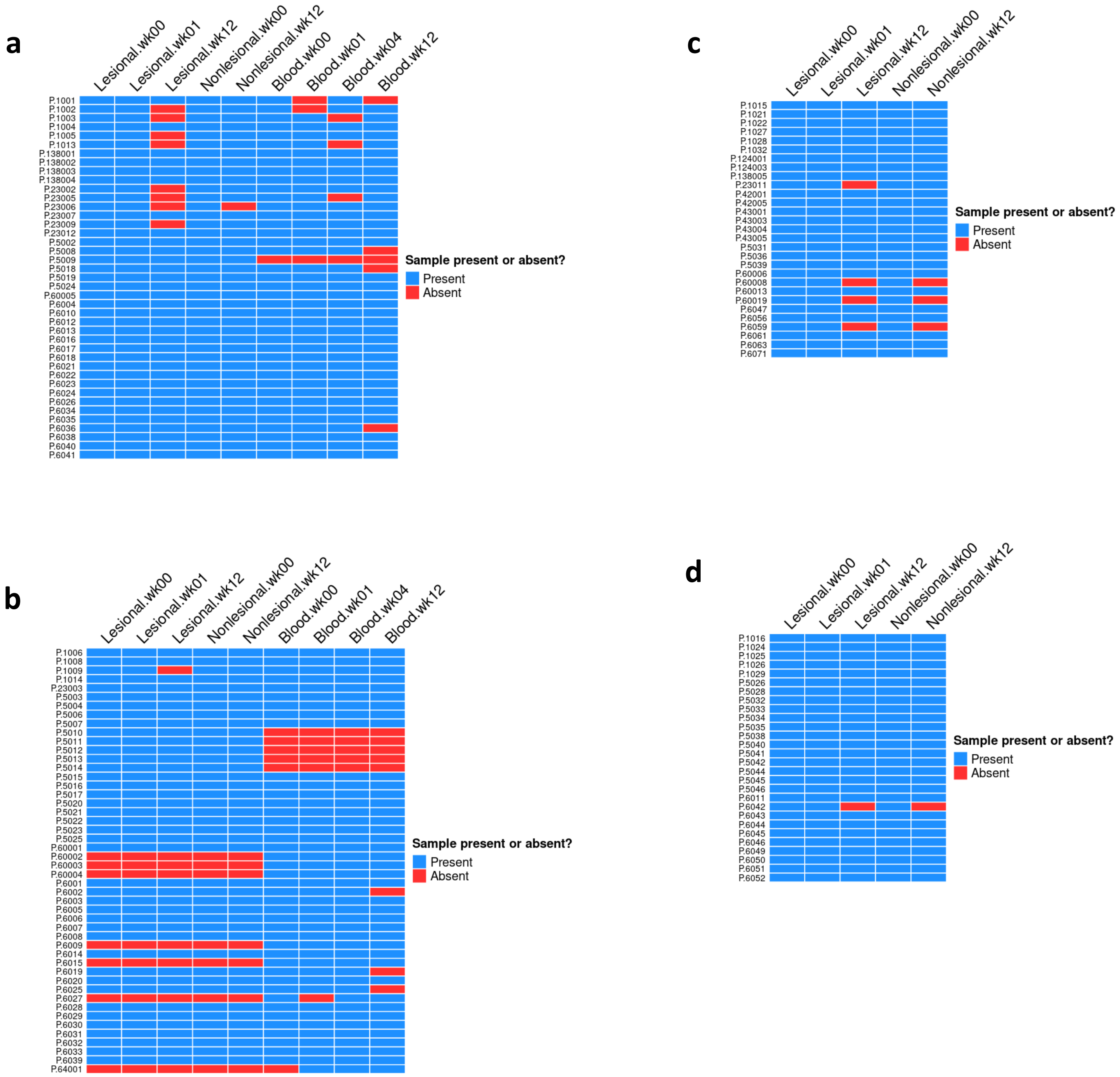


**Supplementary figure 21. Graphical representations of RNA-Seq sample completeness.**
Graphs are shown for the **(a)** adalimumab group and **(b)** ustekinumab group in the discovery cohort, and the **(c)** adalimumab group and **(d)** ustekinumab group in the replication cohort. In each graph, patients are displayed on the x-axis and tissue-times are displayed on the y-axis. Blue indicates that a sample was collected, sequenced and used for analysis at the corresponding tissue-time for the corresponding patient; red indicates a missing sample for the corresponding tissue-time and patient.

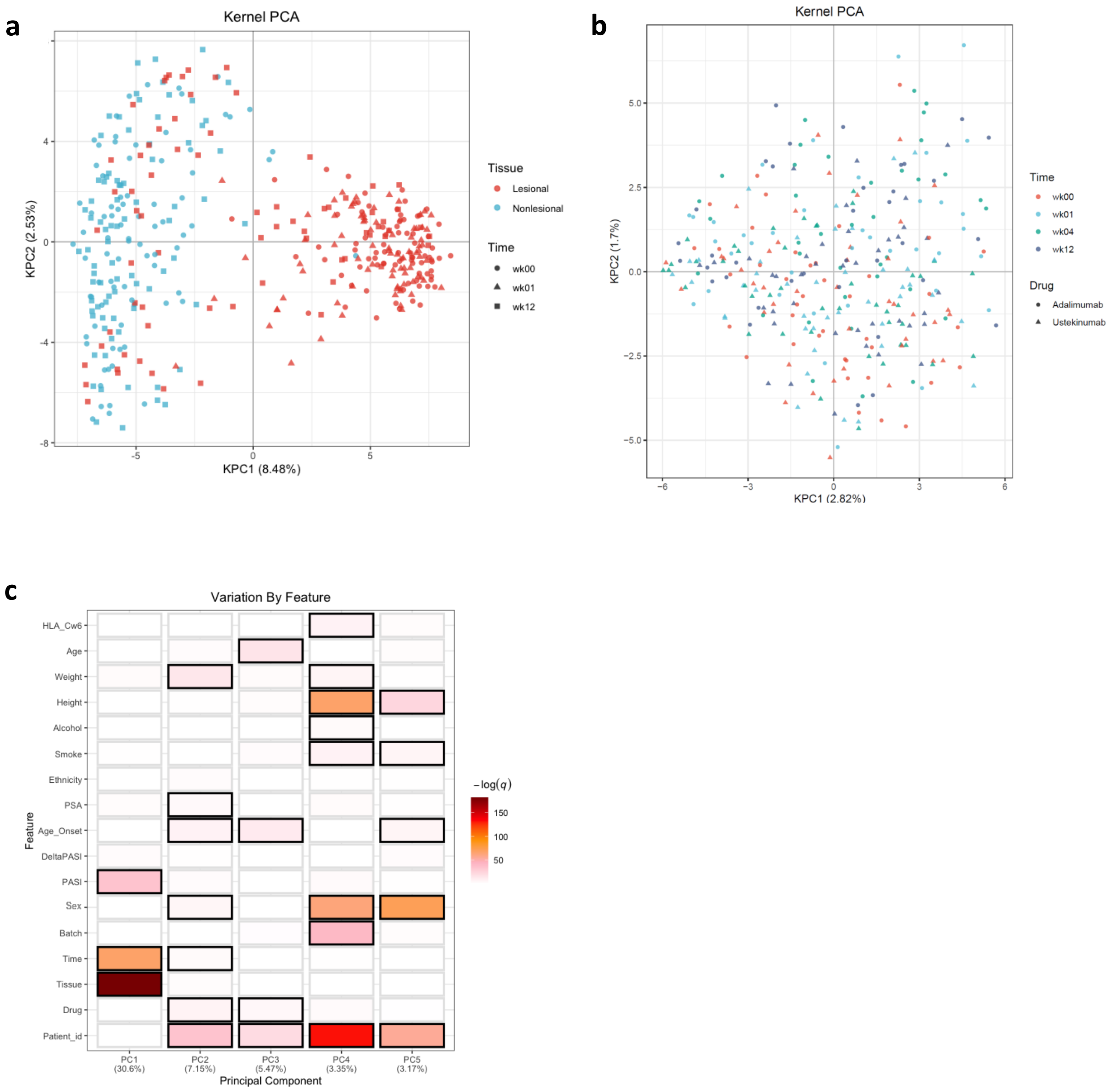


**Supplementary Figure 22. Drivers of transcriptomic variation in skin and blood.**
As an exploratory analysis, prior to conducting gene-level differential expression analysis or dimensionality reduction with WGCNA or ICA, we used Kernal PCA to explore the effect of tissue type, time point and drug on transcriptomic variation in the discovery cohort skin and blood RNA-Seq data. PCA plots of the first two principal components in **(a)** skin and **(b)** blood are shown. We used statistical analysis to investigate associations between the first five principal components in skin and an extended set of clinical and demographic traits; this is visualised in **(c)**.
*Abbreviations:* WGCNA, weighted gene correlation network analysis; ICA, independent component analysis; PCA, principal component analysis.

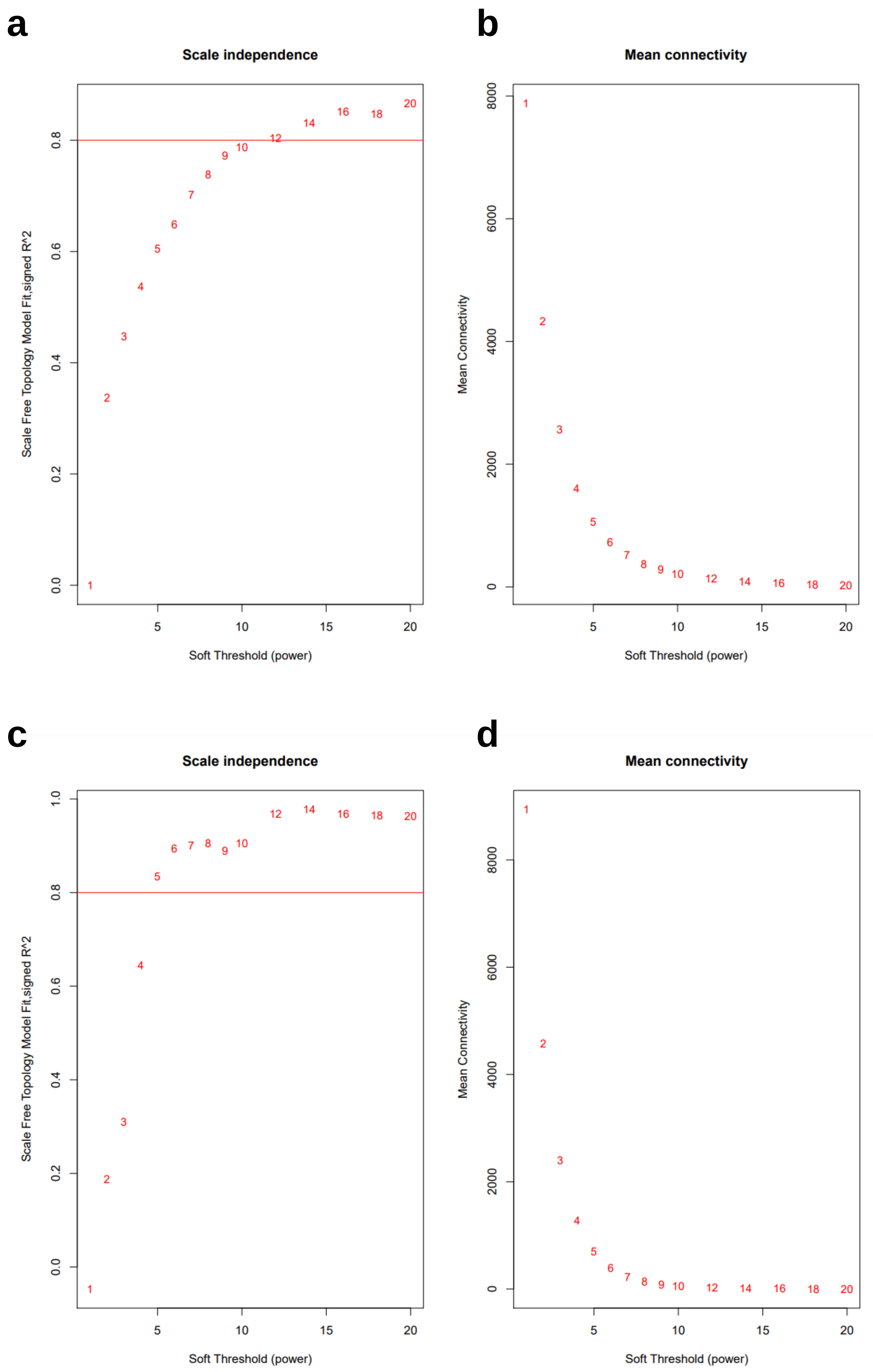
a
Scale independence
Scale Free Topology Model Fit,signed R^2
Soft Threshold (power)
b
Mean connectivity
Mean Connectivity
Soft Threshold (power)
c
Scale independence
Scale Free Topology Model Fit,signed R^2
Soft Threshold (power)
d
Mean connectivity
Mean Connectivity
Soft Threshold (power)

**Supplementary Figure 23. Identifying appropriate soft-thresholding powers for WGCNA in skin and blood.**

WGCNA involves creation of a gene-gene correlation matrix; prior to hierarchical clustering of this matrix, the correlation estimates are raised to a power in order to amplify the differences between high and low correlations; this power is chosen based on scale independence and mean connectivity. The scale independence **(a, c)** and mean connectivity **(b, d)** of the candidate soft-thresholding powers in skin **(a, b)** and blood **(c, d)**. The smallest power that passed an R2 threshold of 0.8 was selected for the downstream module identification steps. In skin this was 12 and in blood this was 5. Further details are available in the supplementary methods.

### Supplementary Table 1a. Demographic information of study recruits for PSORT-D

| | | Overall n=89 | Adalimumab n=41 | Ustekinumab n=48 |
|---|---|---|---|---|
| Biologic Naive, n (%) | No | 27 (30.3) | 10 (24.4) | 17 (35.4) |
| | Yes | 62 (69.7) | 31 (75.6) | 31 (64.6) |
| Ethnicity, n (%) | Asian or Asian British | 6 (6.7) | 3 (7.3) | 3 (6.2) |
| | Black or Black British | 2 (2.2) | 1 (2.4) | 1 (2.1) |
| | White | 80 (89.9) | 37 (90.2) | 43 (89.6) |
| | Other | 1 (1.1) | | 1 (2.1) |
| Sex, n (%) | F | 33 (37.1) | 15 (36.6) | 18 (37.5) |
| | M | 56 (62.9) | 26 (63.4) | 30 (62.5) |
| PsA, n (%) | Negative | 62 (69.7) | 27 (65.9) | 35 (72.9) |
| | Positive | 27 (30.3) | 14 (34.1) | 13 (27.1) |
| Flexural, n (%) | No | 53 (59.6) | 23 (56.1) | 30 (62.5) |
| | Not Done | 4 (4.5) | 2 (4.9) | 2 (4.2) |
| | Yes | 32 (36.0) | 16 (39.0) | 16 (33.3) |
| Scalp, n (%) | No | 17 (19.1) | 9 (22.0) | 8 (16.7) |
| | Not Done | 4 (4.5) | 2 (4.9) | 2 (4.2) |
| | Yes | 68 (76.4) | 30 (73.2) | 38 (79.2) |
| Palms, n (%) | No | 66 (74.2) | 33 (80.5) | 33 (68.8) |
| | Not Done | 4 (4.5) | 2 (4.9) | 2 (4.2) |
| | Yes | 19 (21.3) | 6 (14.6) | 13 (27.1) |
| Soles, n (%) | No | 67 (75.3) | 32 (78.0) | 35 (72.9) |
| | Not Done | 4 (4.5) | 2 (4.9) | 2 (4.2) |
| | Yes | 18 (20.2) | 7 (17.1) | 11 (22.9) |
| Nails, n (%) | No | 25 (28.1) | 13 (31.7) | 12 (25.0) |
| | Not Done | 4 (4.5) | 2 (4.9) | 2 (4.2) |
| | Yes | 60 (67.4) | 26 (63.4) | 34 (70.8) |
| Age, mean (SD) | | 44.4 (12.3) | 41.4 (10.3) | 46.9 (13.4) |
| Age of onset, mean (SD) | | 22.6 (13.2) | 21.2 (10.8) | 23.9 (15.1) |
| wk00 PASI, mean (SD) | | 15.3 (5.9) | 14.5 (5.2) | 15.9 (6.4) |
| wk01 PASI, mean (SD) | | 13.7 (6.0) | 12.4 (5.0) | 14.8 (6.6) |
| wk04 PASI, mean (SD) | | 9.1 (5.7) | 8.3 (5.2) | 9.8 (6.0) |
| wk12 PASI, mean (SD) | | 3.9 (4.9) | 4.1 (6.1) | 3.8 (3.8) |
| wk00 DLQI, mean (SD) | | 17.5 (7.0) | 16.9 (6.9) | 17.9 (7.1) |
| wk01 DLQI, mean (SD) | | 15.3 (7.2) | 14.3 (6.5) | 16.2 (7.7) |
| wk04 DLQI, mean (SD) | | 9.7 (6.7) | 7.9 (5.6) | 11.2 (7.2) |
| wk12 DLQI, mean (SD) | | 4.2 (5.2) | 3.6 (4.5) | 4.6 (5.7) |
| Cw6, n (%) | Negative | 46 (56.1) | 23 (56.1) | 23 (56.1) |
| | Positive | 36 (43.9) | 18 (43.9) | 18 (43.9) |
| DeltaPASI, mean (SD) | | 0.7 (0.2) | 0.8 (0.2) | 0.7 (0.2) |
| Smoking, n (%) | No | 27 (30.3) | 12 (29.3) | 15 (31.2) |
| | Yes | 62 (69.7) | 29 (70.7) | 33 (68.8) |
| Drink Alcohol, n (%) | No | 24 (27.0) | 4 (9.8) | 20 (41.7) |
| | Yes | 65 (73.0) | 37 (90.2) | 28 (58.3) |
| Alcohol (units/week), mean (SD) | | 8.3 (16.7) | 8.5 (10.2) | 8.1 (20.8) |
| BMI, mean (SD) | | 30.8 (6.4) | 31.6 (7.0) | 30.1 (5.7) |
| BMI status, n (%) | Healthy weight (>=18.5 <30) | 17 (19.3) | 8 (19.5) | 9 (19.1) |
| | Obese (>=30) | 44 (50.0) | 23 (56.1) | 21 (44.7) |
| | Overweight (>=25 <30) | 27 (30.7) | 10 (24.4) | 17 (36.2) |

**Supplementary Table 1b. Demographic information of study recruits for PSORT-R**

| | | Overall n=57 | Adalimumab n=29 | Ustekinumab n=28 |
|---|---|---|---|---|
| Biologic Naive, n (%) | No | 8 (14.0) | 1 (3.4) | 7 (25.0) |
| | Yes | 49 (86.0) | 28 (96.6) | 21 (75.0) |
| Ethnicity, n (%) | Asian or Asian British | 2 (3.5) | 1 (3.4) | 1 (3.6) |
| | Other | 2 (3.5) | 2 (6.9) | |
| | White | 52 (91.2) | 26 (89.7) | 26 (92.9) |
| | Black or Black British | 1 (1.8) | | 1 (3.6) |
| Sex, n (%) | F | 28 (49.1) | 13 (44.8) | 15 (53.6) |
| | M | 29 (50.9) | 16 (55.2) | 13 (46.4) |
| PsA, n (%) | Negative | 39 (68.4) | 16 (55.2) | 23 (82.1) |
| | Positive | 18 (31.6) | 13 (44.8) | 5 (17.9) |
| Flexural, n (%) | No | 24 (42.1) | 14 (48.3) | 10 (35.7) |
| | Yes | 33 (57.9) | 15 (51.7) | 18 (64.3) |
| Scalp, n (%) | No | 7 (12.3) | 5 (17.2) | 2 (7.1) |
| | Yes | 50 (87.7) | 24 (82.8) | 26 (92.9) |
| Palms, n (%) | No | 42 (73.7) | 20 (69.0) | 22 (78.6) |
| | Yes | 15 (26.3) | 9 (31.0) | 6 (21.4) |
| Soles, n (%) | No | 43 (75.4) | 20 (69.0) | 23 (82.1) |
| | Not Done | 1 (1.8) | 1 (3.4) | |
| | Yes | 13 (22.8) | 8 (27.6) | 5 (17.9) |
| Nails, n (%) | No | 24 (42.1) | 11 (37.9) | 13 (46.4) |
| | Yes | 33 (57.9) | 18 (62.1) | 15 (53.6) |
| Age, mean (SD) | | 46.2 (12.5) | 44.3 (11.6) | 48.2 (13.4) |
| Age of onset, mean (SD) | | 24.1 (15.4) | 22.0 (13.5) | 26.1 (17.1) |
| wk00 PASI, mean (SD) | | 16.2 (7.6) | 16.2 (7.2) | 16.3 (8.2) |
| wk01 PASI, mean (SD) | | 14.4 (6.9) | 14.0 (6.8) | 14.7 (7.1) |
| wk04 PASI, mean (SD) | | 9.1 (5.1) | 8.6 (4.7) | 9.6 (5.5) |
| wk12 PASI, mean (SD) | | 3.9 (4.4) | 4.2 (4.8) | 3.7 (4.0) |
| wk00 DLQI, mean (SD) | | 17.0 (6.1) | 17.9 (5.4) | 16.1 (6.8) |
| wk01 DLQI, mean (SD) | | 14.3 (6.7) | 14.1 (6.4) | 14.5 (7.1) |
| wk04 DLQI, mean (SD) | | 8.8 (6.7) | 8.0 (5.4) | 9.6 (7.8) |
| wk12 DLQI, mean (SD) | | 3.4 (4.2) | 4.6 (5.2) | 2.1 (2.1) |
| Cw6, n (%) | Negative | 27 (48.2) | 12 (41.4) | 15 (55.6) |
| | Positive | 29 (51.8) | 17 (58.6) | 12 (44.4) |
| DeltaPASI, mean (SD) | | 0.7 (0.3) | 0.7 (0.3) | 0.7 (0.3) |
| Smoking, n (%) | No | 21 (36.8) | 12 (41.4) | 9 (32.1) |
| | Yes | 36 (63.2) | 17 (58.6) | 19 (67.9) |
| Drink Alcohol, n (%) | No | 21 (36.8) | 16 (55.2) | 5 (17.9) |
| | Yes | 36 (63.2) | 13 (44.8) | 23 (82.1) |
| Alcohol (units/week), mean (SD) | | 7.1 (10.6) | 4.3 (7.9) | 10.1 (12.2) |
| BMI, mean (SD) | | 32.2 (8.5) | 33.7 (8.7) | 30.7 (8.2) |
| BMI status, n (%) | Healthy weight (>=18.5 <30) | 13 (22.8) | 4 (13.8) | 9 (32.1) |
| | Obese (>=30) | 29 (50.9) | 17 (58.6) | 12 (42.9) |
| | Overwight (>=25<30) | 15 (26.3) | 8 (27.6) | 7 (25.0) |

**Supplementary Table 6: PASI prediction performance of Ridge models. Shown are means ± SD for 20 randomly shuffled 10-fold testing datasets.**

| **Inputs** | **Test MAE** | **Test $R^2$** |
|---|---|---|
| Demographics/clinical features<br>WGCNA modules<br>ICA factors | 0.47 ± 0.10 | 0.50 ± 0.10 |
| All WGCNA modules | 0.45 ± 0.06 | 0.54 ± 0.09 |
| All ICA factors | 0.46 ± 0.10 | 0.50 ± 0.11 |

**Supplementary Table 7: Performance metrics for modules at the gene-level using additive Gaussian process models. Mean performance metrics ± SD for testing datasets.**

| Model inputs | Test MAE | Test $R^2$ |
|---|---|---|
| Top 10 Factor S1 genes | $0.47 \pm 0.06$ | $0.40 \pm 0.10$ |
| Top 10 Factor S2 genes | $0.48 \pm 0.06$ | $0.37 \pm 0.15$ |
| Top 10 Factor S9 genes | $0.54 \pm 0.08$ | $0.21 \pm 0.16$ |
| Top 10 Factor S6 genes | $0.51 \pm 0.08$ | $0.26 \pm 0.14$ |
| Top 10 Factor S8 genes | $0.62 \pm 0.09$ | $0.04 \pm 0.09$ |
| Top 10 Factor S21 genes | $0.64 \pm 0.08$ | $-0.02 \pm 0.05$ |

**Supplementary Table 8: Performance metrics for modules at the gene-level using additive Gaussian process models. Mean performance metrics ± SD for testing datasets.**

| Model inputs | Test MAE | Test $R^2$ |
|---|---|---|
| Top 10 Turquoise genes | $0.43 \pm 0.06$ | $0.47 \pm 0.13$ |
| Top 10 Violet genes | $0.52 \pm 0.07$ | $0.29 \pm 0.13$ |
| Top 10 Steelblue genes | $0.57 \pm 0.07$ | $0.16 \pm 0.14$ |
| Top 10 Blue genes | $0.47 \pm 0.07$ | $0.44 \pm 0.12$ |
| Top 10 Tan genes | $0.50 \pm 0.06$ | $0.33 \pm 0.09$ |

**Supplementary Table 9: Correlations between factors and baseline PASI**

| Factor | Correlation | P-value | Adjusted p-value |
|---|---|---|---|
| **S18** | -0.278 | 0.00092 | 0.01286 |
| **S2** | 0.275 | 0.00107 | 0.01286 |
| **S1** | 0.245 | 0.00366 | 0.02925 |
| **S17** | 0.228 | 0.00685 | 0.04110 |
| **S10** | -0.170 | 0.04587 | 0.22020 |
| **S23** | -0.149 | 0.07913 | 0.31585 |
| **S16** | 0.143 | 0.09212 | 0.31585 |
| **S21** | 0.125 | 0.14280 | 0.42841 |
| **S12** | 0.109 | 0.20286 | 0.51030 |
| **S13** | 0.097 | 0.25614 | 0.51030 |
| **S9** | -0.096 | 0.26254 | 0.51030 |
| **S24** | -0.093 | 0.27603 | 0.51030 |
| **S4** | -0.090 | 0.29293 | 0.51030 |
| **S14** | -0.089 | 0.29768 | 0.51030 |
| **S3** | -0.071 | 0.40563 | 0.63005 |
| **S8** | -0.069 | 0.42003 | 0.63005 |
| **S15** | 0.060 | 0.48076 | 0.67871 |
| **S20** | -0.056 | 0.51191 | 0.68255 |
| **S19** | 0.045 | 0.59873 | 0.74658 |
| **S5** | 0.042 | 0.62215 | 0.74658 |
| **S11** | -0.031 | 0.71812 | 0.82071 |
| **S6** | 0.021 | 0.80188 | 0.87478 |
| **S7** | 0.011 | 0.90199 | 0.94121 |
| **S22** | 0.004 | 0.96012 | 0.96012 |
